# Genetically-Encoded Discovery of Perfluoroaryl-Macrocycles that Bind to Albumin and Exhibit Extended Circulation *in-vivo*

**DOI:** 10.1101/2022.08.22.504611

**Authors:** Jeffrey Y.K. Wong, Steven E. Kirberger, Ryan Qiu, Arunika I. Ekanayake, Payam Kelich, Susmita Sarkar, Edgar R. Alvizo-Paez, Jiayuan Miao, Shiva Kalhor-Monfared, John J. Dwyer, John M. Nuss, Yu-Shan Lin, Matthew S. Macauley, Lela Vukovic, William C.K. Pomerantz, Ratmir Derda

**Affiliations:** Department of Chemistry, University of Alberta, Edmonton, AB T6G 2G2, Canada; Department of Chemistry, University of Minnesota, Minneapolis, MN 55455, USA; Department of Chemistry and Biochemistry, University of Texas at El Paso, El Paso, TX 79968, USA; Department of Chemistry, Tufts University, Medford, MA 02155, USA; Ferring Research Institute, San Diego, CA 92121, USA

## Abstract

In this paper, we report selection of albumin-binding macrocyclic peptides from genetically encoded libraries of peptides modified by perfluoroaryl-cysteine S_N_Ar chemistry. Modification of phage-displayed libraries SXCX*_n_*C-phage, *n*=3–5, where X is any amino acid except for cysteine by decafluoro-diphenylsulfone (**DFS**), yields genetically-encoded library of octafluoro-diphen-ylsulfone-crosslinked macrocycles (**OFS**-SXCX*_n_*C-phage). Selection from these libraries using albumin as a bait identified a family of significantly enriched perfluoroaryl-macrocycles. Synthesis of perfluoroaryl-macrocycles predicted by phage display and testing their binding properties by ^19^F NMR and fluorescent polarization identified **OFS**-macrocycle with SICRFFC sequence as the most potent albumin binder. We observed that **OFS**-macrocycles slowly react with biological nucleophiles such as glutathione. Replacing decafluoro-diphenylsulfone by nearly isosteric pentafluorophenyl sulfide yielded perfluorophenylsulfide (**PFS**)-crosslinked macrocycles devoid of undesired reactivity. The augmented lead **PFS**-macrocycle with SICRFFC sequence exhibited *K_D_* = 4–6 μM towards human serum albumin and similar affinities towards rat and mouse albumins. When injected in mouse, the **PFS**-SICRFFCGGG compound was significantly retained in circulation *in vivo* when compared to control **PFS**-macrocyclic peptide. The perfluoroaryl-macrocycles with SICRFFC motif are the smallest known peptide macrocycle with significant affinity for human albumin and they are a productive starting point for future development of compact macrocycles with predictable circulation half-life *in vivo*.

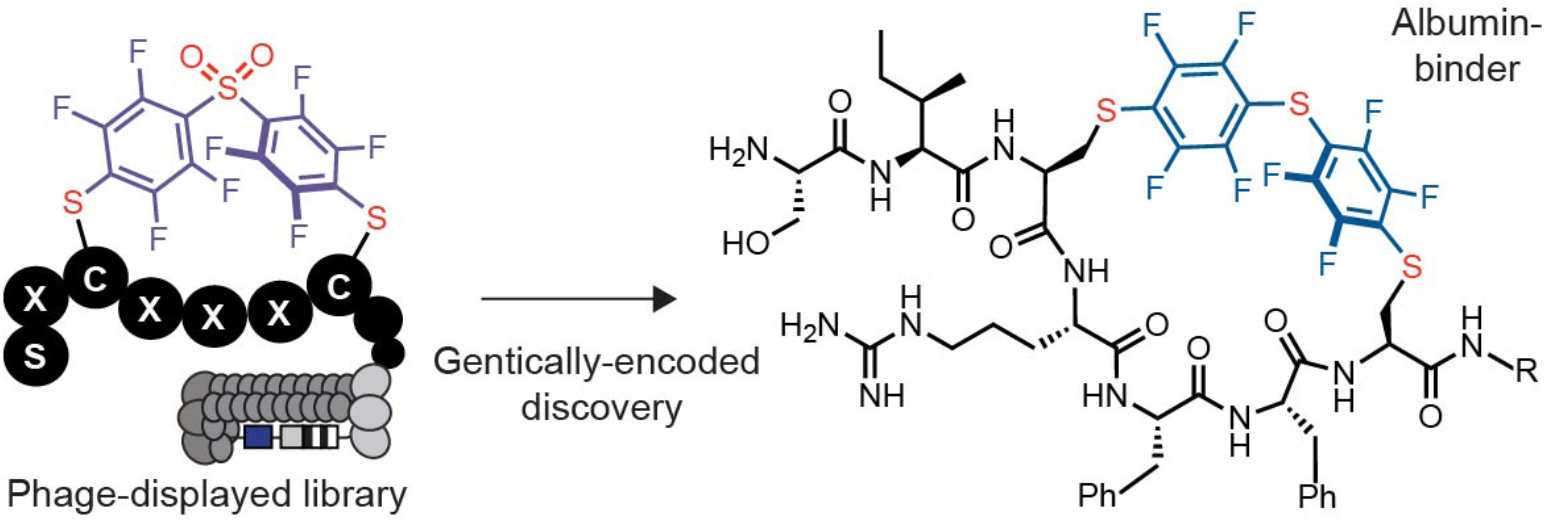

## Introduction

There are around 80 peptide drugs on the global market; more than 150 peptides are in clinical development and another 400–600 peptides undergoing preclinical studies.^1^ The large surface area of peptides, compared to a typical small molecule drugs allows peptides to interact with expanded binding interfaces commonly found in protein–protein interactions, protein–carbohydrate and protein–DNA interactions. These proteins, classified as “undruggable targets”, have been difficult to target using conventional small molecule therapeutic but many of them have been addressed by peptide, proteins, or antibody therapeutics. Peptides are the smallest among the latter three modalities—2 kDa to 10 kDa for peptides versus 150 kDa for full-sized antibodies—and they possess distinct pharmacokinetic (PK) properties. For example, bio-distribution of peptides and small proteins inside tumors and other non-vascularized tissues is improved when compared to full-size antibodies. Several clinical candidates (TH1902, TH1904, BT5528, BT8009, BT1718, MMP-14) capitalize on such improved biodistribution.^2–4^ Unlike antibodies, which remain in circulation for 1–3 weeks due to the association with the neonatal Fc-receptor (FcRn) on the surface of immune cells,^5^ peptide therapeutics clear within minutes to hours from plasma by renal filtration. Fast clearance is a beneficial property in several therapeutic applications, such as imaging (e.g., “tumor paint”), radionuclide delivery (e.g., Lurathera™),^6^ and in administration of short acting peptide hormones. For more widespread adaptation of peptide modalities in diverse therapeutic applications it is desired to tune the circulation life-time of peptides from minutes to hours.

Only few peptides exhibit a natural extended circulation lifetime: a therapeutically relevant example is a natural venom, 39-residue peptide ‘exendin 4’, with low renal clearance in humans (5–7 h).^7^ This peptide gave rise to FDA-approved drug exenatide for the treatment of type 2 diabetes.^8, 9^ Despite favourable circulation half-life, modified derivatives of exenatide—liraglutide, albiglutide, dulaglutide, lixisenatide and semaglutide^1^— have been developed to tune circulation half-life and other PK properties. The majority of peptides and small proteins have to be modified as well to increase their circulation time. Such modification could be divided into several classes: Class 1: increase in size via covalent linkage to polyethylene glycol (PEG),^10, 11^ polyglycerol^12^ and other synthetic macromolecules. Interestingly, steric hindrance by these size-increasing moieties also protects against proteolytic degradation.^13–15^; Class 2: increase in size via controlled oligomerization;^16^ Class 3: covalent linking to long-living serum protein (e.g., FDA-approved drugs albiglutide and dulaglutide, exanatide that conjugated to albumin and the IfG4 Fc domain)^17–19^ and Class 4: incorporation of moieties that bind non-covalently to serum proteins such as albumin,^20–23^ immunoglobulin,^24, 25^ FcRn,^26^ transthyretin,^27^ and transferrin.^28, 29^ An important example in the last class is lipidation of peptides to allow interaction with serum albumin. Lipidation has been one of the most successful strategies to prolong the half-life of peptides and small proteins such as insulin giving rise to FDA-approved drugs such as Levemir^®^, Tresiba^®^, Victoza^®^, Saxenda^®^, and Ozempic^®^ with extended serum half-life^30, 31^. The improved properties of these and many other drugs stemming from their association with albumin mandate investigation of albumin as carrier for therapeutic applications.

Albumin is the most abundant protein in plasma with an average concentration of 600 μM and has an average half-life of 19 days.^32^ The main mechanism leading to the long half-life of albumin and antibody are similar: both proteins interact with FcRn on the surface of immune cells. This binding results in transient endocytosis of these proteins, and as a result, they are frequently sequestered from circulation and protected from clearance. At physiological pH, the binding affinity between albumin and FcRn is low; however, the interaction under acidic conditions in the endosome is strong to avoid lysosomal degradations and recycling of albumin to the extracellular space.^5^ Albumin acts as a versatile carrier of essential fatty acids and diverse small organic molecules.^32^ Among all the long-circulating serum proteins, albumin is considered to be one of the most important targets because of its ability to interact with hydrophobic small molecule drugs and enhance their pharmacokinetic properties. Recurrent therapeutic success of rationally lipidated peptides and proteins^33^ fuels interest in rational development of small molecules as well as non-lipidated proteins and peptides that bind to albumin.

Many FDA-approved small molecule drugs have an intrinsic affinity for human serum albumin (HSA). Targeted development of small molecules with high affinity for HSA has been a topic of research over the last 15 years (see recent review^23^). Anti-HSA antibodies, nanobodies,^34^ DARPins,^35^ and other protein domains have been also developed. Such proteins can be fused to therapeutic proteins of interest to extend their *in vivo* circulation. Similarly short peptides that bind to HSA could be used in tandem with therapeutic peptide or protein sequences to dial in predictable half-life for such therapeutics. Such short albumin-binding peptides could empower development of many future therapeutic peptides because they could be built into *any* genetically encoded peptide library (e.g., displayed on phage, RNA and other platforms) to give rise to billion-scale libraries with predictable *in vivo* half-life. However, short HSA-binding peptides are scarce. A 31-mer peptide DX-236 (Ac-AEGTGDFWFCDRIAWYPQHLCEFLDPEGGGK-NH_2_) with a binding affinity of 1.9 μM was identified by Dyax Corp., and used to purify albumin (**Figure 1A**).^36^ A 21-mer peptide SA-21 (Ac-RLIEDICLPRWGCLWEDD-NH_2_) with a binding affinity of 467 nM to HSA was identified at Genentech (**Figure 1B**)^37^ and subsequently conjugated to ligands for urokinase-type plasminogen activator,^20, 38^ Fab antibody fragments^39, 40^ and small proteins^41^ to prolong their circulation half-lives. Heinis and co-workers developed a short heptapeptide modified by fluorescein isothiocyanide (FITC) and palmitic acid (FITC-EYEY-K_palm_ESE-NH_2_) with a binding affinity of 39 nM to HSA (**Figure 1C**)^21^ and the presence of both lipid moiety and fluorescein was critical for the binding of this peptide. This FITC-lipopeptide was fused to two different bicyclic peptides to boost the half-lives from minutes to hours.^21^ Success of DX-236, SA-21 and FITC-lipopeptide and other examples from the literature demonstrated the possibility of using HSA as a target for genetically-encoded selection to identify HSA-binding peptides with extended circulation half-life. There is a need for development of other albumin binding peptide modalities that have lower molecular weight.^21, 37^ Towards this goal, we employ genetically encoded phage-displayed libraries of chemically modified macrocycles to develop new classes of albumin binding mini scaffolds. To hone on shortest possible peptide sequences, we employed a phage-displayed libraries SXCX*_n_*C, *n*=3–5 modified with decafluorodiphenyl sulfone (**DFS**)^42, 43^ where X is any amino acid except for cysteine (**Figure 1D**). We hypothesized that a perfluoroaromatic linchpin would serve as useful pharmacophore recognized by one of the binding sites of HSA similarly to the binding of fatty acid in lipidated peptides.

**Figure 1:**
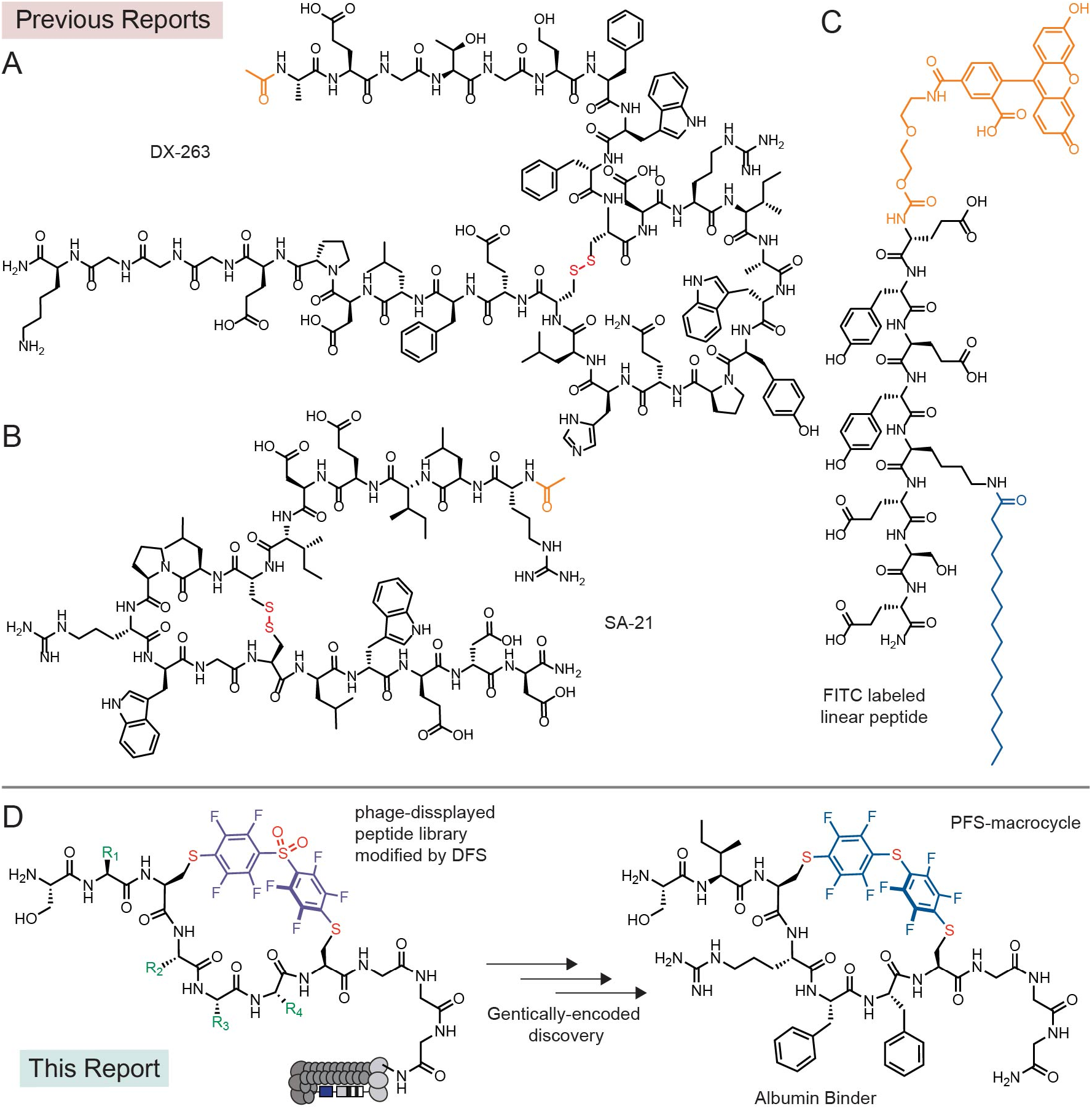
Discovery of albumin binding peptides. Previous reports of (A) macrocyclic peptide: DX-263,^36^ (B) macrocyclic peptide: SA-21,^37^ (C) a linear peptide: FITC-EYEYK_palm_ESE-NH_2_.^17^ (D) This report describes a chemically modified phage-displayed library for discovery of a small macrocyclic albumin binder.

## Results and discussion

### Selection of albumin binders

We devised and conducted three discovery campaigns that used different library architecture and selection strategies. In the first discovery campaign, we modified the phage libraries of structure SXCX_4–5_C with **DFS** following a previously published protocol and confirmed that 85% of the phage library is modified to yield octafluoro-diphenylsulfone-crosslinked macrocycles (**OFS**-SXCX_4–5_C-phage) (**Figure 2A**, **Figure S1A**).^42^

**Figure 2:**
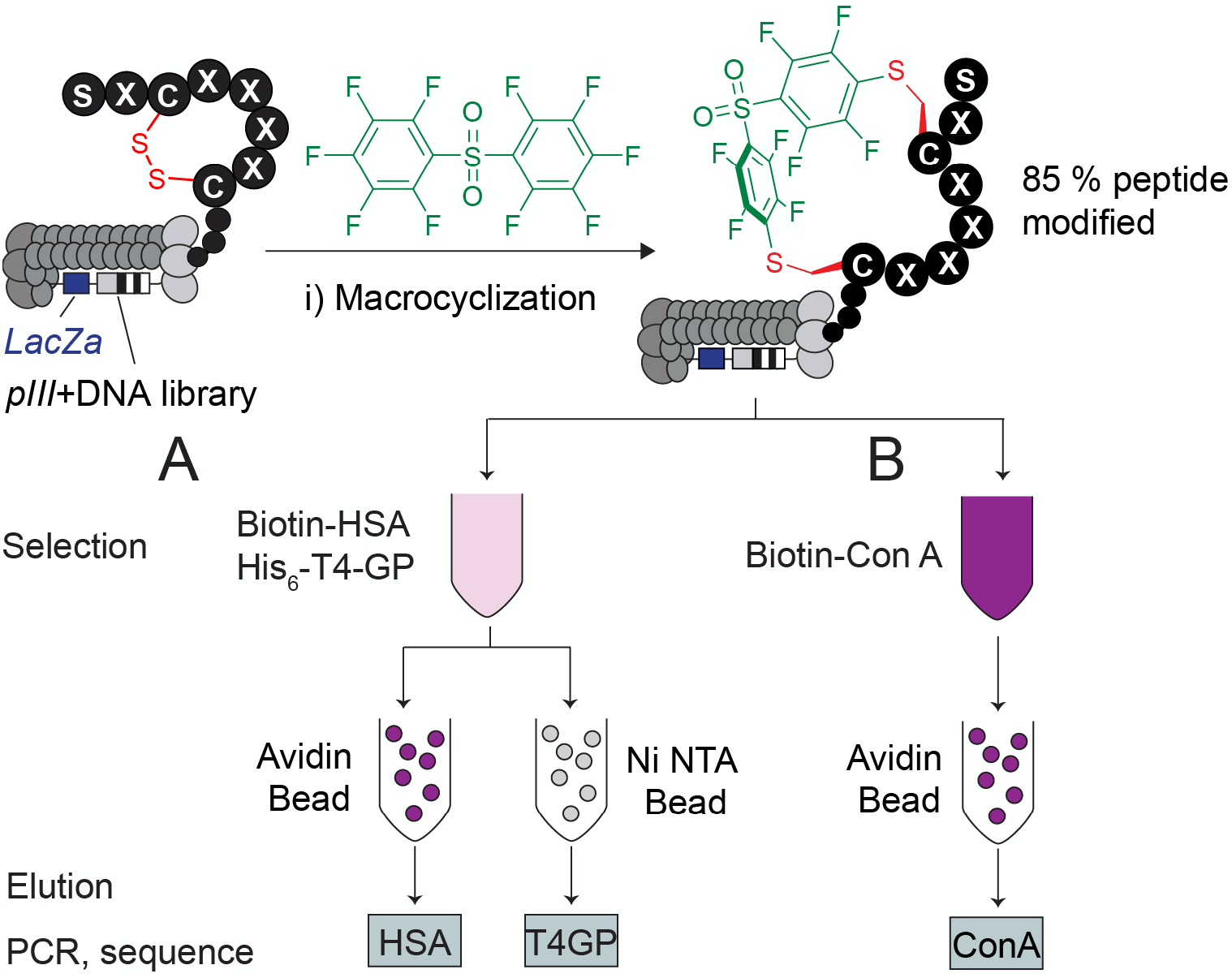
Phage-displayed peptide modified by **DFS**. (A) The modified phage-displayed library panned against two targets (biotinylated HSA and His-tag expressed T4-GP) in solution and captured separately with avidin beads and Ni-NTA beads affinity beads. (B) In the negative control, phage-displayed library modified by **DFS** was panned against biotinylated ConA and captured with avidin beads.

We performed three rounds of phage selection using HSA coated to the surface of 96 well polystyrene plates as bait. In parallel, we screened the same library on polystyrene wells coated with Protein A (negative control) to distinguish specific HSA-binding sequences from poly-specific protein binding sequences (**Figure S1A**). In round 3, the phage recovery of the **OFS**-macrocycle library selection against HSA two-fold increase compared to round 1 and round 2 but only a minor increase compared to selection against unrelated protein (**Figure S1D**). The recovery of unmodified round 3-library panned against HSA was 17-fold lower than the recovery of the **OFS**-macrocycle library, indicating that the **OFS** linchpin contributes to protein binding (**Figure S1D**). Differential enrichment (DE) analysis of the next-generation sequencing (NGS) of all test and control experiments (Table S1) identified several families of peptide macrocycles that had statistically significantly higher (*p*<0.05) enrichment in binding to HSA when compared to binding to unrelated protein (**Figure S1 B–C**). The analysis yielded three consensus motifs: STCHDITC (**1a**), STCHYIGC (**2a**) and STCHANC (**3a**) (**Figure S1E**).

The second discovery campaign employed HSA immobilized on a 96 well plate in rounds 1 and 3, and biotinylated HSA as bait immobilized onto streptavidin beads in round 2 (**Figure S2A**). In round 3, the phage recovery of the **OFS**-macrocycle library selection against HSA increased by a factor of 200 when compared to round 1 and round 2. The recovery of the unmodified library panned against HSA was insignificant (**Figure S2D**). The binding of the **OFS**-macrocycle phage library recovered from round 3 to Protein A, ConA and Casein was 2, 14, and 300-fold lower respectively when compared to recovery on HSA-coated wells (**Figure S3**). These observations suggested that (i) specific albumin-binding sequences had been selected, and (ii) the binding of these sequences to albumin required presence of **OFS** linchpin (**Figure S3**). A DE analysis of NGS data (**Figure S4, Table S2**) identified sequences that were significantly (*p*<0.05) enriched in the screen against HSA but not control proteins. The LOGO analysis yielded a consensus motif: STCHTIYC (**4a**) (**Figure S2E**). Although the original libraries were designed as SXCX_n_C where n=4 and 5, they contained a small fraction of SXCX_3_C sequences,^44^ and we observed the enrichment of such sequences in the selection. To explore the apparent preference for smaller macrocycles, we devised a third selection campaign that employed only SXCX_3_C libraries modified with **DFS** (**Figure 2A**).

The small diversity of the library made it possible to employ a single round of panning and NGS-DE analysis and to identify the binders. To mimic the complex serum environment, the panning was conducted using a mixture of biotinylated HSA (Bio-HSA), His-tag fusion T4-PG protein (His_6_-T4-PG) and unlabelled milk proteins as bait. In a control selection, we used the same mixture with biotinylated ConA (Bio-ConA) in place of Bio-HSA (**Figure 2B**). Proteins were captured with streptavidin or Ni-NTA affinity beads, respectively. The captured phage DNA was liberated from beads by treatment with hexane and the released DNA was amplified by PCR (**Figure S4**) and sequenced with Illumina deep sequencing (**Figure 2B, Table S3**). A DE analysis identified a set of 85 sequences that were significantly enriched (*p*<0.05, >3-fold) in the screen against Bio-HSA when compared to the screen against His_6_-T4-GP and Bio-ConA (**Figure 3A–B, Figure S5**). We applied a pairwise amino acid clustering to identify the 85 hit sequences (**Figure 3C**) and observed 8 motifs: FF, MF, MG, TK, GM, PV, VY and KR associated with these enriched sequences (Figure 3D). Based on this analysis, we nominated sequences SICRFFC (**5a**), SFCPMFC (**6a**) and SLCKREC (**7a**) as hits and STCQGEC (**8a**) as a negative control for chemical synthesis, and further validation (**Figure 3E**).

**Figure 3:**
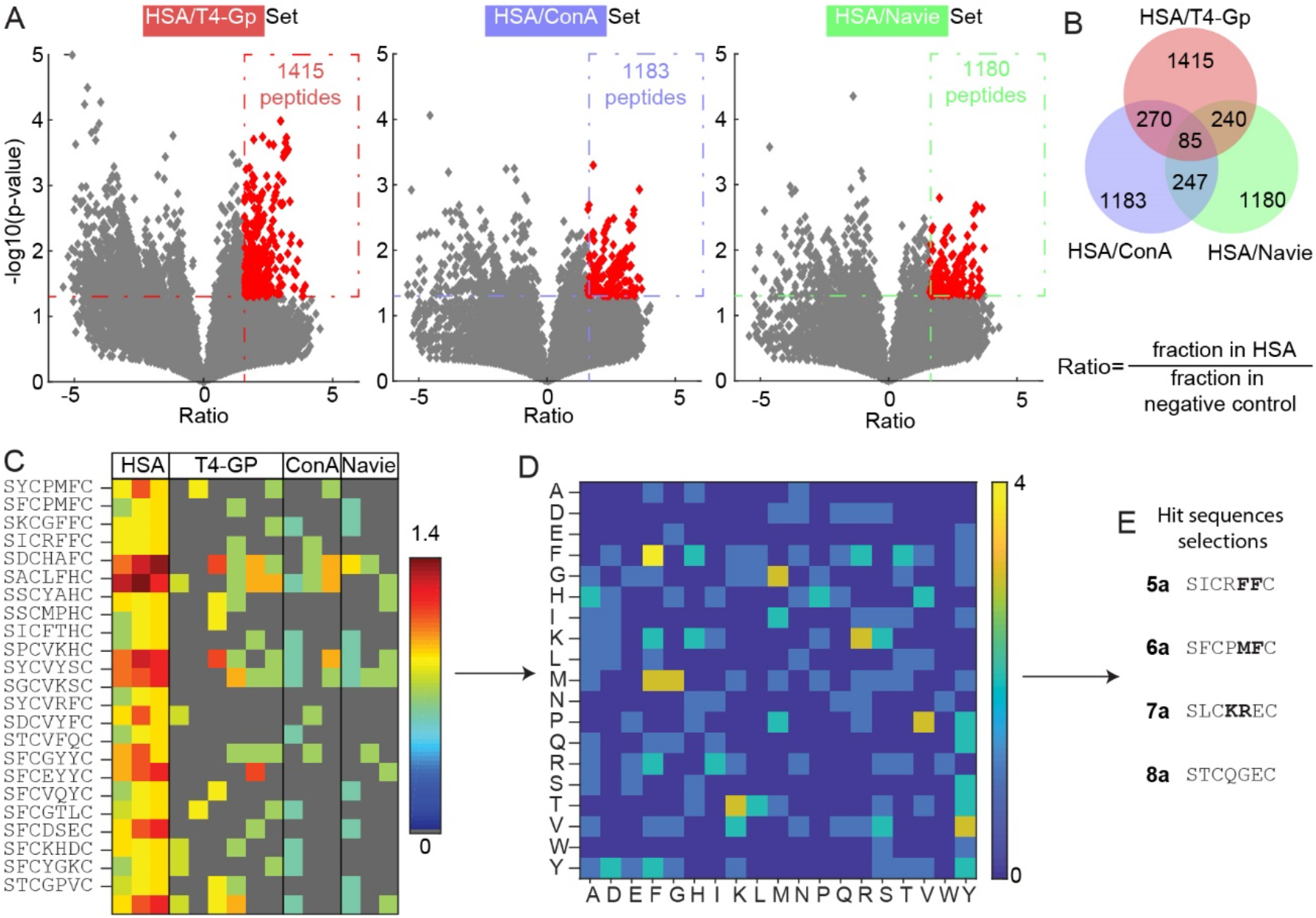
Student’s *t*-test analysis of the third screening campaign: (A, B) A volcano plot and venn diagram visualizing the sequences from the **OFS**-SXCX_3_C phage-displayed library that were significantly enriched in the HSA screen when compared to the naïve library or selection against T4-GP, ConA. (C) A heat map display of the top 25 of 85 hits sequences from differential enrichment results. (D) Dipeptide motif analysis of all 85 hits. (E) Selected sequences for chemical synthesis of macrocycles for validation.

### Validation of albumin binders

We observed non-specific reactivity of **OFS**-macrocyclic peptides with thiol nucleophiles such as glutathione (GSH) over several hours in basic pH (**Figure S6**). Replacing **DFS** with a less reactive pentafluorophenyl sulfide (**Figure 1D**) abolished the undesired reactivity: The resulting perfluorophenylsulfide (**PFS**)-macrocycles were unreactive to 2-mercaptoethanol over three weeks and unreactive towards free thiol on HSA (**Figure S7**). Molecular dynamics simulation suggested the **OFS**-macrocycles and the **PFS**-macrocycles exhibit similar ground state conformational landscape (**Figure S8**). Many perfluoro-aryl crosslinked macrocycles were poorly soluble in water, and we synthesized them with either a GGKKK or GGG tag at the C-terminus to increase their solubility; some sequences were synthesized with both tags to check whether these affect HSA binding. The C-terminal tags aided in providing sufficient solubility properties for downstream analyses (**Figure S9**).

The unique fluorine handle in perfluoro-aryl modified peptides made it possible to determine their binding to HSA using ^19^F NMR (**Figure 4**). In a typical experiment, we maintained peptide concentration at 50uM and HSA at 100uM (**Table S4**). We observed broadening of and disappearance of ^19^F signals that correspond to fluoroaromatic groups, which indicated the binding of the peptide to HSA (**Figure 4A**, **Figure S10**). We could not fit a definitive *K*_d_ value to the binding response due to the complex binding behaviour and quality of the NMR signal. However, in an albumin titration series, one can use qualitative estimates such as the concentration of albumin necessary to suppress 50% of the initial fluorine signal. Based on these qualitative analyses, it was apparent that some peptides (e.g., **PFS**-SICRFFCGGG) have stronger binding to HSA, whereas other macrocycles (e.g., **PFS**-STCQGECGGG) have weaker binding towards HSA (**Figure 4A**, **Figure S10**). By measuring the decrease in the signal at a fixed concentration of peptide and HSA, we evaluated 8 sequences found in all discovery campaigns (**Figure 4B, S11**) and we nominated **PFS**-SICRFFCGGG (**5c**) as the “hit” and **PFS**-STCQGECGGG (**8c**) as the negative control for further investigation. Peptides modified at the C-terminus with either GGKKK or GGG solubility tags have similar binding affinity (**Figure 4B, Figure S11**). We titrated **5c** and **8a** against rat serum albumin and observed similar binding to rat and human albumin (**Figure S12**). We attempted to confirm the binding affinity of these sequences by isothermal titration calorimetry (ITC) using SA-21 as a control;^37^ however, a complex multi-site binding behaviour for all peptides obscured the accurate evaluation of binding affinity by ITC (**Figure S13–15**). The ^19^F NMR assay, thus, was critically enabling for validation and ranking of the albumin binding leads.

**Figure 4:**
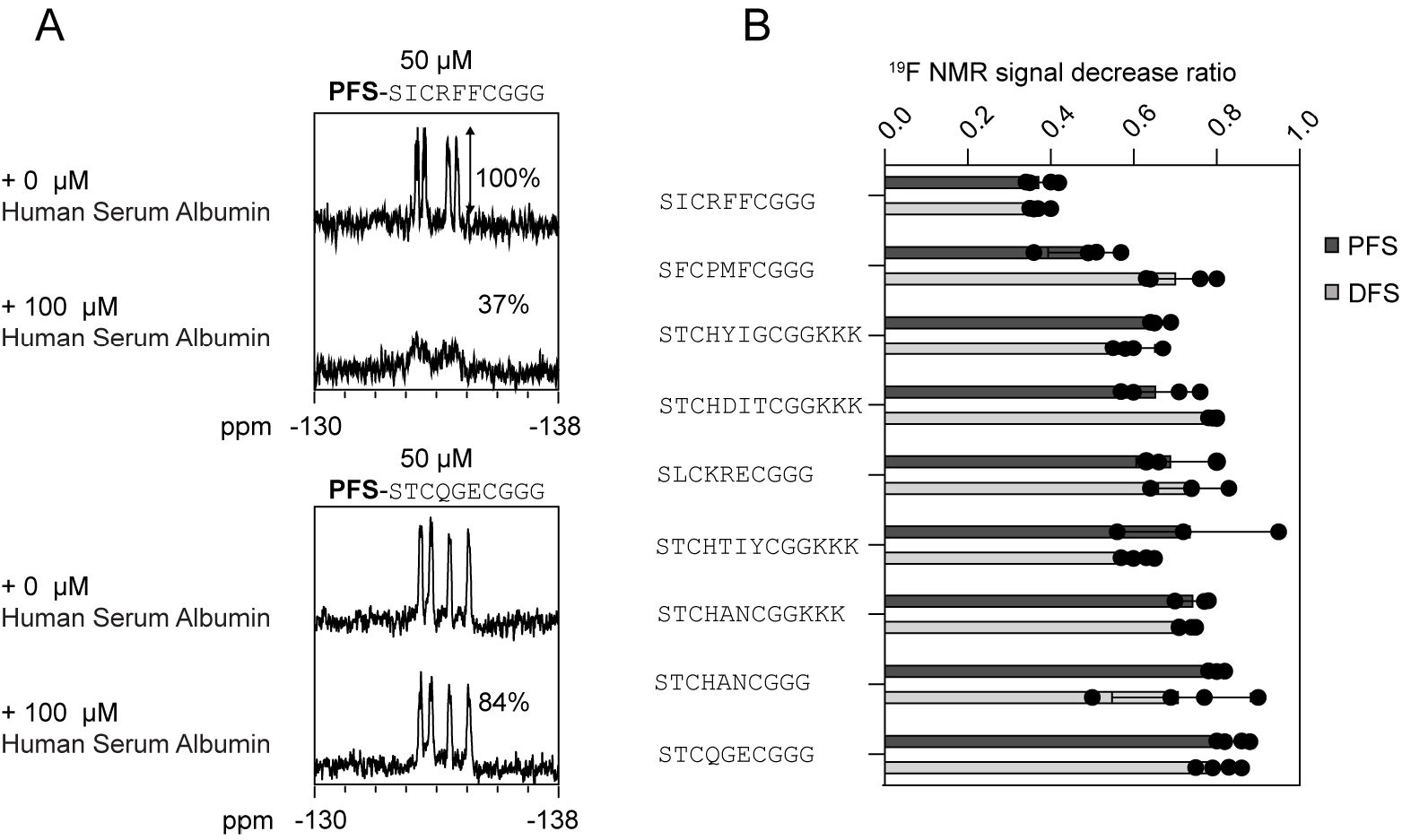
Summary of the ^19^F NMR binding measurement. (A) HSA titration spectra of 50 μM of **1b**-**8b** and **1c**-**8c** against 100 μM of HSA. (B) The percentage represents the remaining peak intensity following the addition of HSA.

A fluorescence polarization binding assay (FP) successfully measured the binding affinities of the macrocycles with the fluorophore BODIPY at the C- or N-terminus. In a typical experiment, we used **PFS**-SICRFFCGGG **(5c)**or **PFS**-SFCPMFCGGG **(6c)** at 1 μM concentration and titrated HSA from 0.1 μM to 100 μM. The dose-response curve could be fit to a single-state binding model with binding affinity of *K*_d_ = 4**–**6 μM for **5c** and at least 100 times weaker affinity for **6c** (**Figure 5A** and **S16–S19**). BODIPY alone bound weakly to HSA with > 300 μM binding affinity (**Figure 5A** and **S16–S19**). The FP-assay made it possible to measure binding to other proteins or even complex mixtures (serum). A titration of the mouse serum (**Figure S18**) yielded a similar binding profile to that observed in binding to pure albumin (**Figure 5A**). Replacing HSA with lysozyme and RNAse, the assay detected no binding response, confirming that **5c** binding was specific to HSA (**Figure 5B**).

**Figure 5:**
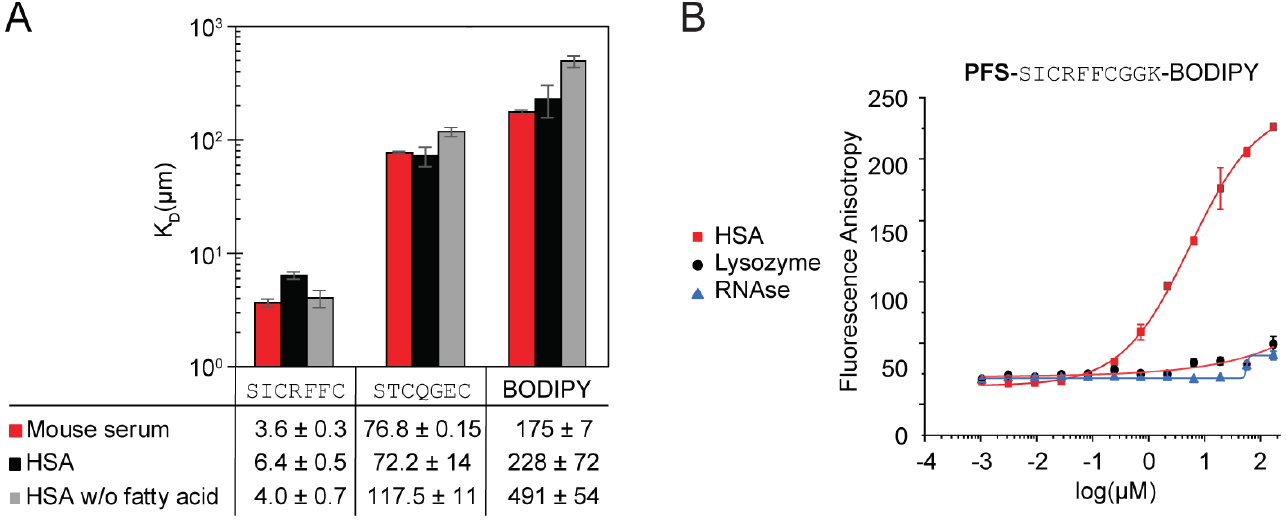
Binding assay data: (A) FP assay measured the *K*_d_ of the **5c** and **8c** against various albumins. (B) The FP assay for BODIPY labelled **5c** titrated against HSA (black), lysozyme (red), and RNAse A (blue).

Switching location of the fluorescent probe from the N-terminus to C-terminus did not significantly change the affinity of **5c** (*K_d_* = 4-6 μM, **Figure S19**). The switching from **DFS** to **PFS** also exhibited a minimal effect on the binding of peptides **5b** and **5c** (**Figure S20**). The results from FP were in the same order of magnitude as the semi-qualitative estimates acquired for BODIPY-free peptides by the ^19^F NMR binding assay, indicating that the presence of a fluorophore did not significantly increase the binding (**Figure S21**). Heinis and co-workers recently observed that fluorophores could dramatically increase binding affinity for albumin, and removing the fluorophore is detrimental to the binding of the albumin binder.^21^ To exclude this possibility, we conducted an NMR-binding assay of N-terminally-labelled **PFS**-SICRFFCGG and BODIPY-free peptides. We observed that the binding affinity was similar (**Figure S21**).

### Elucidation of the binding pocket for perfluoro-macrocycles

We evaluated whether binding pockets of **5c** are similar to known albumin binders: carbamazepine, diclofenac and ibuprofen (**Figure 6A**). We observed that the binding of **PFS**-SICRFFCGGG (**5c**) did not decrease in the presence of any of these drugs; thus, it does not share the same binding pocket as carbamazepine, diclofenac, or ibuprofen (**Figure 6B**). To follow on this observation, we performed a series of docking calculations to seek the most favorable binding locations of **PFS**-SICRFFCGGG (**5c**) on the surface of HSA.

**Figure 6:**
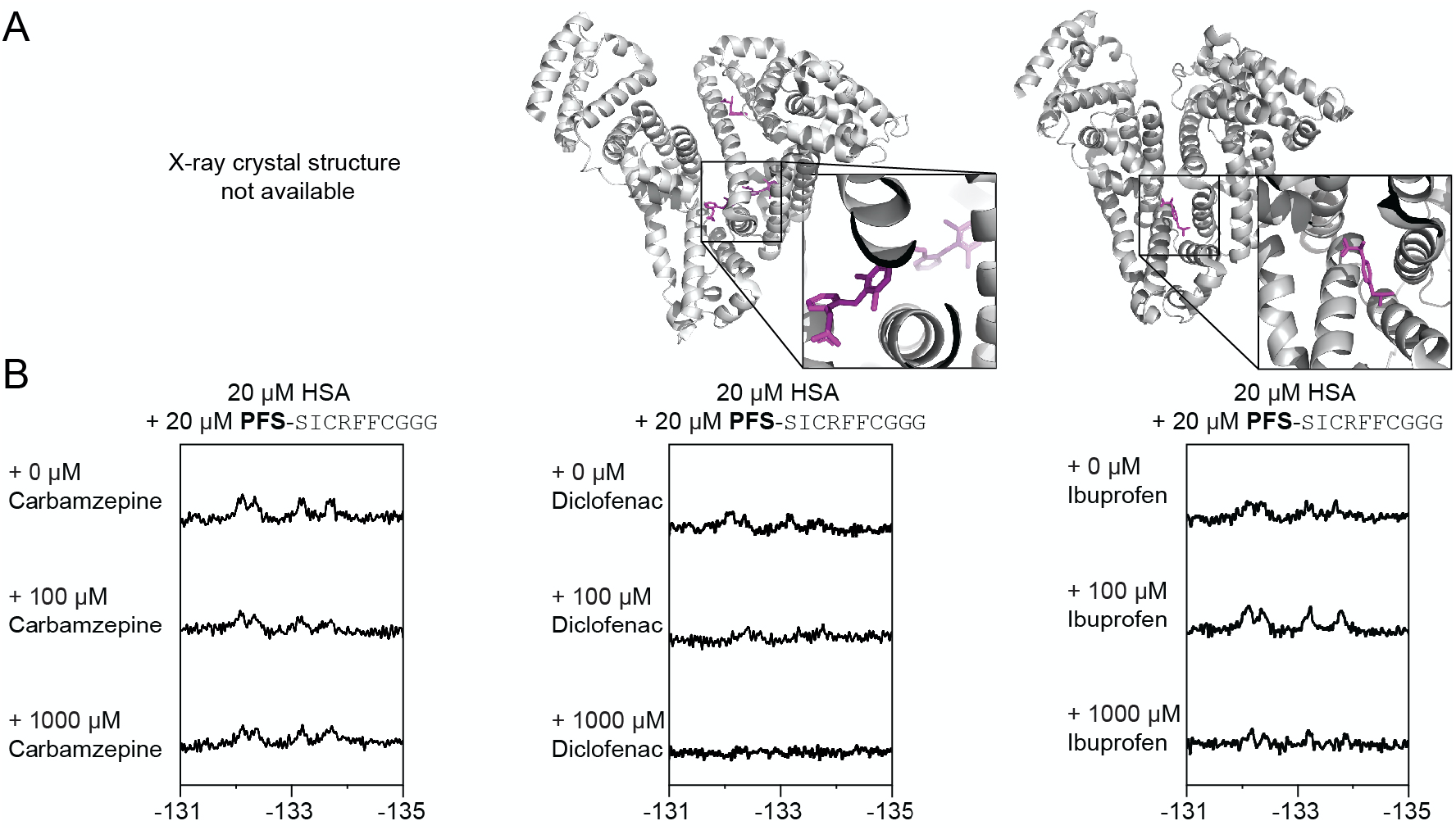
The ^19^F NMR HSA binding and competitive binding assay comparison. (A) X-ray crystal structure of diclofenac bound HSA (pbd: 4Z69) and ibuprofen bound HSA (pbd: 2BXG). (B) ^19^F NMR competitive inhibition assay with **5c** against carbamazepine, diclofenac and ibuprofen.

Nine distinct sites on HSA were previously shown to bind to fatty acids,^45^ some of which also bind to other ligands such as ibuprofen and diclofenac^46, 47^ (**Figure S22**). These nine reported fatty acid binding sites on five different initial HSA structures were selected for docking of **5c**. **Figure 7A** shows the HSA protein with some of its bound fatty acids, based on the pdbID 1e7e^45^. Overlaid with this structure is **5c** docked to the corresponding fatty acid binding sites on the HSA surface. **Figure 7B** shows the binding scores for **5c–** HSA complexes across different HSA structures, based on the distinct pdbIDs and different binding site locations (complete results summarized in **Figure S23** and **Table S5**). Consistently, **5c** has the most favorable binding score in binding site 1, with the value of – 8.95 ± 1.0 kcal/mol, averaged over all the docking calculations performed. Therefore, the results in **Figure 7B** suggest that the primary HSA binding site for **5c** is binding site 1. The next most favorable binding sites are sites 8, 6, and 7, with the most favorable binding scores of −6.6 ± 1.1 kcal/mol, −6.2 ± 0.7 kcal/mol, and −6.0 ± 1.2 kcal/mol, respectively (**Figure 7**, **Table S5**). We observed binding sites 1 and 8 are near to each other on the HSA surface, with the center of mass distance between fatty acids occupying these sites being 5.3 Å. As such, it is unlikely that binding sites 1 and 8 can be simultaneously occupied by two **5c** molecules. **Figure 7A** shows the four HSA residues that interact with the fatty acid in binding site 1 via charge and nonpolar interactions. In contrast, **PFS**-SICRFFCGGG (**5c**) has more interactions with this HSA binding site, including the HSA residues R114, R117, Y138, Y161, I142, L154, S193. Notably, HSA residues R117, Y138, and Y161 in binding site 1 are found to mediate HSA interactions with both the fatty acid and **5c**.

**Figure 7:**
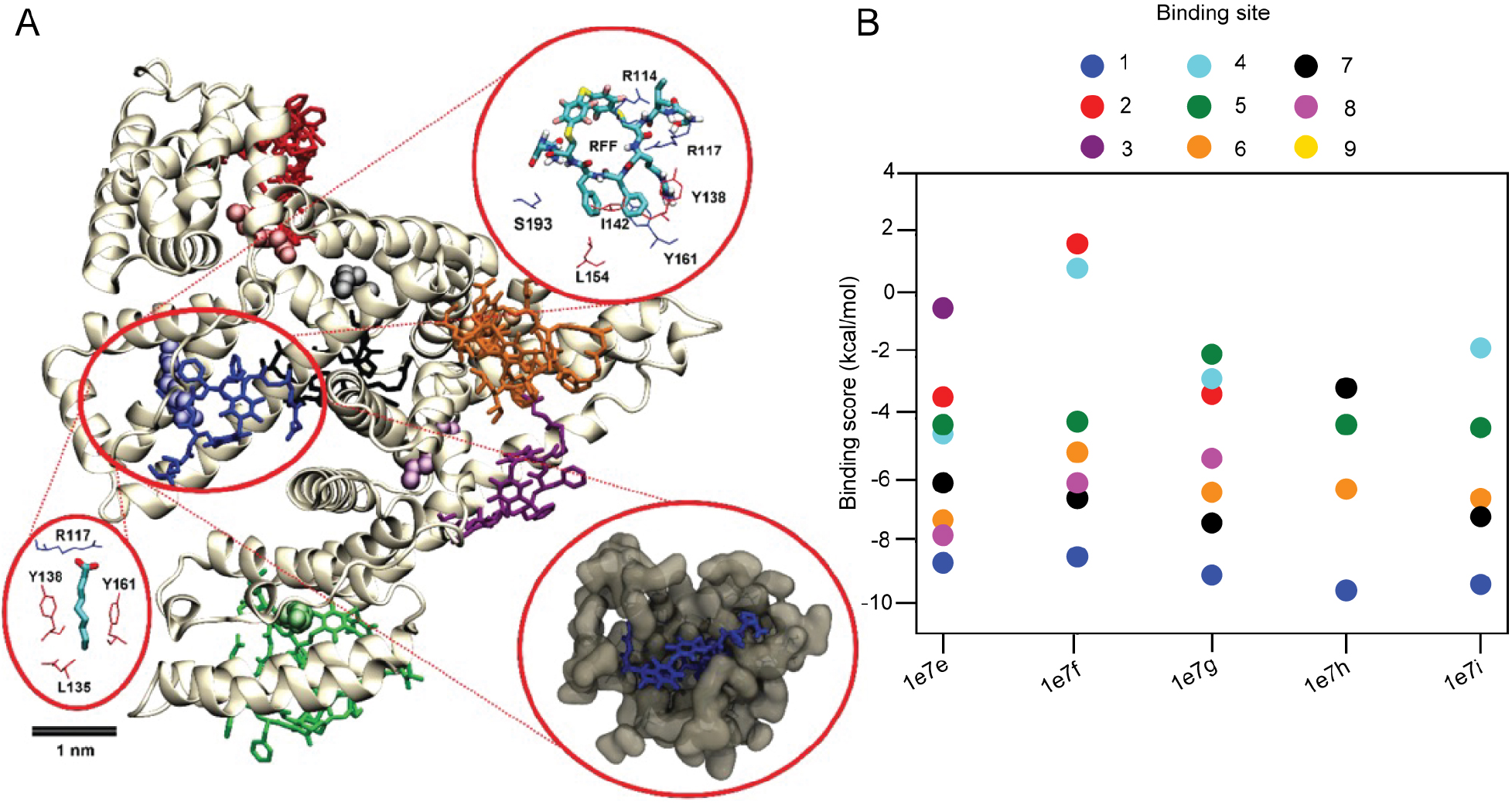
Docking calculations for **PFS**-SICRFFCGGG (**5c**) and HSA protein. (A) HSA protein structure with bound fatty acids (crystal structure-based), overlaid with **5c** docked to the respective fatty acid binding sites on the HSA surface. Binding site 1 is circled in red. The bottom left inset shows a fatty acid bound to the binding site 1 in the HSA crystal structure (pdbID 1e7e). The HSA amino acids with direct contact to the fatty acid are highlighted in thin licorice representation. Amino acids shown in blue form hydrogen bonds with the fatty acid, and amino acids shown in red form van der Waals (nonpolar) interactions with the fatty acid. The top inset shows **5c** docked in binding site 1. The HSA amino acids with direct contact to **5c** are highlighted in thin licorice representation. The bottom right inset shows a magnified **5c** docked into the binding site 1 pocket of HSA (grey surface). (B) Plots of **5c**–HSA binding scores obtained in one set of docking calculations. Separate docking calculations were performed for different HSA structures extracted from the PDB Databank files with PDB IDs: 1e7e, 1e7f, 1e7g, 1e7h, and 1e7i. **5c** was docked to the binding sites on the HSA surface that were occupied by the fatty acids in the crystal structures.^45^ Six of the nine HSA binding sites, the corresponding bound fatty acids, and docked **5c** are shown in panel A.

Combined docking results (**Figure 7**) and binding observation (**Figure 6**) suggested that ibuprofen, diclofenac and **5c** bind to different locations on the HSA surface. The structural studies^46^ demonstrated that ibuprofen binds to the binding sites labeled by 3/4 and 6 (pdbID 2bxg, **Figure S22**), which are distant from the HSA binding site 1. Structure of HSA bound to diclofenac^48^ (pdbID 4z69, **Figure S22**) contains two HSA chains in it. One of the HSA chains has a single diclofenac at the binding site 7, while the second HSA chain has three bound diclofenac ligands in total, with two also located at the binding site 7, and the third located near the binding site 1, which is also occupied by a bound fatty acid. The structure locations suggest that diclofenac has the strongest binding to binding site 7, since it is observed there in both HSA chains, and a weaker binding to binding site 1, as only one single HSA chain is observed with diclofenac nearby.

### Circulation lifetime of albumin peptides in mice

To evaluate the half-life circulation of albumin-binding perfluoro-macrocycles, we injected a mixture of peptides **PFS**-SICRFFCGG (**5c**), weak binding peptide **PFS**-STCQGECCGGG (**8c**) as the negative control and SA-21 as the positive control into mice and monitored the remaining peptide level by LC–MS (**Figure 8A**, **Figure S24**).

**Figure 8:**
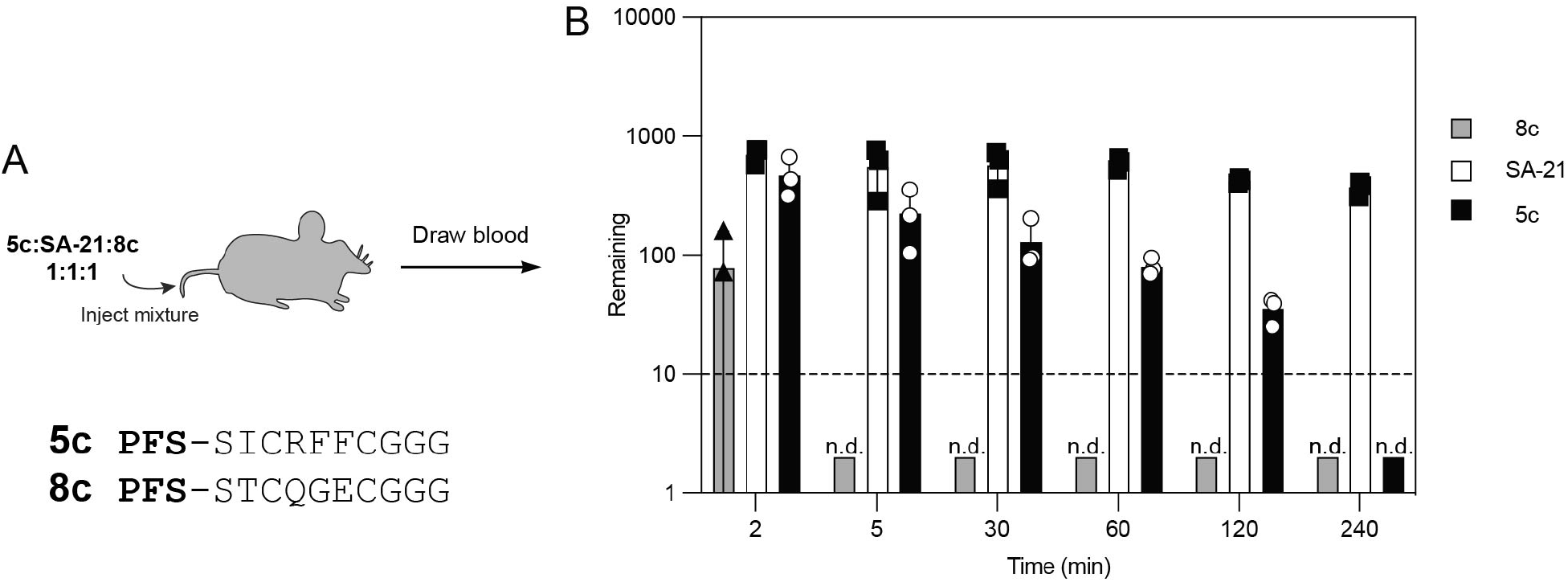
Pharmacokinetic studies for **5c**, **8c** and SA-21. (A) An equimolar (0.1 mM) mixture of macrocycles was injected into mice and blood was drawn at various time points. (B) Blood samples were collected at time points 2, 5, 30, 60, 120, and 240 mins and analyzed by LC-MS (*n* = 3). The dotted horizontal line represents the limit of detection. (n.d.: not detected).

We observed that the negative control **8c** disappeared below the limit of detection after 5 min (**Figure 8B**). The concentration of **5c** decreased 10-fold and SA-21 concentration decreased 5-fold after 2 hours. The combined results confirm a significant retention of **PFS**-SICRFFCGG peptide in circulation when compared to unrelated macrocyclic peptides with minimal to no detectable binding to HSA. The single digit micromolar peptide does not rival the mid-nanomolar SA-21 peptide, and the observed differences in half-life likely reflect the relative affinities for albumin. The **PFS**-SICRFFCGG peptide, thus, provides an attractive minimalistic starting point for further attenuation of binding affinity for albumin and subsequent attenuation of circulation half-life.

## Conclusion

Late-stage modification of peptides and genetically-encoded (GE) libraries of peptides by cross-linkers (linchpins) is one of the common approaches to incorporate beneficial attributes to their properties.^49^ Alkylation of cysteine residues in peptides via an S_N_2 reaction using bi- or tri-dentate alkyl halides has been use for cyclization of peptides, incorporation of unnatural fragments into the resulting macrocycles,^50–52^ and late-stage modification of phage- and mRNA-displayed libraries to yield billion-scale GE libraries. Peptide cyclization via SNAr reaction with perfluoroarenes popularized by the Pentelute group forms alkyl-aryl thioethers; ^43^ other classes reactions have been developed to form aryl^53–55^, alkenyl and alkynyl thioethers^56^ in unprotected peptides. Aryl and perfluoroaryl thioethers are more resistant toward oxidation when compared to traditional bis-alkyl thioesters^42^. Decreased conformational mobility or aryl-thioether bond has been proposed to equip the resulting macrocycles with favourable properties such as cell permeability and proteolytic stability.^45, 57^ Our report described the first selection from perfluoro-aryl macrocyclic GE libraries. There exists only one example of GE selection from SNAr-modified phage libraries: Lu and co-workers recently employed 2,4-Difluoro-6-hydroxy-1,3,5-benzenetricarbonitrile (DFB) as a reagent that can modify phage libraries in water.^58^ Chen and co-workers also used Pd-catalyzed SNAr reaction to yield DNA-encoded libraries.^59^ Both SNAr reaction yield macrocycles do not contain any fluorine atoms. On the other hand, fluorine handles present in perfluoroaryl-crosslinked macrocycles offer a unique possibility to use of ^19^F NMR to measure protein-macrocycle interactions. Interaction of perfluorinated aryls with proteins is also electronically distinct from non-fluorinated aromatic residues and in some cases it can offer uniquely advantageous interactions^60^.

An important observation in selection of GE **OFS**-macrocycle libraries is mild reactivity of these structures towards thiol nucleophiles.^61^ Libraries of mild electrophiles^62–64^ and phage-displayed libraries with built-in electrophiles^65^ have emerged as important starting point for discovery of covalent and reversibly covalent inhibitors. While we do not show it in our report, it is possible that an attenuated reactivity of **OFS**-macrocycles towards thiols can be used as an advantageous features in discovery or inhibitors that form covalent bonds with thiol residues in proteins. If reactivity of the selected macrocycles is not desired, one can perform late-stage replacement of **OFS** moiety in the identified hits with nearly isosteric perfluorophenyl-sulfide. The **PFS** linchpin is not sufficiently reactive for direct modification of phage-displayed libraries in water, but replacement of **DFS** linchpin by **PFS** “post discovery” maintains the conformation and binding affinity of the discovered macrocycles while alleviating their undesired electrophilicity. Our report, thus, suggest a general approach for the future utility of perfluoroaryl-modified libraries: Step 1: Select phage-displayed libraries of **OFS**-macrocycles against the desired target. Step 2: evaluate **PFS**-modified synthetic macrocycles for their ability to bind to these targets.

Human serum albumin (HSA) target used in this publication is a commonly employed model target in screen of phage-displayed or DNA-encoded libraries (DEL) and traditional high-throughput screening (HTS). Albumin is a complex multi-pocket receptor with regions that can bind to fatty acid-like moieties, dicarboxylic acids as well a wide variety of aromatic and heterocyclic compounds and large dye molecules.^46, 66^ Albumin also contains several binding sites for peptides as well as small proteins that have been utilized for half-life extension strategies.^21, 67^ Peptide macrocycles discovered in this report add to a diverse set of known albumin binders and they constitute the first example of small macrocyclic peptide binders for albumin. We note that single digit micromolar affinity of the discovered **PFS**-SICRFFC motif was not sufficient to retain this macrocycle in circulation as effectively as benchmark SA-21 albumin binding peptide. However, it should be relatively straightforward to optimize this structure to improve the affinity because our docking calculations suggest that a C-terminal extension to this scaffold might be productive avenue for the future optimization. Exploration of such extensions could be done using perfluoroaryl modified phage-displayed libraries of SICRFFCX*_n_* peptides with several randomized C-terminal amino acids. Such optimization should yield a collection of albumin binders with a range of affinities for albumin and, in turn, a range of the circulation half-lives. The small size of such peptide-macrocycle families makes it trivial to make them by solid-phase synthesis or incorporate them a part of another sequence produced by solid phase synthesis. More importantly, the SICRFFC motif or optimized SICRFFCX*_N_* motifs emerging from phage display screen can be easily re-introduced into phage-displayed libraries to serve as a constant N-terminal albumin binding motif and giving rise to libraries with predictable circulation half-life.

## Supporting information

Supporting information document contains Supporting Figures S1–S62, Tables S1–S5, synthetic methods and characterization of compounds, details of phage display selection, next generation sequencing and bioinformatics analysis and all biochemical assays. Supporting data folder contains PDB files produced by docking.

## Acknowledgements

This research was supported by a research contract from the Ferring Research Institute, Natural Sciences and Engineering Research Council of Canada (NSERC, RGPIN-2016-402511 to R. D.), and NSERC Accelerator Supplement (to R. D.). This work was also supported by the National Institute of General Medical Sciences of the National Institutes of Health under award number R01GM124160 (PI: Y.-S.L.) Infrastructure support was provided by CFI New Leader Opportunity (to R. D.). We thank Dr. Randy Whittal for assistance with LCMS, and Mark Miskolzie for assistance with ^19^F NMR kinetics.

