## Supplementary Information for "Genetically-Encoded Discovery of Perfluoroaryl-Macrocycles that Bind to Albumin and Exhibit Extended Circulation *in-vivo*"

**Figure S35:** Synthesis summary of **5c**. 59
**Figure S36:** Synthesis summary of **6b**. 60
**Figure S37:** Synthesis summary of **6c**. 61
**Figure S38:** Synthesis summary of **7b**. 62
**Figure S39:** Synthesis summary of **7c**. 63
**Figure S40:** Synthesis summary of **8b**. 64
**Figure S41:** Synthesis summary of **8c**. 65
**Figure S42:** Synthesis summary of C-terminus labeled **5c**-BODIPY. 66
**Figure S43:** Synthesis summary of N-terminus labeled **5d-**BODIPY 67
**Figure S44:** Synthesis summary of N-terminus labeled **5b-**BODIPY 68
**Figure S45:** Synthesis summary of C-terminus labeled **8c-**BODIPY. 69
**Figure S46:** STCHANCGGKKK $^1$H NMR Spectra in $D_2O$, 600 70
**Figure S47:** STCHDITCGGKKK $^1$H NMR Spectra in $D_2O$, 600 70
**Figure S48:** STCHYIGCGGKKK $^1$H NMR Spectra in $D_2O$, 600 71
**Figure S49:** STCHTIYCGGKKK $^1$H NMR Spectra in $D_2O$, 600 71
**Figure S50:** **DFS**-STCHANCGGKKK $^1$H NMR Spectra in $D_2O$, 600 72
**Figure S51:** **DFS**-STCHDITCGGKKK $^1$H NMR Spectra in $D_2O$, 600 72
**Figure S52:** **DFS**-STCHYIGCGGKKK $^1$H NMR Spectra in $D_2O$, 600 73
**Figure S53:** **DFS**-STCHTIYCGGKKK $^1$H NMR Spectra in $D_2O$, 600 73
**Figure S54:** **PFS** stapled STCHANCGGKKK $^1$H NMR Spectra in $D_2O$, 600 74
**Figure S55:** **PFS**-STCHDITCGGKKK $^1$H NMR Spectra in $D_2O$, 600 74
**Figure S60:** **PFS**-STCHTIYCGGKKK $^1$H NMR Spectra in $D_2O$, 600 75
**Figure S57:** **PFS** stapled STCHYIGCGGKKK $^1$H NMR Spectra in $D_2O$, 600 75
**Figure S58:** SLCKRECGGG $^1$H NMR Spectra in $D_2O$, 600 76
**Figure S59:** SICRFFCGGG $^1$H NMR Spectra in $D_2O$, 600 76
**Figure S60:** STCQGECGGG $^1$H NMR Spectra in $D_2O$, 600 77
**Figure S61:** **PFS** stapled SFCPMFCGGG $^1$H NMR Spectra in DMSO, 700 77
**Figure S62:** **PFS** stapled SLCKRECGGG $^1$H NMR Spectra in DMSO, 700 78
**Figure S63:** **PFS** stapled SICRFFCGGG $^1$H NMR Spectra in DMSO, 700 78
**21.** References 79

**List of abbreviations**:

| | |
|---|---|
| BIA | biotin-PEG2-iodoacetamide |
| Boc | *tert*-butyloxycarbonyl |
| BODIPY | 4,4-difluoro-4-bora-3a,4a-diaza-s-indacene |
| BSH | biotin-thiol |
| Calc | calculated |
| Da. | daltons(s) |
| DFS | |
| DMF | N, N-Dimethylformamide |
| ESI | electrospray ionization |
| eq. | equivalent(s) |
| EDT | 1,2-ethanedithiol |
| h | hour(s) |
| HSA | Human serum albumin |
| HPLC | high performance liquid chromatography |
| HRMS | high-resolution mass spectrometry |
| LCMS | liquid chromatography mass spectrometry |
| MHz | megahertz |
| MsCl | methanesulfonyl chloride |
| NMR | nuclear magnetic resonance |
| mL | milliliter(s) |
| mM | millimolar |
| min | minute(s) |
| mmol | millimoles |
| PBS | phosphate buffered saline |
| PCR | Polymerase chain reaction |
| PFS | pentafluorophenylsulfide |
| ppm | parts per million |
| rt | room temperature |
| RSA | Rat serum albumin |
| TCEP | tris(2-carboxyethyl)phosphine) |
| TIPS | triisopropylsilane |
| TFA | trifluoroacetic acid |
| Tris | tris(hydroxymethyl)aminomethane |
| v/v | volume/volume |

**1.** Biochemistry method

HSA was purchased from Sigma-Aldrich (cat# A4327-1G), as were Protein A from Sigma-Aldrich (cat# P6031-1MG), and Concanavalin A (cat# C2010-100MG). The proteins were immobilized on a high binding plate (Corning, ref# 3369) with 100 μL of HSA or Protein A (100 μg/mL). The wells were washed 6 times with 200 μL of 1×PBS-T+0.1% tween 20, at pH 7.4 prior to phage incubation. The proteins were biotinylated in section **6**. Prior to capture proteins, the magnetic streptavidin beads (Promega, cat# Z5482) were washed with 1 × PBS. All proteins were captures with 20 μL magnetic streptavidin beads.

**2.** Preparation of $SXCX_3C$ phage-displayed library

The procedures have been adopted and modified as previously described in two publications that produced the M13-displayed SXCXXXC library[1] and M13-SDB vector[2]. In short, the vector SB4 QFT*LHQ was digested with Kpn I HF (NEB cat# R3142S) and Eag I HF (NEB cat# R3505S). A primer/template pair consisting of primer 5'-AT GGC GCC CGG CCG AAC CTC CAC C-3' and template 5'-CC CGG GTA CCT TTC TAT TCT CAC TCT TCT X TGT XXX TGT GGT GGA GGT TCG GCC GGG CGC TTG ATT-3' with 'X' representing a trinucleotide formed by annealing. The primer/template was then extended using Klenow DNA polymerase (NEB) according to the manufacturer's instructions. The insert fragment was then digested with Kpn1 HF and Eag1 HF, gel purified and ligated into the cut vector. The ligation products were then transformed into electrocompetent *E. coli* cells and the transformants were grown overnight on *E. coli* TG1 to allow for phage production. Phage cultures were then centrifuged to remove cells and debris and then the phage was precipitated by PEG precipitation (5% PEG 0.5 M NaCl). Other SD vectors have been processed identically. We sequenced the naïve libraries by Illumina sequencing and the naïve library of $SXCX_nC$ ($n$=3-5) composition are publicly available at the following links: https://48hd.cloud/file/1470

**3.** Panning strategy 1: panning on plate

$SXCX_4C$ and $SXCX_5C$ libraries were prepared as the described previous reports protocol.[3]

Round 1-3: The following protocol was repeated 3 times.

In Protein A coated wells, 100 μL of $2\times10^9$ PFU/mL **DFS** modified library or unmodified library was incubated for O/N at 4 °C to remove unspecific bindings. The **DFS** modified library supernatant was then transferred to wells with HSA and incubated for 1.5 h at RT. In parallel, **DFS** modified library supernatants were also incubated with Protein A, and unmodified libraries were incubated with HSA and protein as a negative control. After panning, all wells were washed 10 times with 1×PBST. The phage remaining in the wells were eluted with 200 μL of glycine elution buffer (Glycine-HCl pH 2.2, 0.1% BSA) for 9 min. The elution buffer was transferred into a new 1.7 mL microcentrifuge tube and neutralized with 20 μL of 1 M Tris-HCl (pH 9.1). The recovered phage solution was amplified for the next round of bio panning and for deep sequencing.

**4.** Panning strategy 2: panning on plate and in solution

Round 1:

In Protein A coated wells, 100 μL of $2\times10^9$ PFU/mL **DFS** modified library or unmodified library was incubated for O/N h at 4 °C to remove unspecific bindings. The **DFS** modified library supernatant was then transferred to wells with HSA and incubated for 1.5 h at RT. In parallel, **DFS** modified library supernatants were also incubated with Protein A, and unmodified libraries were incubated with HSA and protein A as a negative control. After panning, all wells were washed 10 times with 1×PBST. The phage remaining in the wells were eluted with 200 μL of glycine elution buffer (Glycine-HCl pH 2.2, 0.1% BSA) for 9 min. The elution buffer was transferred into a new 1.7 mL microcentrifuge tube and neutralized with 20 μL of 1 M Tris-HCl (pH 9.1). The recovered phage solution was amplified for the next round of bio-panning and for deep sequencing.

Round 2:

50 μL of magnetic streptavidin beads transferred to a 1.7 mL centrifuge tube. 1 mL of PBS+0.1% Tween was added to wash the beads. The beads then responded in 1 mL of blocking buffer (PBS+2% milk) for 1 hour at 4 °C. In parallel, 200 μL ($2\times10^9$ PFU/mL) of **DFS** modified phage and unmodified phage library were incubated for 1 hour at RT with pre-block beads to delete beads binders. The beads were captured with a magnetic rack and the supernatant was transferred to a new 1.7 mL centrifuge tube for panning. 100 μL of depleted **DFS** modified phage was combined with 10 μg of biotinylated HSA, increased the volume to 200 μL with 1×PBS and incubated for 1 hour at RT. In parallel, 100 μL of the depleted **DFS** modified phage was combined with 10 μg of biotinylated protein A, increased the volume to 200 μL with 1×PBS and incubated for 1 hour at RT, and 100 μL of depleted bead binder unmodified phage was combined with 10 μg of biotinylated protein A and HSA, increased the volume to 200 μL with 1×PBS and incubated for 1 hour at RT. After incubation, 50 μL of blocked streptavidin beads were added to the samples. Then, the beads were captured with a magnetic rack and washed 10 times with 1 mL of 1×PBST. The phage remaining on the beads were eluted with 200 μL of glycine elution buffer (Glycine-HCl pH 2.2, 0.1% BSA) for 9 min. The elution buffer was transferred into a new 1.7 mL microcentrifuge tube and neutralized with 20 μL of 1 M Tris-HCl (pH 9.1). The recovered phage solution was amplified for the next round of biopanning and for deep sequencing.

Round 3:

In Protein A coated wells, 100 µL of $2\times10^9$ PFU/mL **DFS** modified library or unmodified library was incubated for 1 h at 4 ºC to remove unspecific bindings. The **DFS** modified library supernatant was then transferred to wells with HSA and incubated for 1.5 h at RT. In parallel, **DFS** modified library supernatants were also incubated with Protein A, and unmodified libraries were incubated with HSA and protein as a negative control. After panning, all wells were washed 10 times with 1×PBST. The phage remaining in the wells were eluted with 200 µL of glycine elution buffer (Glycine-HCl pH 2.2, 0.1% BSA) for 9 min. The elution buffer was transferred into a new 1.7 mL microcentrifuge tube and neutralized with 20 µL of 1 M Tris-HCl (pH 9.1). The recovered phage solution was amplified for the next round of biopanning and for deep sequencing.

**5.** Panning strategy 3: panning in solution

Round 1: This strategy is 1 round of panning.

Magnetic streptavidin beads were blocked with blocking phage (phage were not able to PCR) overnight and washed 3 times with 1 mL of 1×PBS. **DFS** modified phage library ($2\times10^{11}$ pfu/mL) and blocking phage ($2\times10^{12}$ pfu/mL) were mixed in a 1:10 ratio and incubated with blocked streptavidin beads for 30 mins to depleted beads binders. The depleted **DFS** modified phages were incubated with 5 mg of biotinylated HSA and T4GP-$His_6$ in a total volume 100 µL in 1×PBS with 2% milk for 30 min at RT. In parallel, the depleted **DFS** modified phages were incubated with 5 mg of ConA-Bio in a total volume 100 µL in 1×PBS with 2% milk for 30 min at RT. After incubation, 25 µL of blocked magnetic streptavidin beads were added to each mixture and incubated for 20 mins at RT. Pelleted by magnetic and remove supernatant. The beads were captured with a magnetic rack and were washed 9 times with 1 mL of 1×PBS. At last, the beads were resuspended in 1 mL 1×PBS and incubated for 30 mins. The beads were captured with a magnetic rack, discarded the supernatant, and resuspended with 30 µl of hexane and 30 µl of water (DNAase free water) and shaken for 15 mins (1500 rmp). The samples were heated to 55 °C for 10 mins or until hexane completely evaporated. The reminding water was collected, further converted to Illumina compatible DNA for deep sequencing by Illumina Next seq by PCR.

**6.** General protocol for proteins biotinylation

Protein was dissolved to 1 mg/mL in 1×PBS at pH 7.4. 5 molar fold excess of EZ-Link Sulfo-NHS-Biotin (Thermofisher, cat# 21217) were added to the protein solution. The reaction mixture was incubated O/N at 4 °C. The next day, the protein was dialysis 3 times in 4 L of 1×PBS at pH 7.4. The biotinylated protein was captured with magnetic streptavidin beads to confirm biotinylation. The final concentration was determined with Nano-drop.

**7.** PCR amplification protocol for Illumina deep sequencing

Take 25 μL of eluted or amplified phage solution was used as a template for PCR with a total volume of 50 μL.

A Typical 50 μL PCR mixture contained:

| | | |
|---|---|---|
| 1. | 5× Phusion buffer | 10 μL |
| 2. | 10 mM dNTPs | 10 μL |
| 3. | Phusion® High-Fidelity DNA Polymerase (NEB, cat# M0530S) | 0.5 μL |
| 4. | Forward primer (3’-CAAGCAGAAGACGGCATACGAGATC GGTCTCGGCATTCCTGCTGAACCGCTCTTCCGATCTXXXXC CTTTCTATTCTCACTCT-5’, 10 μM) | 2.5 μL |
| 5. | Reverse primer (3’-AATGATACGGCGACCACCGAGATCTA CACTCTTTCCCTACACGACGCTCTTCCGATCTXXXXACAGT TTCGGCCGA-5’, 10 μM) | 2.5 μL |
| 6. | Template solution | 25 μL |
| 7. | Nuclease free water | 8.5 μL |

Thermocycler was preformed using the following setting:

a) 95 °C for 30 sec
b) 95 °C for 30 sec
c) 60.5 °C for 15 sec
d) 72 °C for 30 sec
e) Repeat step b) to d) 25 times
f) 72 °C for 5 min
g) hold at 4°C

**8.** Illumina sequencing of samples before and after panning.

The PCR products were produced by PCR as described in general PCR amplification protocol for Illumina deep sequencing with one exception: in amplification of libraries before panning (input), the volume of the template (phage solution) was 2 µL. All products were quantified by 2% (w/v) agarose gel in Tris-Borate-EDTA buffer at 100 volts for ~35 min using a low molecular weight DNA ladder as standard (NEB, cat# N3233S). PCR products that contain different indexing barcodes were pooled, allowing 10 ng of each product in the mixture. The mixture was purified by eGel, quantified by quBit and sequenced using the Illumina NextSeq paired-end 500/550 High Output Kit v2.5 (2×75 Cycles). Data were automatically uploaded to BaseSpace™ Sequence Hub. Processing of the data is described in section processing of Illumina data.

Processing of Illumina data

The Gzip compressed FASTQ files were downloaded from BaseSpace™ Sequence Hub. The files were converted into tables of DNA sequences and their counts per experiment. Briefly, FASTQ files were parsed based on unique multiplexing barcodes within the reads discarding any reads that contained a low-quality score. Mapping the forward (F) and reverse (R) barcoding regions, mapping of F and R priming regions allowing no more than one base substitution each and F-R read alignment allowing no mismatches between F and R reads yielded DNA sequences located between the priming regions. The files with DNA reads, raw counts, and mapped peptide modifications were uploaded to http://48hd.cloud/ server. Each experiment has a unique alphanumeric name and unique static URL in Table 3-1–3-3

**Table S1:** URL for 1st panning campaign deep sequencing results

| | Modification | HSA | Protein A |
|---|---|---|---|
| R1 | **DFS** | https://48hd.cloud/file/213 | https://48hd.cloud/file/214 |
| | none | https://48hd.cloud/file/217 | https://48hd.cloud/file/218 |
| R2 | **DFS** | https://48hd.cloud/file/213 | https://48hd.cloud/file/214 |
| | none | https://48hd.cloud/file/217 | https://48hd.cloud/file/218 |
| R3 | **DFS** | https://48hd.cloud/file/213 | https://48hd.cloud/file/214 |
| | none | https://48hd.cloud/file/217 | https://48hd.cloud/file/218 |

**Table S2:** URL for 2nd panning campaign deep sequencing results

| | Modification | HSA | Protein A |
|---|---|---|---|
| R1 | **DFS** | https://48hd.cloud/file/236 | https://48hd.cloud/file/237 |
| | none | https://48hd.cloud/file/421 | N/A |
| R2 | **DFS** | https://48hd.cloud/file/236 | https://48hd.cloud/file/237 |
| | none | https://48hd.cloud/file/421 | N/A |
| R3 | **DFS** | https://48hd.cloud/file/236 | https://48hd.cloud/file/237 |
| | none | https://48hd.cloud/file/421 | N/A |

**Table S3:** URL for 3rd panning campaign deep sequencing results

| | | HSA | T4-GP | ConA |
|---|---|---|---|---|
| R1 | Input + Modification | https://48hd.cloud/file/799 | https://48hd.cloud/file/799 | https://48hd.cloud/file/799 |
| | Elution | https://48hd.cloud/file/798 | https://48hd.cloud/file/797 | https://48hd.cloud/file/796 |

**9.** General chemistry method

LC–MS analysis of peptide modifications was obtained on Agilent Technologies 6130 LC–MS. A gradient of solvent A (MQ water) and solvent B (MeCN/$H_2O$ 95/5) was run at a flow rate of 0.5 mL/min (0-4.0 min 5% B; 4.0-5.0 min 5%→60% B; 5.0-5.5 min 60%→100% B; 5.5-7.5 100% B, 7.5-11 min 100%→5% B).

**10.** Peptide synthesis:

Peptides were synthesized on a PreludeX peptide synthesizer (Gyros Protein Technolgoes) by standard Fmoc solid chemistry using Rink Amide AM resin. Fmoc-protected amino acids, HBTU, Rink Amide AM resin were purchased from ChemPrep, Wellington FL USA. Peptides were cleaved from the resin by using a TFA/EDT/TIPS/Water (89.9/2.28/4.54/2.28 v/v) then precipitated and washed with ice-cold diethyl ether, and further purified by HPLC and lyophilized into the product.

General protocol for cyclization with decafluorodiphenylsulfone

Linear peptide (10 mM) was dissolved in 50% acetonitrile and Tris buffer (50 mM Tris-HCl, pH 8.5), then 2 equivalents of **DFS** in 50% acetonitrile and Tris buffer (50 mM Tris-HCl, pH 8.5) was added to the mixture. The mixture was vortex for 30 sec and let to proceed for 2 hours at room temperature. Then, the reaction mixture was purified by HPLC and further lyophilized into the product.

**11.** General protocol for cyclization with pentaflurophenyl-sulfide

Linear peptide (10 mM) was dissolved in 50mM Tris in DMF, then 2 equivalents of **PFS** was added to the mixture. The mixture was vortex for 30 sec and allow to react for 1 hour at RT. The reaction mixture was purified by HPLC and further lyophilized into the product.

**12.** BODIPY fluorescence C-terminus labeling of 5c and 8c

N-terminal Fmoc-protected **PFS** stapled peptides were dissolved in 1×PBS, 50% acetonitrile, then 1.5 equivalents of BODIPY-NHS ester (100 mg/mL DMSO, Anaspec) was added to the solution. The mixture was incubated for O/N at room temperature. Peperdine were add to a final concentration of 20% v/v for 30 mins to deportect the N-terminus Fmoc protecting group. The reaction mixture was purified by HPLC and further lyophilized into the product.

**13.** $^{19}$F NMR binding experiment

NMR experiments conducted at the University of Minnesota were performed on a Bruker Avance III HD with a Prodigy TCI cryoprobe (2100:1 S/N for $^{19}$F). **PFS**-peptides were tested as 20 μM solutions in experiments were performed with a fluorinated peptide concentration of 20 μM in 50 mM phosphate, 100 mM NaCl and 26.5 μM 2,2,2-trifluoroethanol, pH 7.4 with varying concentrations of rat or human serum albumin (from 0-160 μM). Parameters used for each experiment are as follows: 750 scans, acquisition time of 0.05 s, relaxation delay of 0.7 s, spectrum centered at -135 ppm with a sweep width of 20 ppm. A reference spectrum observing 2,2,2-trifluoroethanol was acquired for each sample with the following parameters: 16 scans, acquisition time of 0.5 s, relaxation delay of 1 s, spectrum centered at -75 with a sweep width of 10 ppm. The observed chemical shift of the reference was deducted from -77.75 ppm, and this difference was applied to the peptide spectrum.

**Table S4:** A typical experiment involving titration with HSA

| Component | HSA | **PFS**-peptide | TFE | $D_2O$ | PBS |
|---|---|---|---|---|---|
| Stock (μM) | 2000 | 200 | 26.5 | n.a | n.a |
| | 0 | 20 | 2 | 25 | 423 |
| | 2.5 | 20 | 2 | 25 | 420.5 |
| | 5 | 20 | 2 | 25 | 418 |
| | 10 | 20 | 2 | 25 | 413 |
| | 20 | 20 | 2 | 25 | 403 |
| | 40 | 20 | 2 | 25 | 383 |

n.a: Not applicable

This series would produce a titration with 20 μM **PFS**-peptide held constant and the HSA varying in a two-fold fashion: 0, 10, 20, 40, 80,160 μM. TFE refers to 2,2,2-trifluoroethanol as a 1/1000 dilution (approx. 26.5 μM), and PBS components were 50 mM phosphate, 100 mM NaCl, pH 7.4.

NMR experiments conducted at the University of Alberta were performed on an Agilent MR400 with a OneNMR™ Probe (690:1 S/N for $^{19}$F). **PFS** and **OFS** peptides were tested as 50uM solutions in 1X PBS, 2.5% DMSO, and 10% $D_2O$ with 100uM HSA. Parameters used for each experiment are as follows: 2048 scans, acquisition time of 0.15s, relaxation delay of 0.85s, spectrum was centered between **PFS** and **OFS** fluorine peaks and with a sweep width of 207 ppm. Peptide spectra were referenced to TFA at -76.55 ppm.

**14.** Fluorescence polarization binding assay

384 black well plates (PerkinElmer, cat# 6007270) were used to measure all binding assays. The fluorescent-labeled peptides were dissolved to 10 mM in DMSO and diluted to 20 μM in DMSO for use. Each well added with 19 μL of HSA in 1×PBS with the range of final concentration from 190 μM to 15 nM. 1 μL of 20 μM of fluorescence landed peptide added to the wells to a final concentration of 1 μM. Each measuring point was made in duplicate. Before measuring, the plate was spun down with 500×g for 5 mins at RT, incubated for 10 mins, and shake for 5 mins in the dark. The measurement was performed in Cytation5 Cell Imaging Multi-Mode Reader from BioTek. The data were analyzed and processed in OrginLab.

**15.** Isothermal titration calorimetry (ITC) binding assay

Titration experiments were performed using a Microcal VP-ITC instrument. Pepties were discoved in 1×PBS at pH 7.4 to a final concentration of 400 μM . In the case, where ligands have poor solubility in the prepared buffer, up to 5% (v/v) DMF was used. The HAS solution was prepared with the identical buffer as the peptides to a final concentration of 40 μM. All solutions were degassed with MicroCal ThermoVac. All the titration in this studiy carried out at 37 $^{o}$C and stirring at 300 rpm. An initial injection of 2 μL follwed by a total of 41 injection of 10 μL peptide solution were added at the interal of 4 mins into the HSA solution. The data were evaluated using the MicroCal$^{TM}$ Origin$^{TM}$ Version 5.0, and the heat signals were fitted to "one set of sites" or "two sets of sites" models to obtain the binding enthalpy, affinity, and stoichiometry estimates.

**16.** *In vivo* pharmacokinetic experiment

All the procedures and experiments involving animals were carried out using a protocol approved by the Health Sciences Laboratory Animal Services (HSLAS), University of Alberta. The protocol was approved as per the Canadian Council on Animal Care (CCAC) guidelines. All mice were maintained in pathogen-free conditions at the University of Alberta breeding facility. 50-100 μM of peptides mixture were dissolved in 1×PBS. Mice were administered 200 μL of the peptide mixture solution with tail veil injection. 6 blood samples were collected at serial time points from 1 min to 240 mins. Samples were collected in tubes that contained sodium citrate as an anticoagulant and then centrifuged at 5 min at 2,000×g to collect the blood plasma. 10 μL of plasma portion were transferred into a tube containing 40 μL of 8:2 acetonitrile/water to precipitate proteins. The sample was centrifuged at max speed for 10 min at 4 $^{o}$C. Supernatant then transferred to a clean tube and stored at -20 $^{o}$C for further LC–MS analysis.

**17.** LC–MS analysis for pharmacokinetics

LC–MS studies of the stability of peptides in mice were performed in Hewlett Packard 1100 series instrument using a Phenomenex Jupiter C4 protein column (300 Å,

2×50 mm, 0.3 mL/min, A: 0.1% formic acid in water, B: 0.1% formic acid in acetonitrile (0 min 2% B, 0→10 min 2%→70% B, 10→15 min 70% B, 15→20 min 70%→2% B). The amount of peptide remaining was calculated with the area under the curve of SIM (Selected Ion Monitoring) peak in LC–MS.

**18.** Docking Calculations

Docking of **PFS**-SICRFFCGGG macrocycle to the human serum albumin (HSA) protein was performed using five different HSA crystal structures obtained from the RCSB databank (PDB IDs: 1e7e, 1e7f, 1e7g, 1e7h, 1e7i).[4] In these crystal structures, multiple fatty acids (between six and nine) of different lengths are bound to HSA proteins. The previously characterized binding locations of the fatty acids on the HSA surface[4] are here called binding sites, labeled by distinct indices. In our calculations, we docked **PFS**-SICRFFCGGG macrocycles to each of those binding sites. The **PFS**-SICRFFCGGG structure was constructed with ChemSketch.[2] Both **PFS**-SICRFFCGGG and HSA were converted to Autodock Vina[5] readable format using MGLTools. In docking the **PFS**-SICRFFCGGG to HSA, the grid box center locations were selected to coincide with the centers of mass of the bound fatty acids. The grid boxes had 2.6 x 2.6 x 2.6 $nm^2$ dimensions with a default spacing of 0.0375 nm. The Autodock Vina configuration files and the docking run procedure were generated and ran automatically with a bash script provided by us on Github (https://github.com/vukoviclab/hsa-dock). The docking runs were performed using three different random seeds.

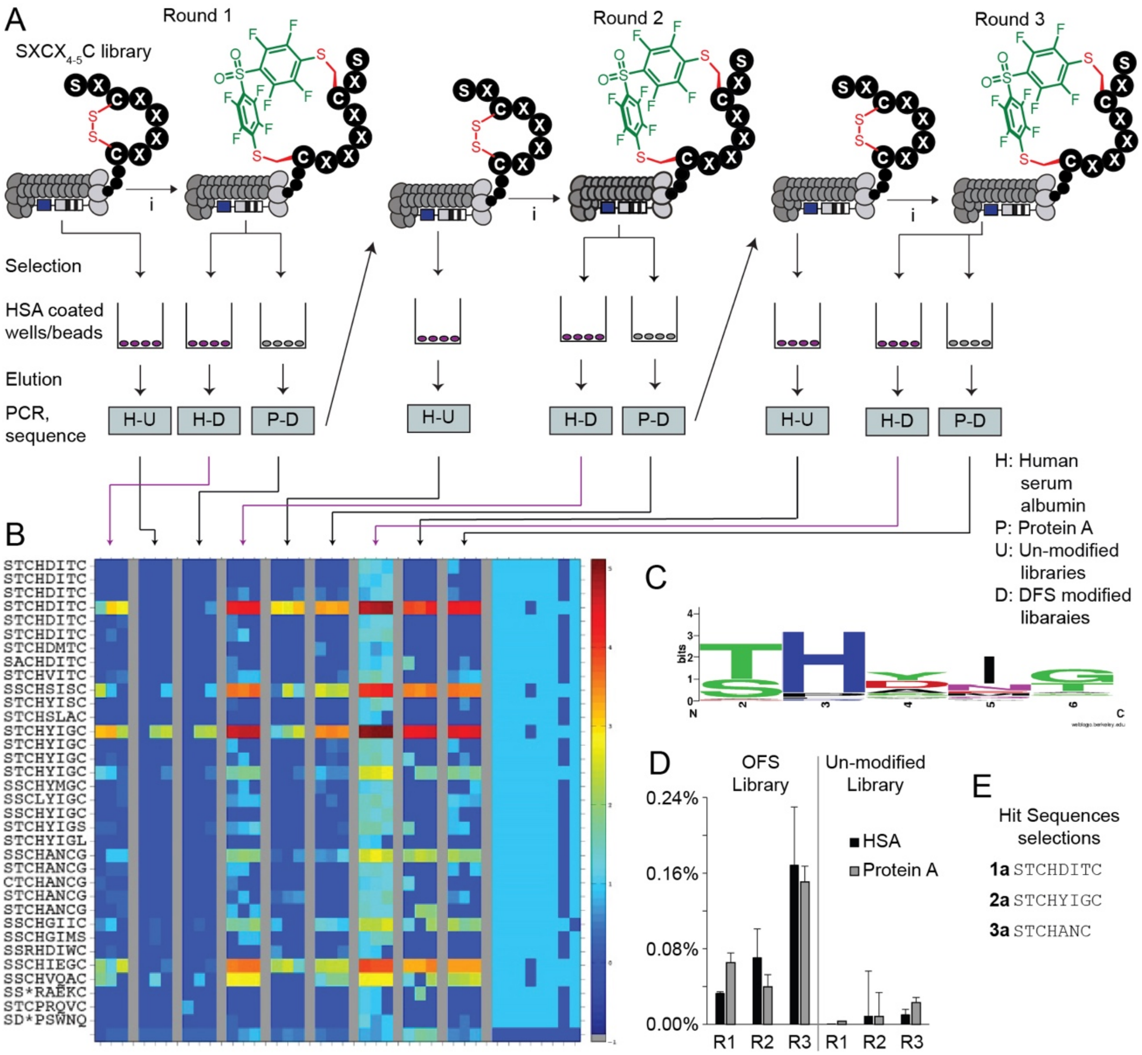

**Figure S1:** First bio-panning campaign of **OFS**-modified phage-displayed peptide library against the HSA proteins. (A) A scheme of three-rounds panning against HSA and negative controls. (B) The top 39 sequences from differential enrichment results. (C) LOGO analysis plot of the enriched sequences. (D) Percentage of the phage recovery after each round of bio-panning. (E) Selected sequences for chemical synthesis of the macrocycles.

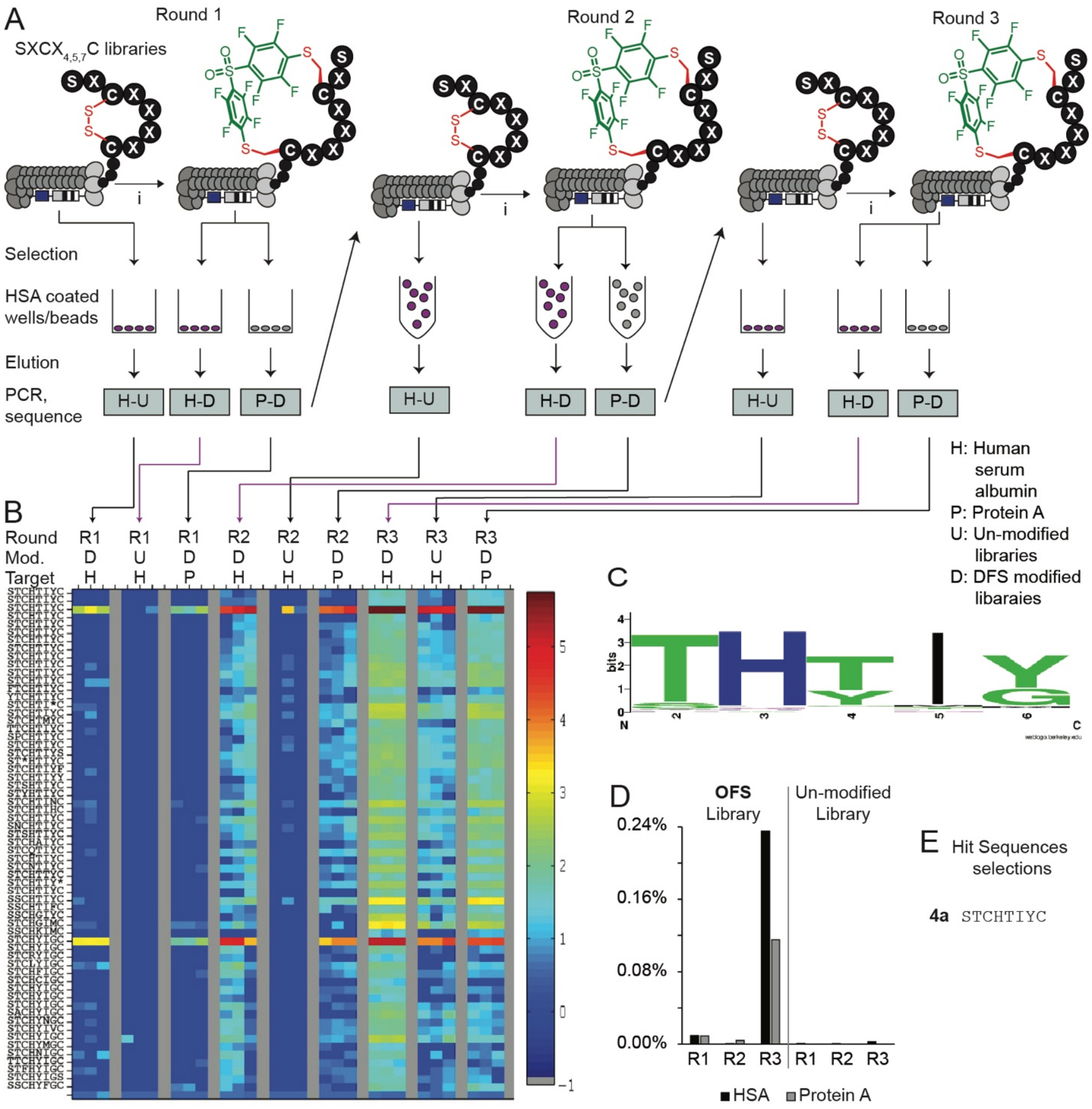

**Figure S2:** Second bio-panning campaign of **OFS-**modified phage-displayed peptide library against the HSA protein. (A) Scheme of a three-rounds panning against HSA and the negative controls. (B) The top 65 sequences from differential enrichment results. (C) LOGO analysis plot of the enriched sequences. (D) Percentage of the phage recovery after each round of bio-panning. (E) Selected sequences for validation synthesized into macrocycles.

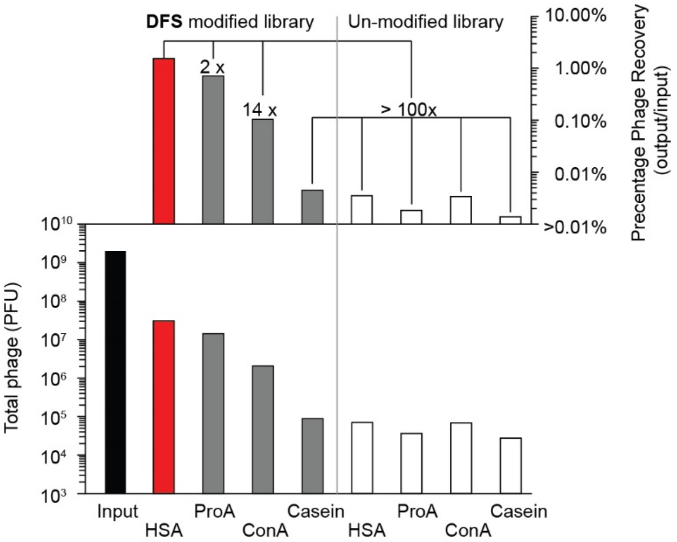

**Figure S3:** Phage recovered from the second selection campaign of round 3 were bound to different proteins on protein coated plates.

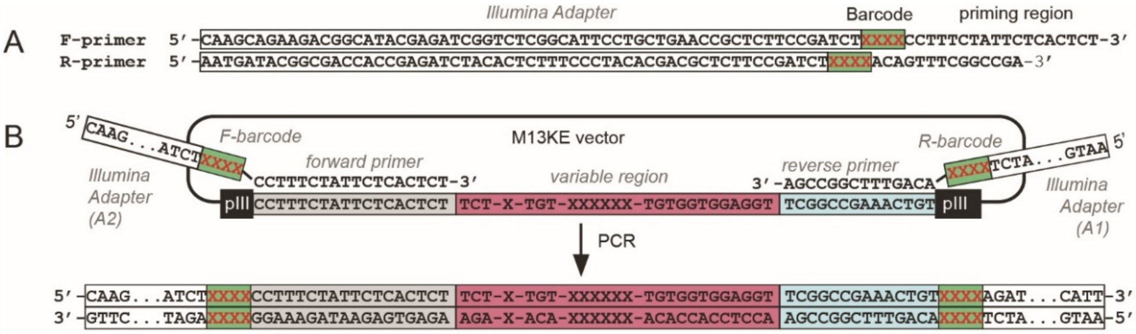

**Figure S4:** PCR amplification protocol for Illumina deep sequencing (A) Primers used for amplifying ligated or naïve oligonucleotide DNA. XXXX denotes 4-nucleotide-long barcodes used to trace multiple samples in an Illumina sequencing experiment. (B) Generation of PCR product. Alignment of forward and reverse primers to 18-bp and 14-bp sequences flanking the variable region at the N-terminus of the pIII gene in M13KE vector, respectively.

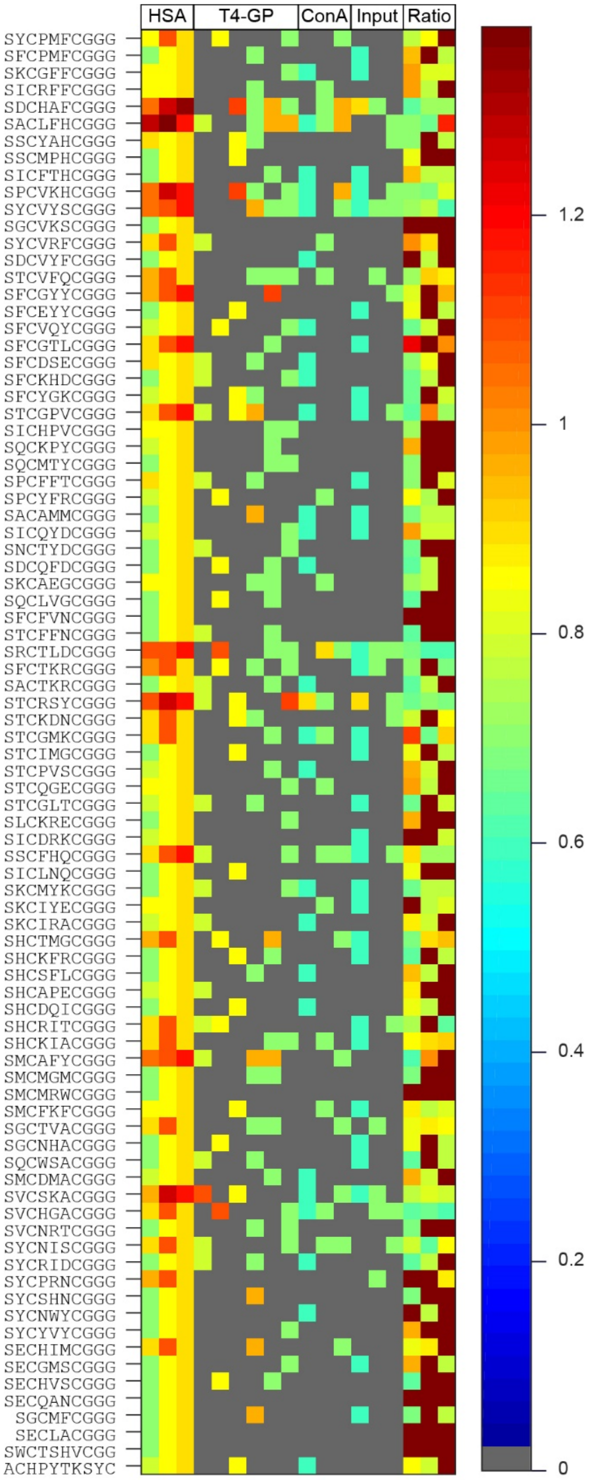

**Figure S5:** A heat map is showing of 85 putative hits discovered from the third panning campaign. The sequences were enriched greater or equal to 4-fold (R>3, p=0.05) when compared to T4-GP and ConA.

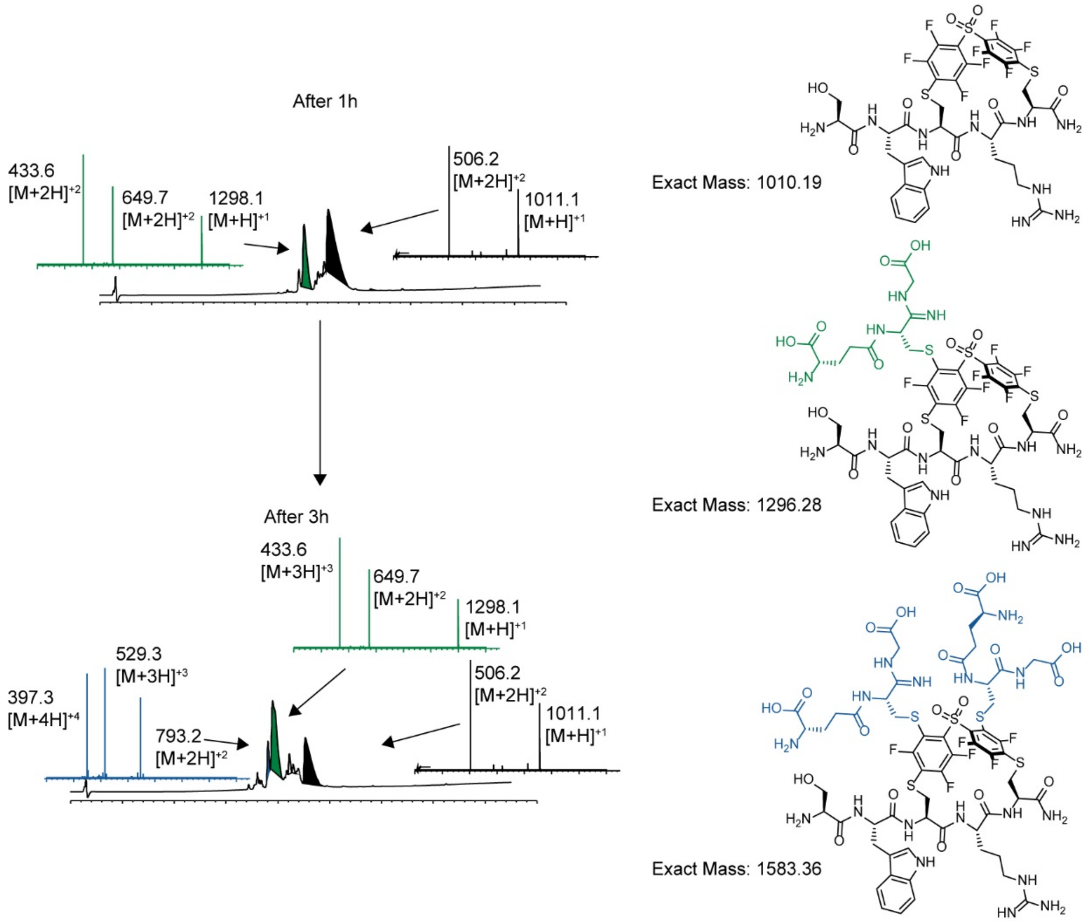

**Figure S6: DFS** stapled peptide reacted with GSH over 3 hours. The **DFS**-SWCRC peptide was added with one equivalent of GSH in 60% acetonitrile in 50 mM Tris-HCl at pH 8.5. The reaction was monitored by LC–MS.

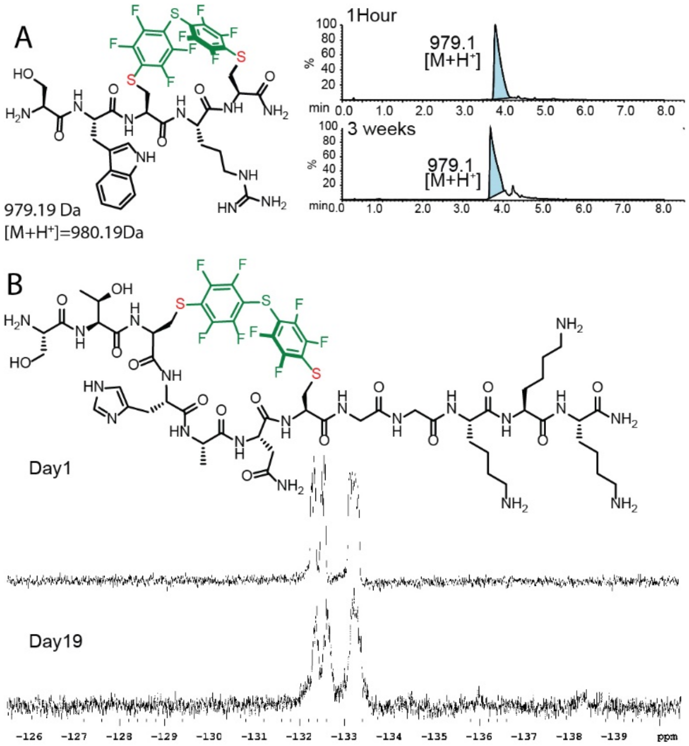

**Figure S7:** Stability of the **PFS** stapled peptides (A) Results of the **PFS**-SWCRC mixed with 2-mercaptoethanol and analyzed by LC–MS after 1 hour and 3 weeks. The **PFS**-SWCRC macrocycle was combined with 1 equivalent of 2-mercaptoethanol in 60% acetonitrile and 50 mM Tris-HCl at pH 8.5. The mixture was monitored by LC–MS. (B) A spectrum of 1 mg of **PFS**- STCHANCGGKKK mixed with 1 mg of HSA over 19 days in 1×PBS. The mixture was monitored by $^{19}$F NMR in 1×PBS, 10% $D_2O$

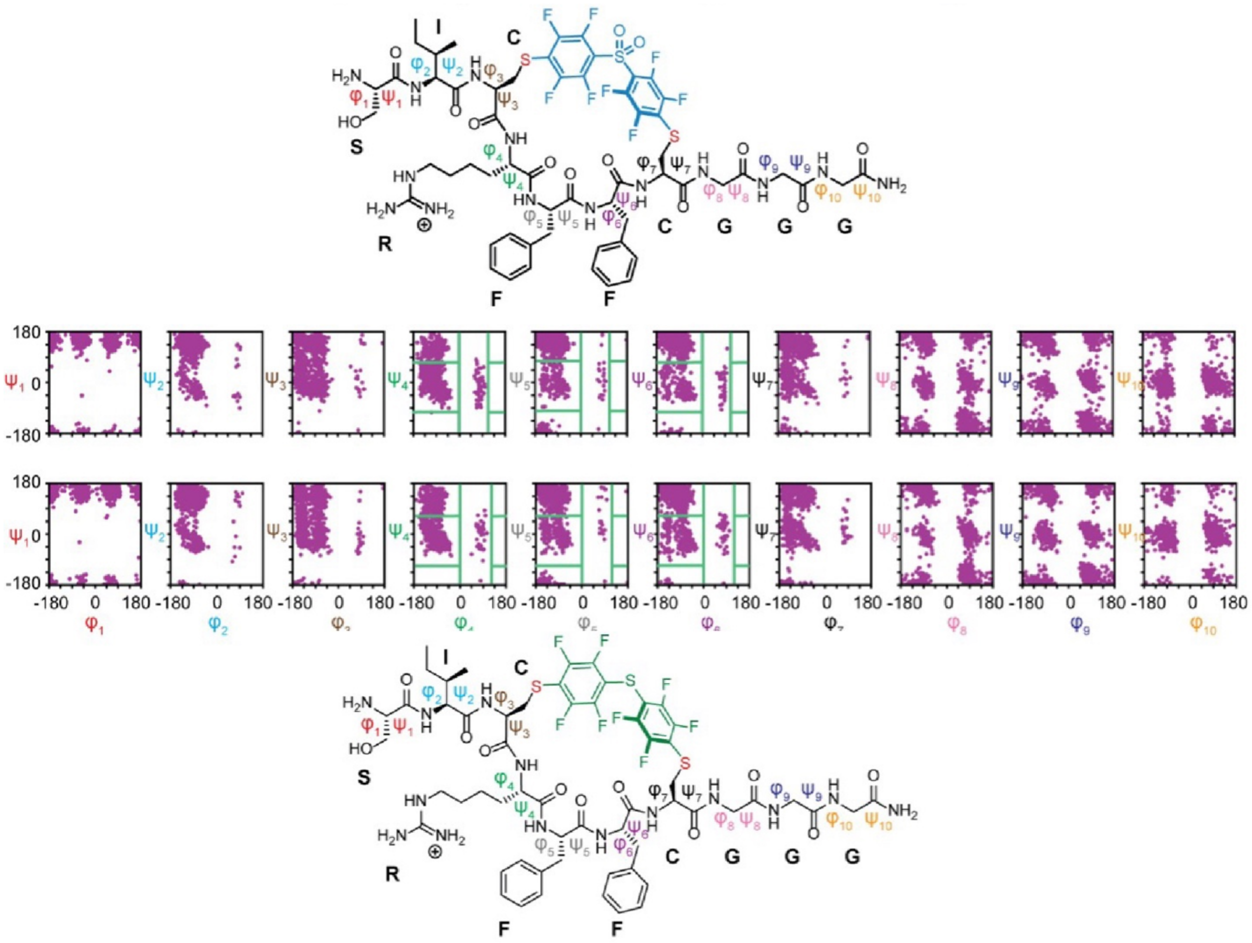

**Figure S8:** Ramachandran plot of the cyclic peptide backbone for **5b** and **5c**: Green lines indicate the binning boundaries used in the cluster analysis.

| Campaign # | | Sequences | **DFS** | Yield | **PFS** | Yield |
|---|---|---|---|---|---|---|
| 1 | **1a** | STCHDITCGGKKK | **1b** | 5% | **1c** | 30% |
| | **2a** | STCHYIGCGGKKK | **2b** | 64% | **2c** | 27% |
| | **$3a_1$** | STCHANCGGG | **$3b_1$** | 24% | **$3c_1$** | 40% |
| | **$3a_2$** | STCHANCGGKKK | **$3b_2$** | | **$3c_2$** | |
| 2 | **4a** | STCHTIYCGGKKK | **4b** | 36% | **4c** | 11% |
| 3 | **5a** | SICRFFCGGG | **5b** | 59% | **5c** | 44% |
| | **6a** | SFCPMFCGGG | **6b** | 16% | **6c** | 37% |
| | **7a** | SLCKRECGGG | **7b** | 39% | **7c** | 28% |
| | **8a** | STCQGECGGG | **8b** | 67% | **8c** | 38% |

**Figure S9:** Summary of the selected peptide sequences nominated from three panning campaigns. The nominated peptides were chemically synthesized and modified with **DFS** and **PFS** for validation.

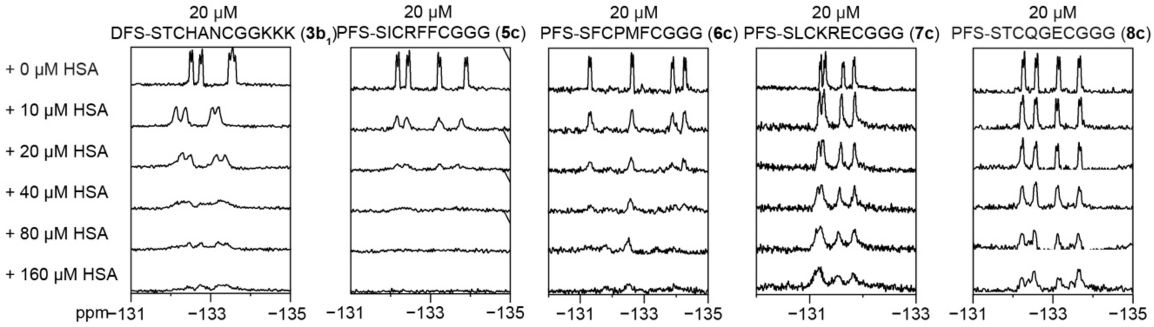

**Figure S10:** [19]F NMR binding assay for **3b** (PFS-STCHANCGKKK-DFS) **5c (**PFS-SICRFFCGGG), **6c** (PFS-SFCPMFCGGG), **7c** (PFS-SLCKRECGGG), and **8c** (PFS-STCQGECGGG) at 20 μM and titrated against various concentrations of HSA.

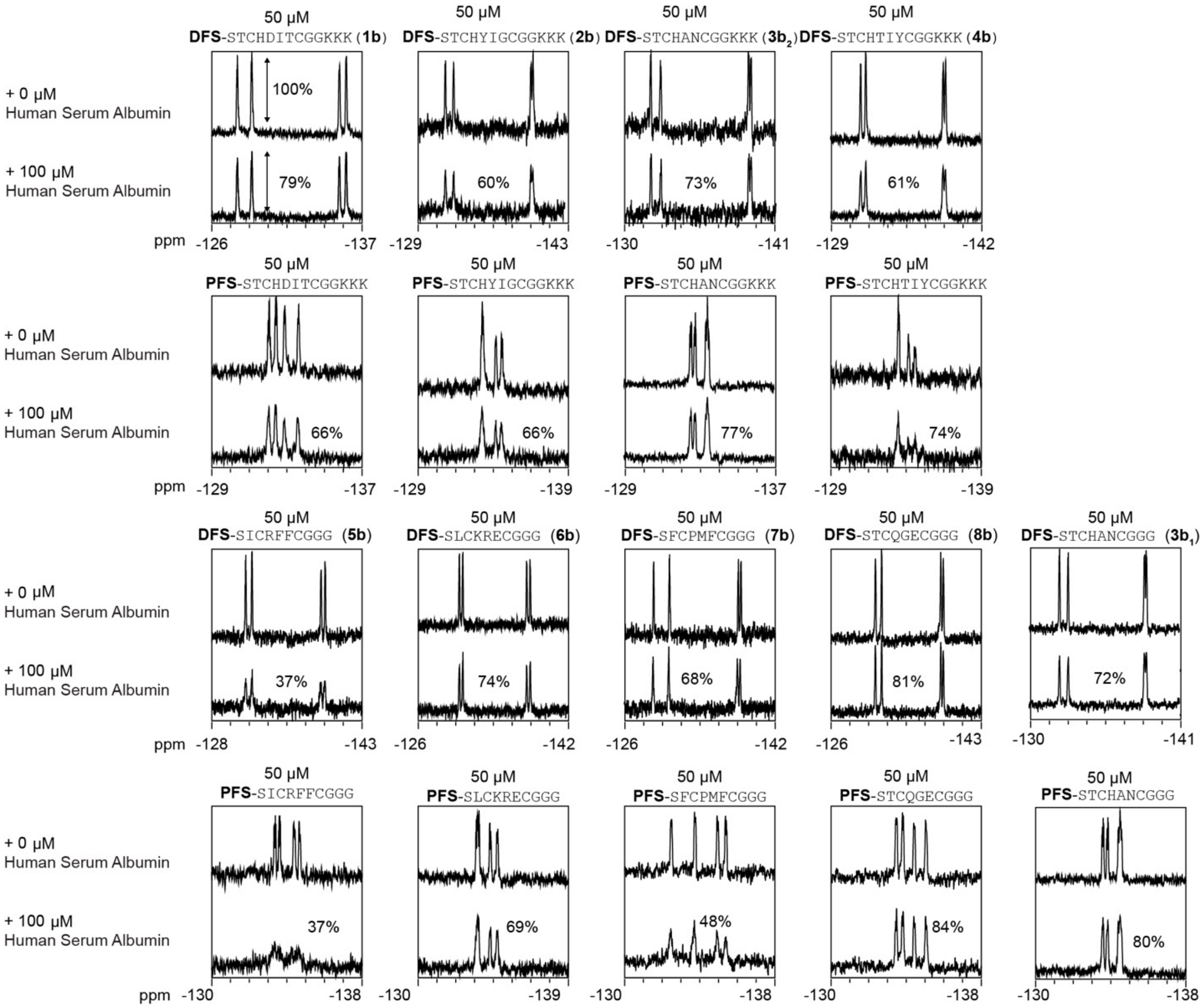

**Figure S11:** Summary of the $^{19}$F NMR binding measurement of the HSA titration spectra of 50 µM of **1b**-**8b** and **1c**-**8c** against 100 µM of HSA. The percentage represents the peak intensity remaining after the addition of HSA.

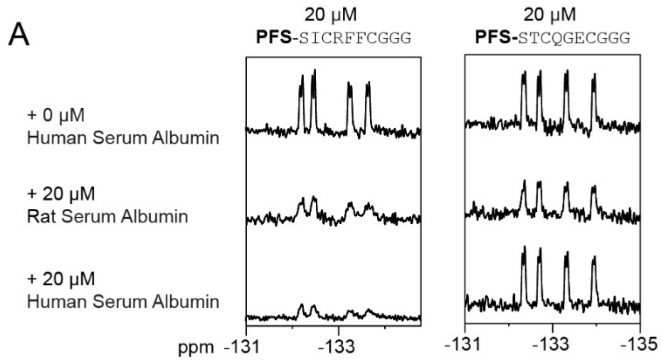

**Figure S12:** The $^{19}$F NMR titration spectra of **5c** and **8c** against rat serum albumin.

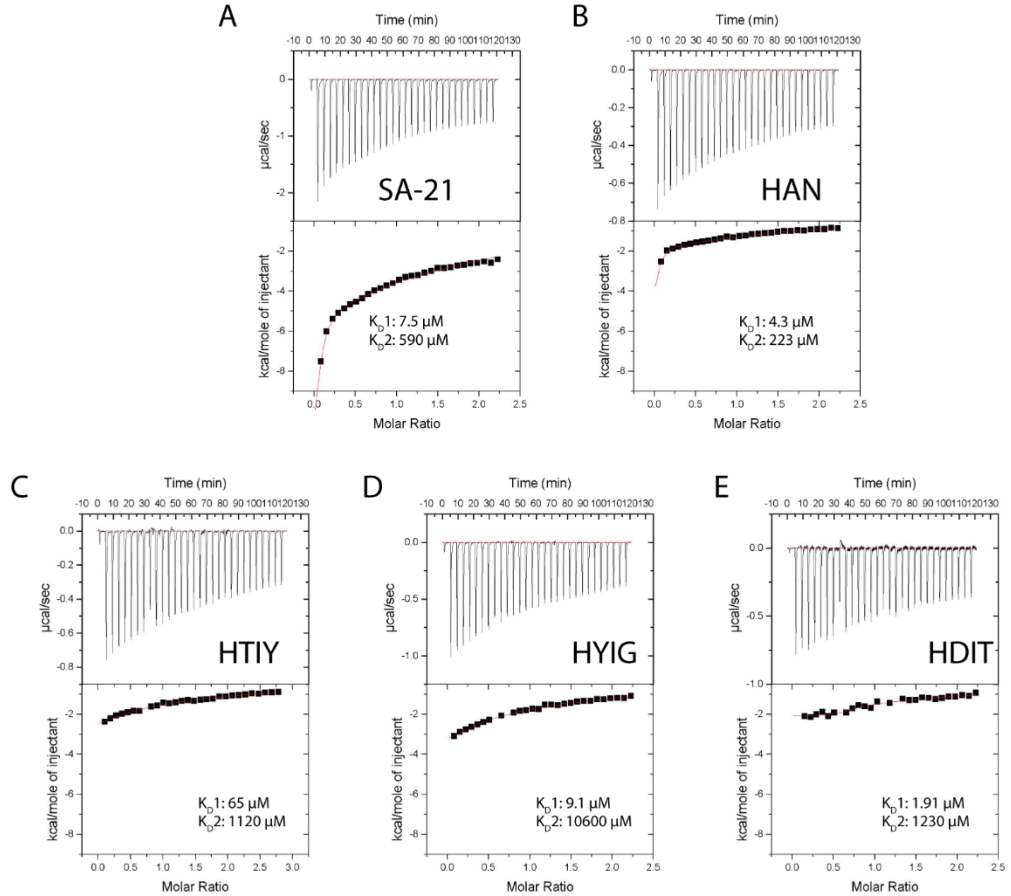

**Figure S13:** The first round of equilibrium dissociation constants of the selected peptides on HSA determined by ITC. The ITC traces and binding isotherms for peptides (A) SA-21 (4 mM), HSA (0.4 mM), (B) **DFS**-STCHANCGGKKK (4 mM), and HSA (0.4 mM), (C) **DFS**-STCHTIYCGGKKK (4 mM), HSA (0.4 mM), (D) **DFS**-STCHYIGCGGKKK (4mM), HSA (0.4 mM) and **DFS**-STCHDITCGGKKK (4 mM), HSA (0.4 mM) in 1×PBS.

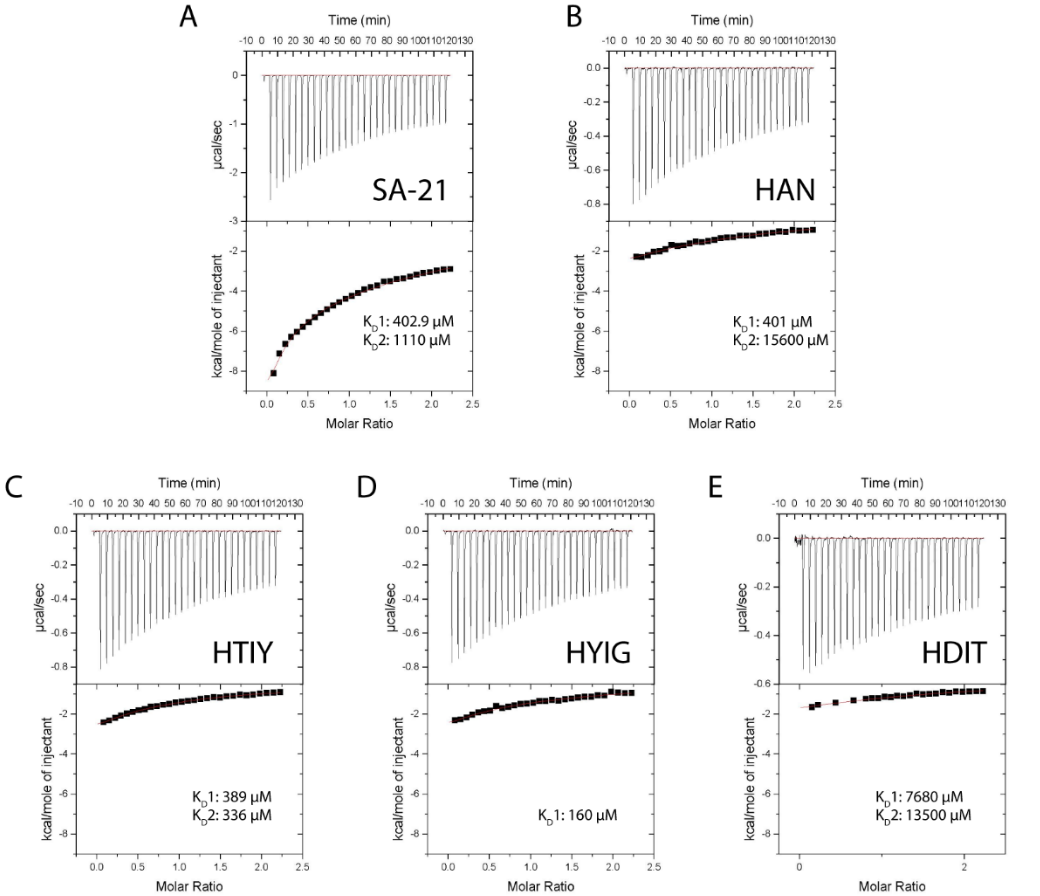

**Figure S14:** Second round of equilibrium dissociation constants of the selected peptides on HSA determined by ITC. The ITC traces and binding isotherms for peptides (A) SA-21 (1 mM), HSA (0.1 mM), (B) **DFS**-STCHANCGGKKK (1 mM), and HSA (0.1 mM), (C) **DFS**-STCHTIYCGGKKK (1 mM), HSA (0.1 mM), (D) **DFS**-STCHYIGCGGKKK (1mM), HSA (0.1 mM) and **DFS**-STCHDITCGGKKK (1mM), HSA (0.1 mM) in 1×PBS.

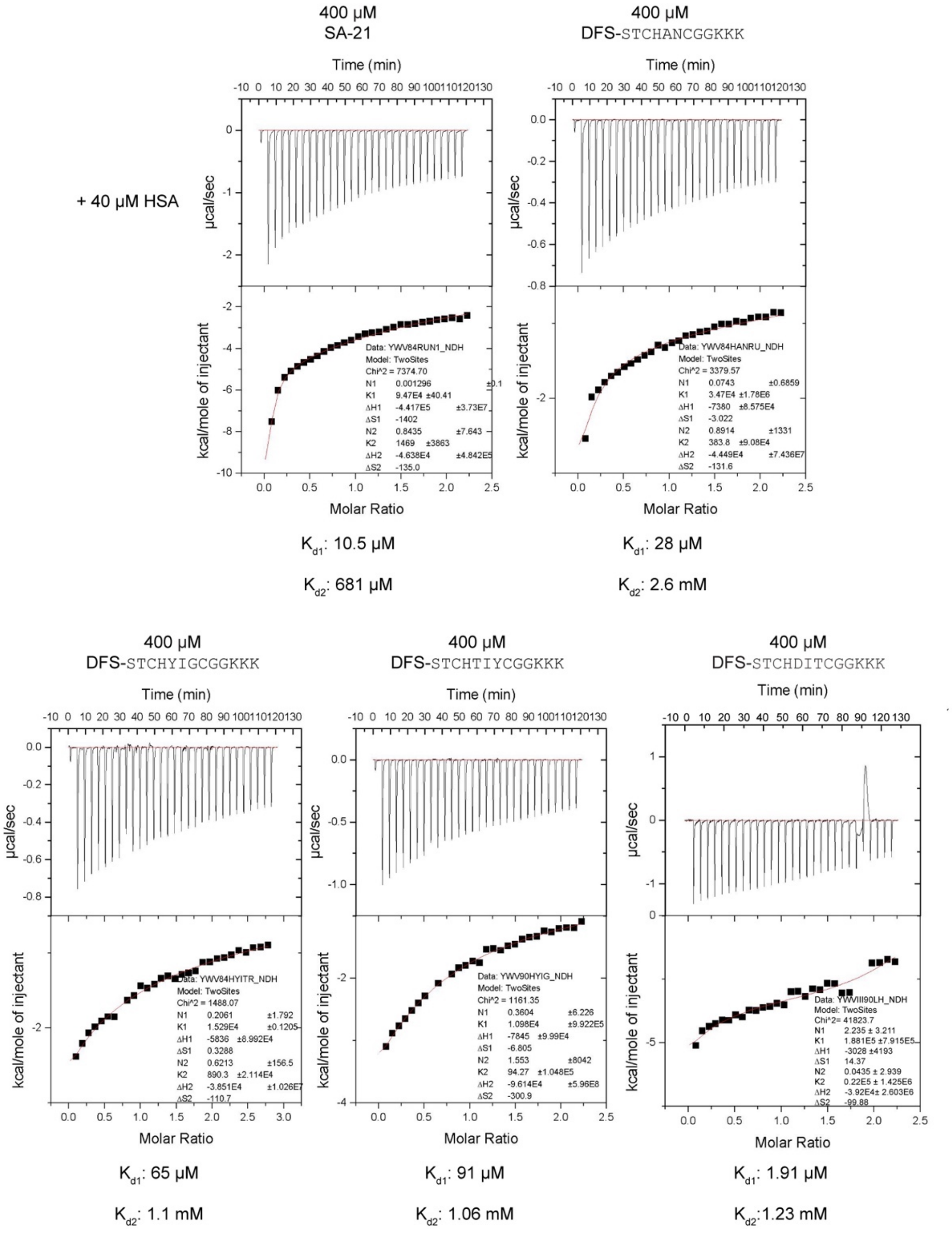

**Figure S15:** Summary of ITC binding assay SA-21 and **1b**-**5b** titrated against HSA.

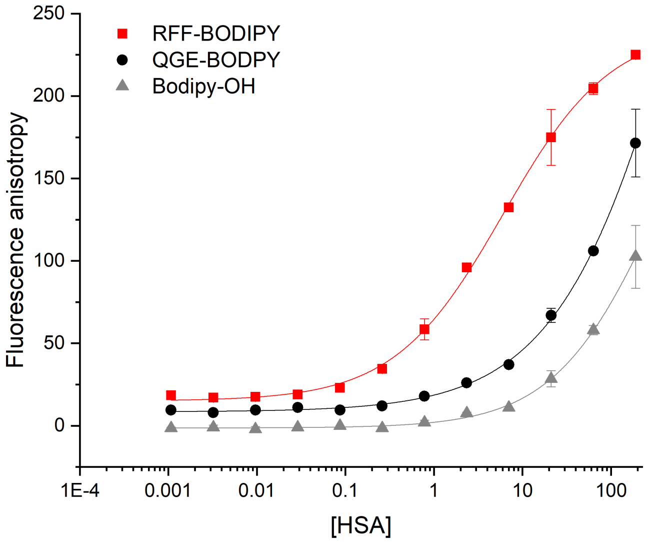

**Figure S16:** FP assay for BODIPY labeled **5d** titrated against **HSA 5d** found to have 6 µM binding affinity, **8d** (Black) was found to have >82 µM and BODIPY-OH (grey) found to have >320 µM binding affinity.

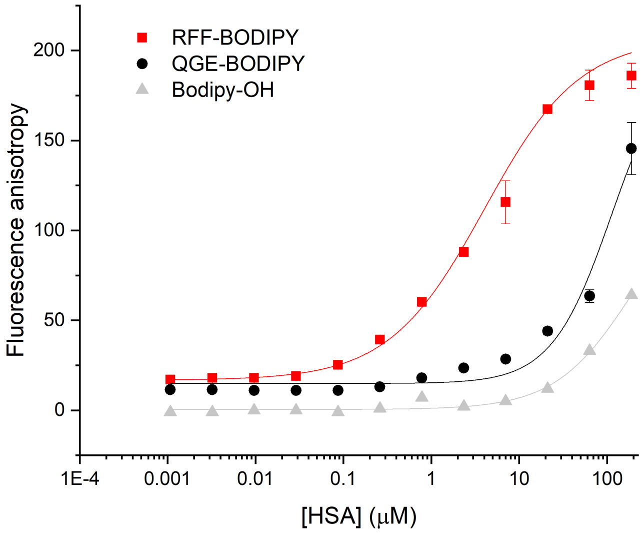

**Figure S17:** FP assay for BODIPY labeled **5d** (red) titrated against fatty acid-free HSA. **5d** was found to have 4 μM binding affinity, **8d** (Black) was found to have >112 μM and BODIPY-OH (grey) was found to have >196 μM binding affinity.

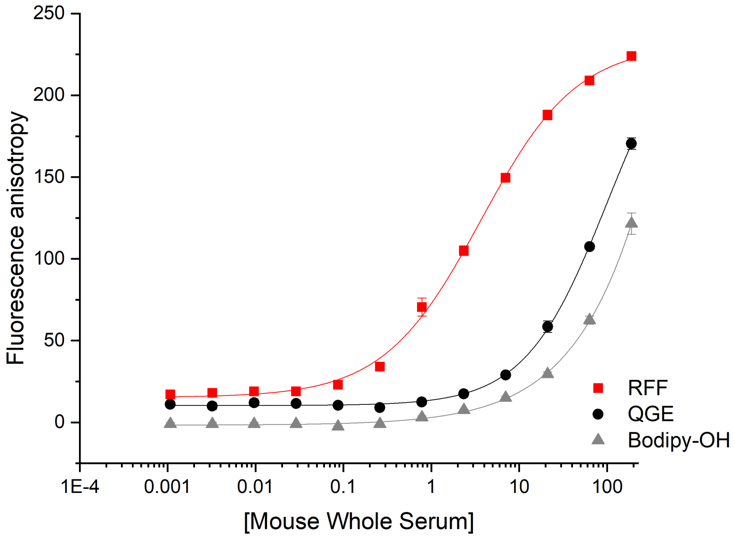

**Figure S18:** FP assay for Bodipy labeled RFF-**PFS**(red) titrated against whole mouse serum found to have 4 µM binding affinity, QGE-PGS(Black) found to have >100 µM and Bodipy-OH (grey) found to have >80 µM binding affinity.

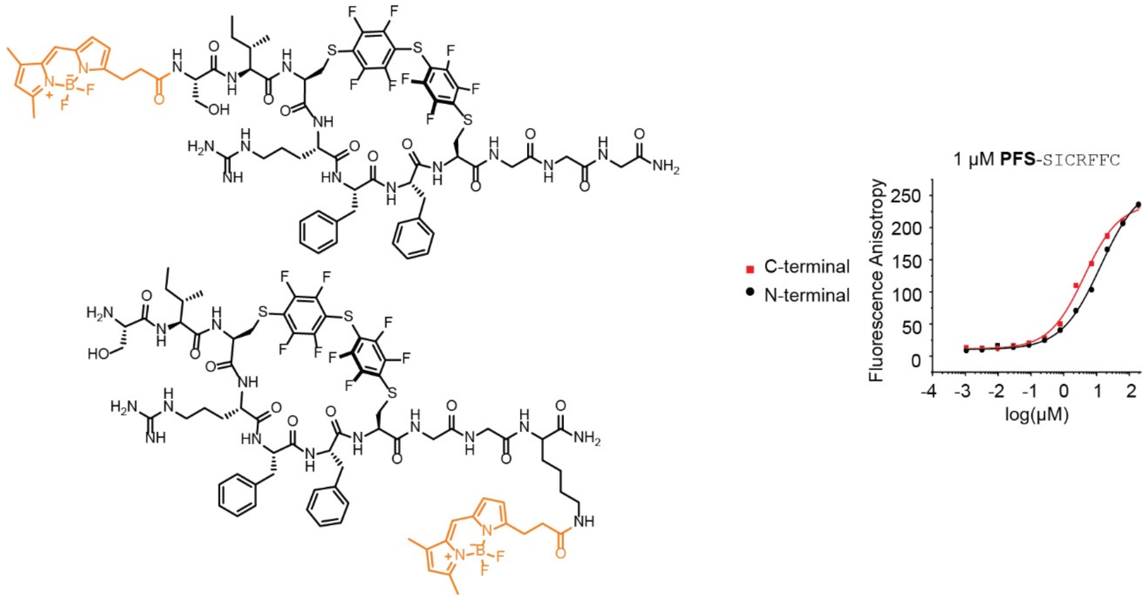

**Figure S19:** The FP binding assay of conjugate **5c** with BODIPY installed at N-terminus and C-terminus

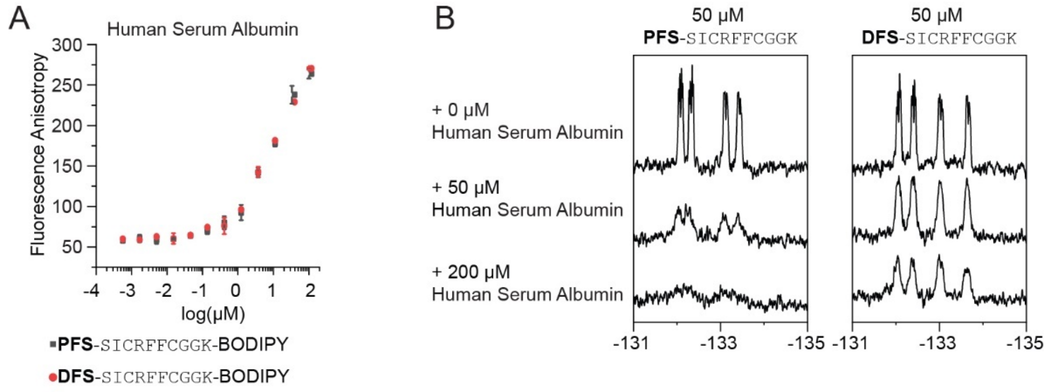

**Figure S20:** HSA binding between **5b** and **5c** (A) FP binding assay of **5b** and **5c**. (B) $^{19}$F NMR binding assay of **5b** and **5c**

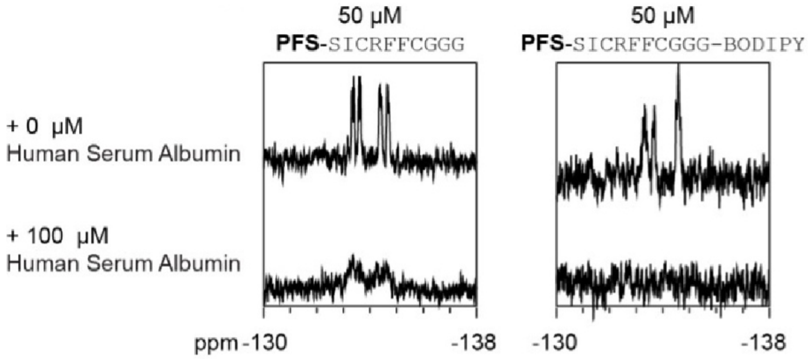

**Figure S21:** $^{19}$F NMR comparison of **PFS-**SICRFFCGGG (**5c)** and BODIPY labeled **5c**.

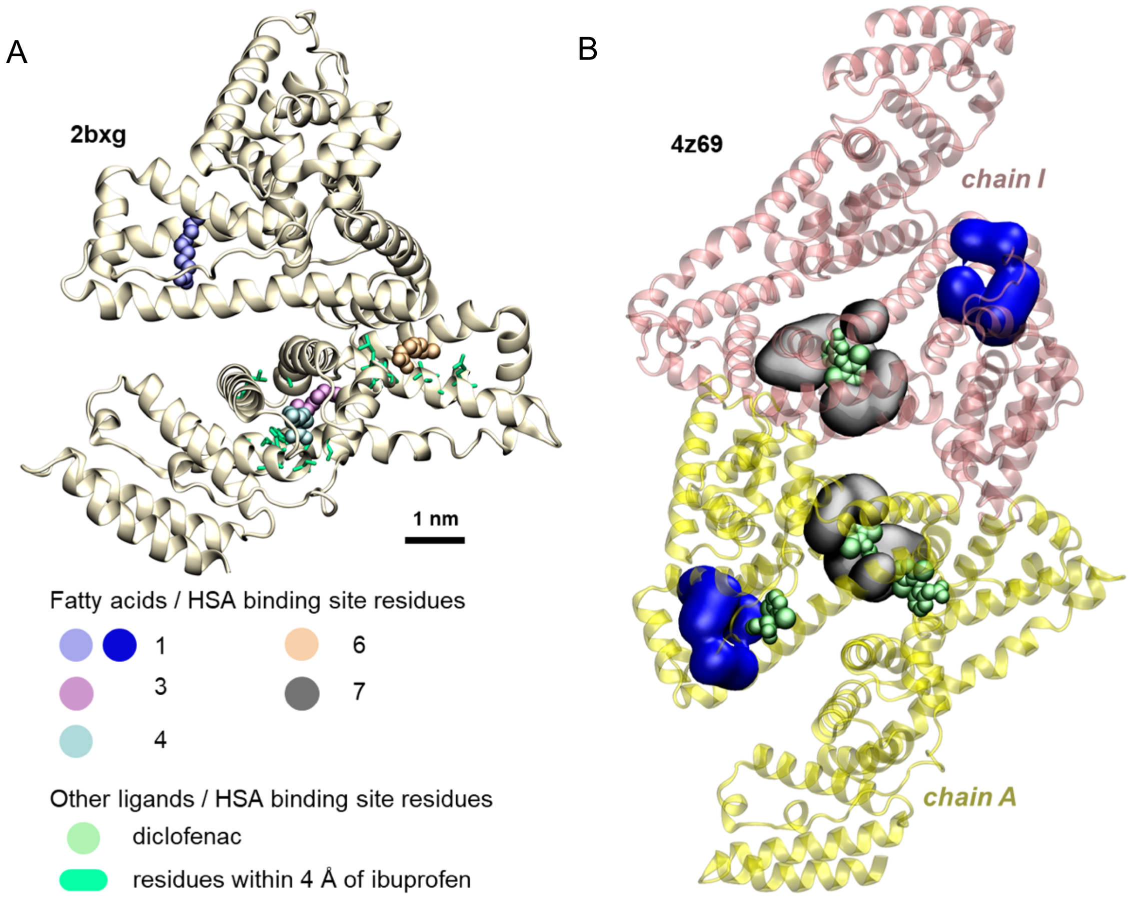

**Figure S22:** HSA binding sites of ibuprofen and diclofenac ligands in crystal structures. a) Structure of HSA in pdb file 2bxg. The atoms in green licorice representation show the HSA residues within 4 Å of the bound ibuprofen molecules, which bind near binding sites 3/4 and 6. Binding site 1, highlighted by the fatty acid shown in blue spheres, is distant from ibuprofen binding sites. **b)** Structure of two HSA chains and four diclofenac molecules in pdb file 4z69. Grey surfaces show HSA residues forming binding site 7, and blue surfaces show HSA residues forming

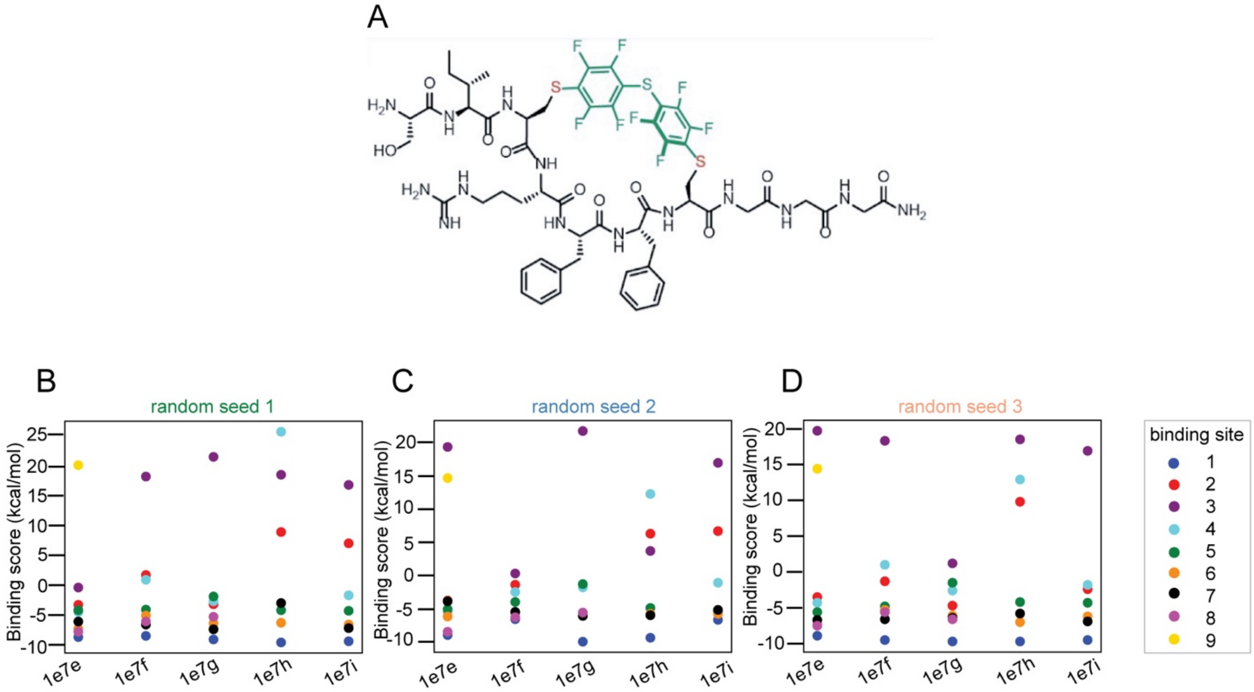

**Figure S23:** Binding scores of **PFS**-SICRFFCGGG macrocycle to HSA binding sites of fatty acids, determined in docking calculations. (A-C) Plots of **PFS**-SICRFFCGGG-HSA binding scores obtained in docking calculations using three different random seed values. Separate docking calculations were performed for different HSA structures extracted from PDB Databank files with PDB IDs 1e7e, 1e7f, 1e7g, 1e7h, and 1e7i. **PFS**-SICRFFCGGG macrocycles were docked to previously reported six to nine fatty acid binding sites on HSA surface[4], each labeled by different color. Some of the HSA fatty acid binding sites and the resulting RFF-HSA binding modes are shown in **Figure 6**. In each HSA structure, RFF was docked only to the sites occupied by the fatty acids in the pdb file. Therefore, docking was not tested for all the binding sites for every HSA structure, and some scores are missing from the plots.

**Table S5:** Binding scores of RFF macrocycle to HSA fatty acid binding sites from all docking calculations. In each HSA structure, the RFF was docked only to the sites occupied by fatty acids in the pdb file. Therefore, docking was not tested for all the binding site for every structure, and some scores are thus missing from the table.

| binding scores (kcal/mol) | | | | | | | | | | | | | | | | average | standard deviation |
|---|---|---|---|---|---|---|---|---|---|---|---|---|---|---|---|---|---|
| binding sites | random seed 1 | | | | | random seed 2 | | | | | random seed 3 | | | | | | |
| | 1e7e | 1e7f | 1e7g | 1e7h | 1e7i | 1e7e | 1e7f | 1e7g | 1e7h | 1e7i | 1e7e | 1e7f | 1e7g | 1e7h | 1e7i | | |
| 1 | -8.7 | -8.5 | -9.1 | -9.6 | -9.4 | -9 | -6.6 | -10 | -9.4 | -6.7 | -8.9 | -9.5 | -9.7 | -9.7 | -9.5 | -8.95 | 1.02 |
| 2 | -3.3 | 1.7 | -3.2 | 8.9 | 7 | -3.8 | -1.4 | -5.6 | 6.3 | 6.7 | -3.5 | -1.3 | -4.7 | 9.8 | -2.4 | 0.75 | 5.44 |
| 3 | -0.4 | 18.2 | 21.5 | 18.5 | 16.8 | 19.4 | 0.3 | 21.8 | 3.7 | 17 | 19.7 | 18.3 | 1.2 | 18.5 | 16.9 | 14.09 | 8.22 |
| 4 | -4.5 | 0.9 | -2.7 | 25.7 | -1.7 | -5.5 | -2.5 | -1.8 | 12.3 | -1.1 | -4.3 | 1 | -2.6 | 12.9 | -1.8 | 1.62 | 8.62 |
| 5 | -4.2 | -4.1 | -1.9 | -4.2 | -4.3 | -5.1 | -4 | -1.3 | -4.9 | -5.7 | -5.6 | -4.8 | -1.5 | -4.2 | -4.3 | -4.01 | 1.37 |
| 6 | -7.3 | -5.1 | -6.4 | -6.3 | -6.6 | -6.2 | -5.6 | -5.6 | -5.8 | -5.8 | -7.3 | -5.3 | -6.1 | -7 | -6.2 | -6.17 | 0.67 |
| 7 | -6.1 | -6.6 | -7.4 | -3 | -7.2 | -3.9 | -5.5 | -6.1 | -6 | -5.2 | -6.7 | -6.6 | -6.4 | -5.8 | -6.9 | -5.96 | 1.19 |
| 8 | -7.8 | -6.1 | -5.3 | - | - | -8.5 | -6.3 | -5.6 | - | - | -7.5 | -5.6 | -6.6 | - | - | -6.59 | 1.11 |
| 9 | 20.1 | - | - | - | - | 14.7 | - | - | - | - | 14.4 | - | - | - | - | 16.40 | 3.21 |

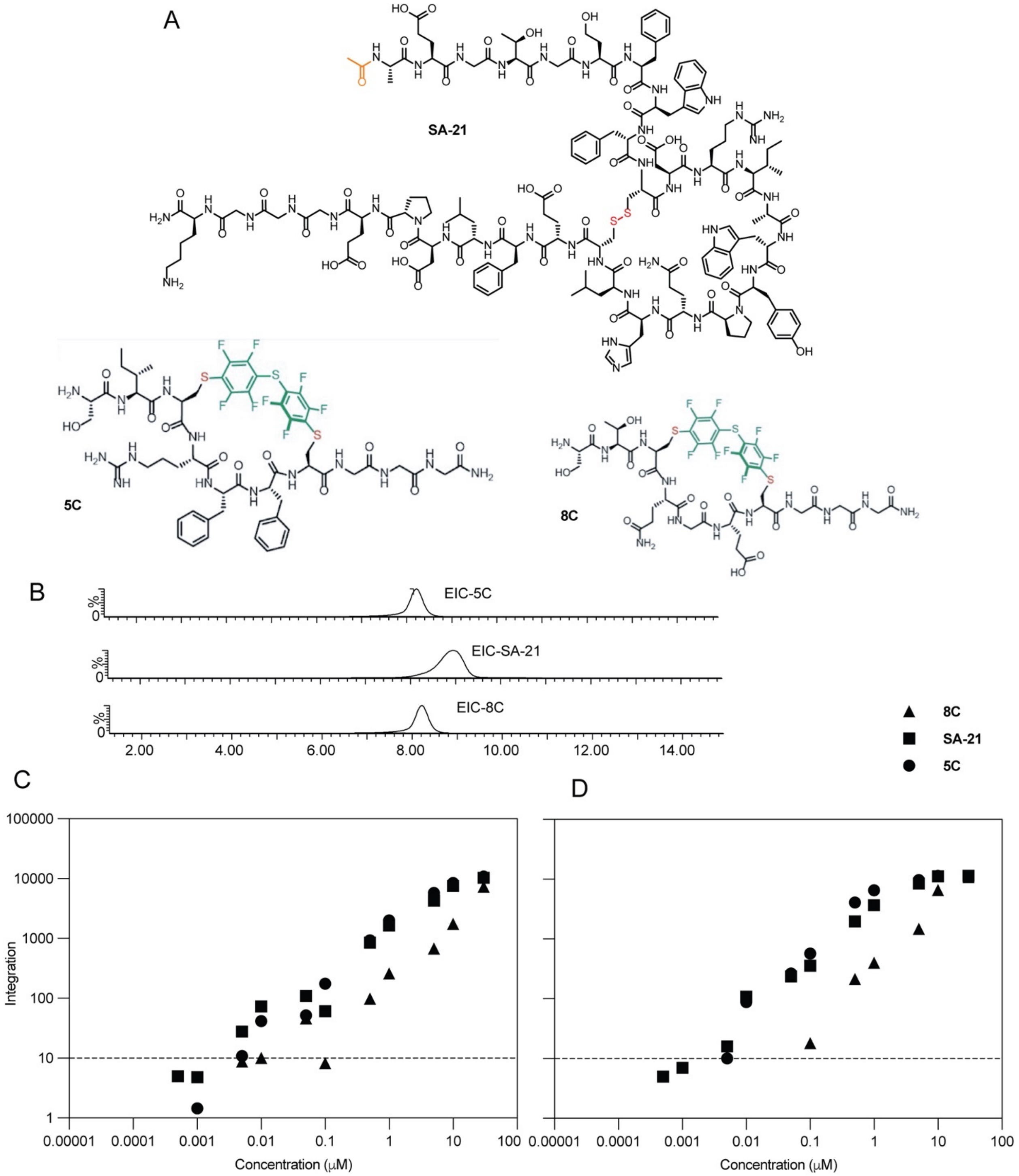

**Figure S24:** Detection of macrocycles using mass spectrometry. (A) Total ion chromatograms (TIC) of macrocycles **8c**, **SA21,** and **5c**. (B) Standard curve for macrocycles in PBS (C) Standard curve for macrocycles in mice serum

#### **19.** MATLAB script for DE analysis

```
clear;

Dir='';
File = 'YW_unfiltered_20170829_ed.txt';

SET⁶  = 1:3;        % HSA
SET⁷ = 4:9;          % T4-GP1
SET⁶  = 10:12;      % ConA
SET⁶  = 13:15;      % Input

TEST_SET = 1;
TARGET = 'HSA';
CONTROL_SETS = [2 3 4];

REREAD=1;   % change to zero if you don’t want to wait for -re-reading of data

close all

OUTPUT='normalized';   % other output_type: 'normalized' 'normalized+1' 'raw'

if REREAD
    disp('reading...');
    [Nuc, AA, Fr] = readMulticolumn('Dir', Dir, 'File', File, ...
                                'column', 1:max(cell2mat(SET)),...
                                'skip', 2, 'output', OUTPUT);
end
%%% To plot Figure S1D chage the variables above to the variables below
% TEST_SET = 9;
% CONTROL_SETS = [10 11];

HITS2DISPLAY = 50; % maximum numer of hits to display
SHOWaminoACIDS = [18 19 20 21 22 23 24 25 26 27];

CLUSTERbyH = 1;         % 1 if you want your hits to be clustered by Hamming dist.
PLOT_VOLCANO = 1;       % set to 1 if you want to see the actual volcano plot
Sort_CXC_file = 1;      % set to 1 if you want to sort the reslt into different CXC files
AA_AXA_Analysis = 1;    % set to 1 if you want to see the actual AA_AXA_Analysis

%%%%%%% volcano plot parameters here %%%%%%%%%%%%%%%%%%%%%%%%%%%%%%%%%%%%%%
p_cutoff = 0.05;             % p-value cutoff
R_cutoff = 3;               % ratio cutoff
MaxX=6;                     % maximum on the X-scale (if plotting volcano)
vert_cutoff = 0.00001;      % maximum on the Y-scale (if plotting volcano)
%%%%%%%%%%%%%%%%%%%%%%%%%%%%%%%%%%%%%%%%%%%%%%%%%%%%%%%%%%%%%%%%%%%%%%%%%%%

%%%%%%%%%%%%%%%%%%%%%% do not change things beyond this point %%%%%%%%%%%%%
%%%%%%%%%%%%%%%%%%%%%% unless you know what you are doing %%%%%%%%%%%%%%%%%

if CLUSTERbyH == 1
    disp('Culster on')
else
    disp('Culster off')
end

if PLOT_VOLCANO == 1
    disp('Volcano Plot on')
else
    disp('Volcano plot off')
end

if Sort_CXC_file == 1
    disp('Sort CXC on')
else
```

```
    disp('Sort CXC off')
end

if AA_AXA_Analysis == 1
    disp('AA AXA Analysis on')
else
    disp('AA AXA Analysis off')
end

SAVEto = [File(1:end-4) TARGET '_'...
                         OUTPUT...
                         'CONTR_' num2str(CONTROL_SETS) '_'...
                         '_P' num2str(p_cutoff)...
                         '_R' num2str(R_cutoff)...
                         '.CSV'];  % keep blank if don't want to save

% select only the aminoacids you want to see
cAA = char(AA);
sAA=cellstr(cAA(:,SHOWaminoACIDS));
SQUARE=zeros(size(Fr,1),1);

i=0;
disp('calculating p and R...');
IX=zeros(size(Fr,1),numel(CONTROL_SETS));

disp('calculating p and R...');
i=0;
for j=CONTROL_SETS
    i=i+1;
    ratio(:,i) = mean(Fr(:,SET{TEST_SET}), 2) ./ mean(Fr(:,SET{j}), 2);

    [~,confi(:,i)] = ttest2(Fr(:,SET{TEST_SET})',Fr(:,SET{j})',....
                            p_cutoff,'both','unequal');

    IX(:,i) = (confi(:,i) <= p_cutoff) & (ratio(:,i) >= R_cutoff);

    SQUARE = SQUARE + ratio(:,i).^2;

    if PLOT_VOLCANO
        subplot(1,numel(CONTROL_SETS),i);

        plot(log2 (ratio(:,i)),...
            -log10(confi(:,i)),'d',...
            'MarkerSize',4,...
            'MarkerFaceColor',0.5*[1 1 1],...
            'MarkerEdgeColor',0.5*[1 1 1]); hold on;

        plot( log2 (ratio(find(IX(:,i)),i)),...
             -log10(confi(find(IX(:,i)),i)),'d',...
                    'MarkerSize',4,...
                    'MarkerFaceColor','r',...
                    'MarkerEdgeColor','r'); hold on;

        line([log2(R_cutoff) MaxX],[-log10(p_cutoff) -log10(p_cutoff)]);
        line([log2(R_cutoff)   log2(R_cutoff)],...
             [-log10(p_cutoff) -log10(vert_cutoff)]);

        xlim([-MaxX MaxX]);
    end
end

    R2 = sqrt(SQUARE);

    IXall = find( (sum(IX,2)==size(IX,2)) ); % hits that satisfy all criteria
```

```
   % you can loosen the stringency if necessary
   % IXall = find( (sum(IX,2)>=2) ); %hits that satisfy two or more criteria

    hits    = char(sAA(IXall,:));
    Rhits   = ratio(IXall,:);
    R2hits  = R2(IXall);

%%%%%%%%%%%% this is part where hits are clustered by H-dist %%%%%%%%%%%%%%

if CLUSTERbyH
    disp('clustering...');
    if numel(hits)>3
        figure(2)
        Y = pdist(hits,'hamming');
        Z = linkage(Y,'complete');
        [H,T,perm] = dendrogram(Z,0,'colorthreshold',20);
        set(H,'LineWidth',2)

        for i =1:size(hits,1)
            label{i} = i;
        end
        set(gca,'XTick', 1:1:size(hits,1), 'XTickLabel',label);

        hits    = hits(perm,:);
        Rhits   = Rhits(perm,:);
        R2hits  = R2(perm);
        IXall   = IXall(perm);
    end
end

%%%%%%%%%%%%%%%%%%%display all results as heat map%%%%%%%%%%%%%%%%%%%%%%%%%

figure(3)

if size(IXall,1)>=HITS2DISPLAY
    N=HITS2DISPLAY;  % display only the first or defined number of hits
else
    N=size(IXall,1); %display all
end

FrPPM = round(10^6*Fr);   % convert normalized fraction frequency to PPM

imagesc( log10([FrPPM(IXall(1:N),:) ratio(IXall(1:N),:) ]+1) );

set(gca,'YTick', 1:1:N, 'YTickLabel',cellstr(hits(1:N,:)),'TickDir','out',...
    'FontName','Courier New','FontSize',14);
set(gca,'XTick', 1:1:size(Fr,2)+4, 'TickDir','out');
jet1=jet;
jet1(1,:)=[0.4 0.4 0.4];
colormap(jet1);
colorbar;

% generate a plain text table for saving or copy from command line
S = char(32*ones(size(hits,1),2));
COM = char(','*ones(size(hits,1),1));

L = [ S(:,1) char(124*ones(size(hits,1),1)) S(:,1)];
if strcmp(OUTPUT,'raw')
    F  = Fr(IXall,:);   % display frequency raw
else
    F  = FrPPM(IXall,:);   % display frequency in ppm
end

toDisp = [hits     S  ];
toSave = [hits     COM  ];

for i=1:numel(SET)
    for j=1:numel(SET{i})
```

```
        toDisp = [toDisp num2str(F(:,SET{i}(j))) S];
        toSave = [toSave num2str(F(:,SET{i}(j))) COM];
    end
    toDisp = [toDisp L];
end
toDisp = [toDisp S  num2str(round(Rhits)) L];
for i=1:size(Rhits,2)
    toSave = [toSave  num2str(round(Rhits(:,i))) COM ];
end

disp(toSave);
disp(toDisp);

if ~isempty(SAVEto)

    fs = fopen(fullfile(Dir,SAVEto),'w');
    RET = char(10*ones(size(toSave,1),1));
    fprintf( fs, '%s\r\n', [toSave(:,1:end-1) RET]');
    fclose all;
    disp('file saved');
end

%%%%%%%%%%%%%%%%%%%%%%%%%%%%%%%%C3C Sorting%%%%%%%%%%%%%%%%%%%%%%%%%%%%%%%%

if Sort_CXC_file

    disp('sorting...') ;

    %Sorting conditions
    C2X = ['S' '\w' 'C' '\w' '\w' 'C' '\w' '\w' '\w'] ;
    C3X = ['S' '\w' 'C' '\w' '\w' '\w' 'C' '\w' '\w' '\w'] ;
    C4X = ['S' '\w' 'C' '\w' '\w' '\w' '\w' 'C' '\w' '\w'] ;
    C5X = ['S' '\w' 'C' '\w' '\w' '\w' '\w' '\w' 'C' '\w'] ;
    C7X = ['A' 'C' '\w' '\w' '\w' '\w' '\w' '\w' '\w' 'C'] ;
    % Creat sorting array
    C2XHits = [];
    C3XHits = [];
    C4XHits = [];
    C5XHits = [];
    C7XHits = [];
    indexs=[];              %for data puposes not really useful for now
    j=0;

    % sort C2C
    disp('Sorting C2C...')
    C2s = regexp(cellstr(hits),C2X);
    for i=1:numel(C2s)
        if ~isempty(C2s{i})
            j=j+1;
            C2XHits = [C2XHits; hits(i,:)];
            indexs(j) = i;
        end
    end
    %C2Cfound(s) = numel(indexs);

    %Sort C3C
    disp('Sorting C3C...')
    indexs=[];
    j=0;
    C3s = regexp(cellstr(hits),C3X);
    for i=1:numel(C3s)
        if ~isempty(C3s{i})
            j=j+1;
            C3XHits = [C3XHits; hits(i,:)];
            indexs(j) = i;
        end
    end
    %C3Cfound(s) = numel(indexs);
```

```
    %Sort C4C
    disp('Sorting C4C...')
    indexs=[];
    j=0;
    C4s = regexp(cellstr(hits),C4X);
    for i=1:numel(C4s)
        if ~isempty(C4s{i})
            j=j+1;
            C4XHits = [C4XHits; hits(i,:)];
            indexs(j) = i;
        end
    end
    %C4Cfound(s) = numel(indexs);

    %Sort C5C
    disp('Sorting C5C...')
    indexs=[];
    j=0;
    C5s = regexp(cellstr(hits),C5X);
    for i=1:numel(C5s)
        if ~isempty(C5s{i})
            j=j+1;
            C5XHits = [C5XHits; hits(i,:)];
            indexs(j) = i;
        end
    end
    %C5Cfound(s) = numel(indexs);

    % Sort C7C
    disp('Sorting C7C...')
    indexs=[];
    j=0;
    C7s = regexp(cellstr(hits),C7X);
    for i=1:numel(C7s)
        if ~isempty(C7s{i})
            j=j+1;
            C7XHits = [C7XHits; hits(i,:)];
            indexs(j) = i;
        end
    end
    %C7Cfound(s) = numel(indexs);

end
if ~isempty(C2XHits)
    C2 = fopen(fullfile(Dir,['C2C',SAVEto]),'w');
    RET = char(10*ones(size(C2XHits,1),1));
    fprintf(C2, '%s\r\n',[C2XHits(:,1:end-1) RET]');
    fclose all;
    disp('C2C saved');
end

if ~isempty(C3XHits)
    C3 = fopen(fullfile(Dir,['C3C',SAVEto]),'w');
    RET = char(10*ones(size(C3XHits,1),1));
    fprintf(C2, '%s\r\n',[C3XHits(:,1:end-1) RET]');
    fclose all;
    disp('C3C saved');
end

if ~isempty(C4XHits)
    C4 = fopen(fullfile(Dir,['C4C',SAVEto]),'w');
    RET = char(10*ones(size(C4XHits,1),1));
    fprintf(C2, '%s\r\n',[C4XHits(:,1:end-1) RET]');
    fclose all;
    disp('C4C saved');
end

if ~isempty(C5XHits)
    C5 = fopen(fullfile(Dir,['C5C',SAVEto]),'w');
    RET = char(10*ones(size(C5XHits,1),1));
    fprintf(C2, '%s\r\n',[C5XHits(:,1:end-1) RET]');
```

```
    fclose all;
    disp('C5C saved');
end

if ~isempty(C7XHits)
    C7 = fopen(fullfile(Dir,['C7C',SAVEto]),'w');
    RET = char(10*ones(size(C7XHits,1),1));
    fprintf(C2, '%s\r\n',[C7XHits(:,1:end-1) RET]');
    fclose all;
    disp('C7C saved');

end

%%%%%%%%%%%%%%%%AA & AxA analysis%%%%%%%%%%%%%%%%%%%%%%%%%%%%%%%%%%%%%%%%%%%%%%

if AA_AXA_Analysis
    disp('Start AA AXA Analysis...')
    figure(100);
    AAA = 'ADEFHIKLMNPQRSTVWY';
    Y = [];

    for i=1:numel(AAA)
        Y(i) = numel(find(hits==AAA(i)));
        xlabel{i} = AAA(i);
    end

    plot(1:numel(AAA), Y, 'ok');
    set(gca, 'xTick', 1:numel(AAA), 'xTickLabel', xlabel, 'TickDir','out');

    %%

    Nfound =  [];
    NfoundS = [];
    toSaveIX = [];
    toSaveIXS = [];

    M = 9;

    fs = fopen(fullfile(Dir,['AA' SAVEto]),'w');
    fsS = fopen(fullfile(Dir,['AxA' SAVEto]),'w');
    fclose all;

    fs = fopen(fullfile(Dir,['AA' SAVEto]),'a+');
    fsS = fopen(fullfile(Dir,['AxA' SAVEto]),'a+');

    for ii = 1:numel(AAA)
        %disp(num2str(ii));
        for jj= 1:numel(AAA)

            phrase = [ AAA(ii) AAA(jj) ] ;
            phraseS = [ AAA(ii) '\w' AAA(jj) ] ;

            phraseHits = [];
            spacedHits = [];
            index=[];
            j=0;

            IX = regexp(cellstr(hits),phrase);
            for i=1:numel(IX)
                if ~isempty(IX{i})
                    %check whether his is S**** or A****; if it is, discard
                    if (phrase(1) =='S' || phrase(1) =='A')
                        if (numel(IX{i})==1 && IX{i}==1)
                            continue
```

```
                    end
                end

                j=j+1;
                phraseHits = [phraseHits; hits(i,:)];
                index(j) = i;

                S1 = char (32*ones(1, M-IX{i}(1)));
                S2 = char (32*ones(1, IX{i}(1)  ));
                spacedHits = [spacedHits; S1 hits(i,:) S2];
            end
        end
        Nfound(ii,jj) = numel(index);

        % lets save this with offsets
        RET = char(10*ones(size(index,2),1));
        fprintf( fs, '%s\r\n', [spacedHits toSave(index,:) RET]');

        phraseHits = [];
        index=[];
        spacedHits = [];
        j=0;
        clear IX

        IX = regexp(cellstr(hits),phraseS);
        for i=1:numel(IX)
            if ~isempty(IX{i})
                %check whether his is S**** or A****; if it is, discard
                if (phraseS(1) =='S' || phraseS(1) =='A')
                    if (numel(IX{i})==1 && IX{i}==1)
                        continue
                    end
                end

                j=j+1;
                phraseHits = [phraseHits; hits(i,:)];
                index(j) = i;

                S1 = char (32*ones(1, M-IX{i}(1)));
                S2 = char (32*ones(1, IX{i}(1)  ));
                spacedHits = [spacedHits; S1 hits(i,:) S2];
            end
        end
        NfoundS(ii,jj) = numel(index);
        toSaveIXS = [toSaveIXS index];

        % lets save this with offsets
        RET = char(10*ones(size(index,2),1));
        fprintf( fsS, '%s\r\n', [spacedHits toSave(index,:) RET]');

    end
end

figure(200);
subplot(1,2,1);
imagesc(Nfound); colorbar;
set(gca, 'xTick', 1:numel(AAA), 'xTickLabel', xlabel, 'TickDir','out',...
    'yTick', 1:numel(AAA), 'yTickLabel', xlabel);

subplot(1,2,2);
imagesc(NfoundS); colorbar;
set(gca, 'xTick', 1:numel(AAA), 'xTickLabel', xlabel, 'TickDir','out',...
    'yTick', 1:numel(AAA), 'yTickLabel', xlabel);

fclose all;
fs = fopen(fullfile(Dir,['AA' SAVEto]),'w');
fsS = fopen(fullfile(Dir,['AxA' SAVEto]),'w');
RET = char(10*ones(size(toSaveIX,2),1));
fprintf( fs, '%s\r\n', [toSave(toSaveIX,:) RET]');
disp('AA Saved')
```

```
    RET = char(10*ones(size(toSaveIXS,2),1));
    fprintf( fsS, '%s\r\n', [toSave(toSaveIXS,:) RET]');
    disp('AxA Saved')
end
```

**20.** Summary of synthesis

#### STCHDITCGGKKK-DFS

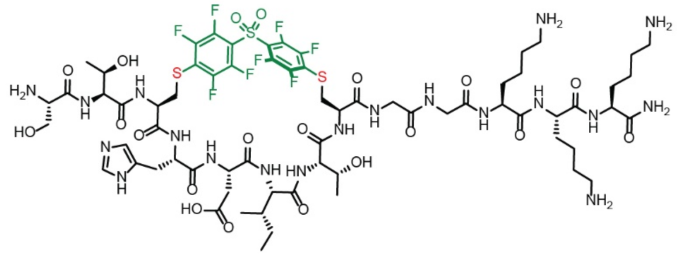

Chemical Formula: $C_{67}H_{95}F_8N_{19}O_{20}S_3$
Exact Mass: 1733.60
Molecular Weight: 1734.78

Starting material mass: 8.8 mg
Final product mass: 0.5 mg

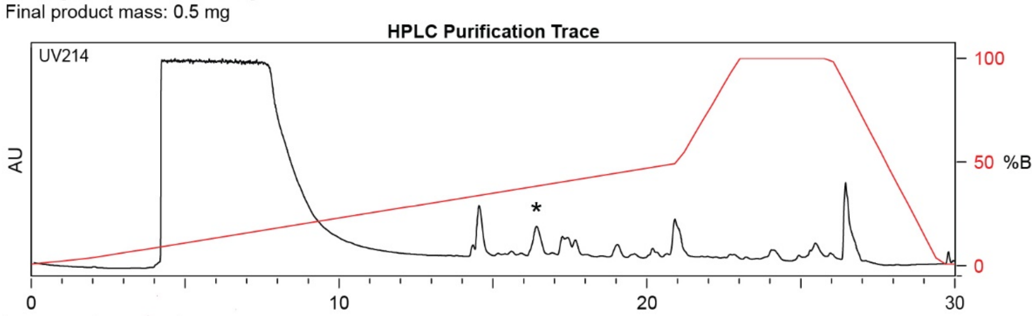

| Time (min) | Solvent B (%) |
|---|---|
| 0 | 2 |
| 5 | 30 |
| 24 | 95 |
| 25 | 100 |
| 27 | 100 |
| 28 | 2 |
| 30 | 2 |

Flow Rate: 13 mL / min
Phenomenex Kinetex EVO C18 Prep Column
(100 Å, 5 µm, 21.5 mm X 250 mm)

Slovent A: $H_2O$ + 0.1% (v/v) TFA
Slovent B: MeCN + 0.1% (v/v) TFA

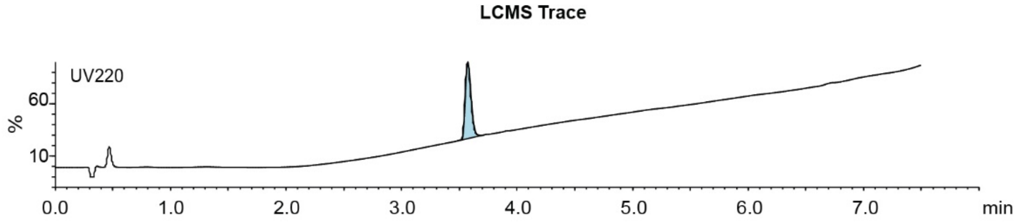

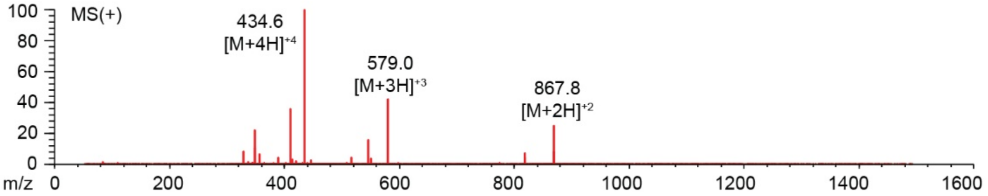

**Figure S25:** Synthesis summary of **1b**

#### STCHDITCGGKKK-PFS

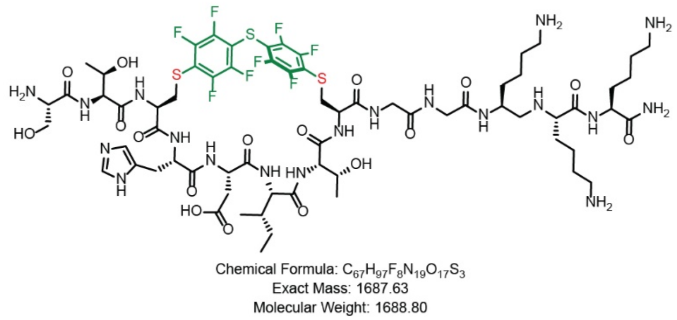

Starting material mass: 9.2 mg
Final product mass: 3.3 mg

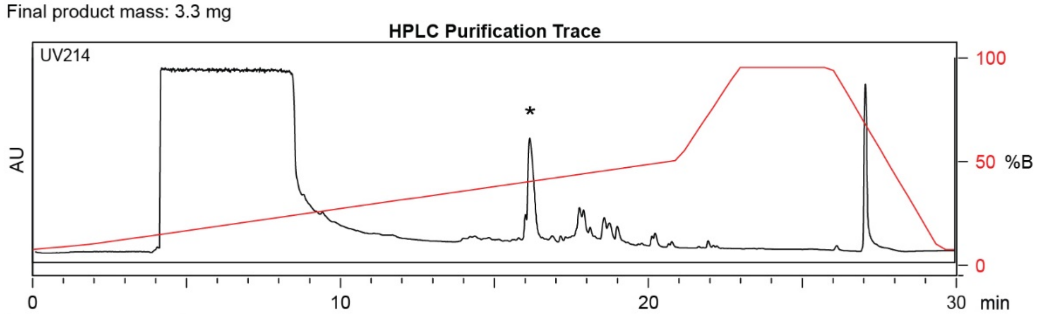

| Time (min) | Solvent B (%) |
|---|---|
| 0 | 2 |
| 2 | 5 |
| 21 | 50 |
| 23 | 100 |
| 26 | 100 |
| 29.5 | 2 |
| 30 | 2 |

Flow Rate: 13 mL / min
Phenomenex Kinetex EVO C18 Prep Column
(100 Å, 5 µm, 21.5 mm X 250 mm)

Solvent A: $H_2O$ + 0.1% (v/v) TFA
Solvent B: MeCN + 0.1% (v/v) TFA

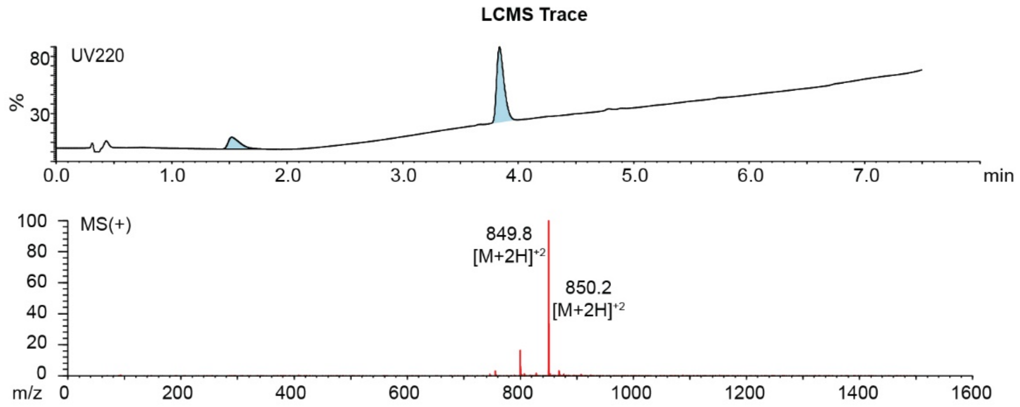

**Figure S26:** Synthesis summary of **1c.**

#### STCHTIYCGGG-PFS

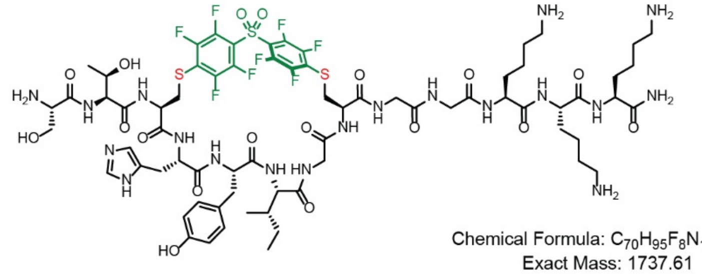

Chemical Formula: $C_{70}H_{95}F_8N_{19}O_{18}S_3$
Exact Mass: 1737.61
Molecular Weight: 1738.81

Starting material mass: 20 mg
Final product mass: 15 mg

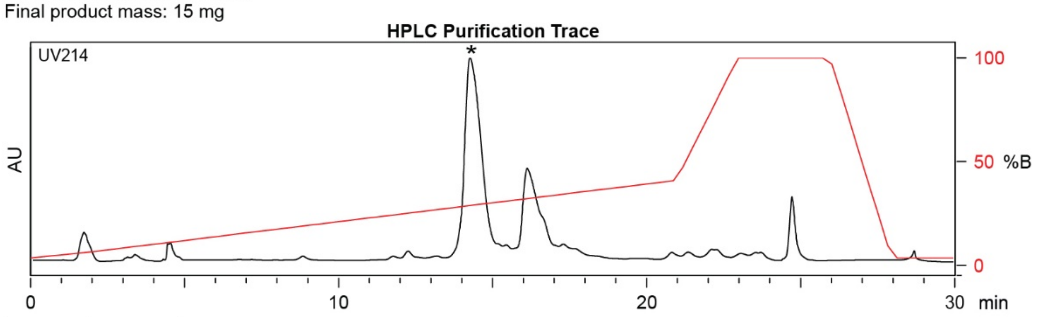

| Time (min) | Solvent B (%) |
|---|---|
| 0 | 2 |
| 2 | 5 |
| 21 | 50 |
| 23 | 100 |
| 26 | 100 |
| 29.5 | 2 |
| 30 | 2 |

Flow Rate: 13 mL / min
Phenomenex Kinetex EVO C18 Prep Column
(100 Å, 5 µm, 21.5 mm X 250 mm)

Solvent A: $H_2O$ + 0.1% (v/v) TFA
Solvent B: MeCN + 0.1% (v/v) TFA

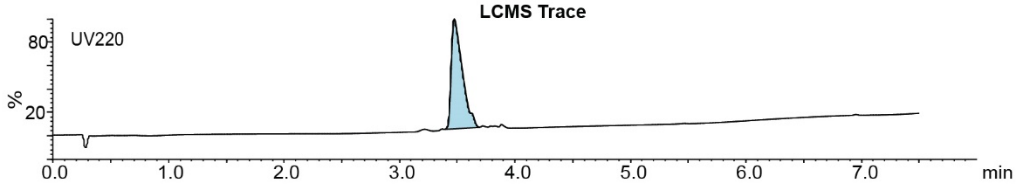

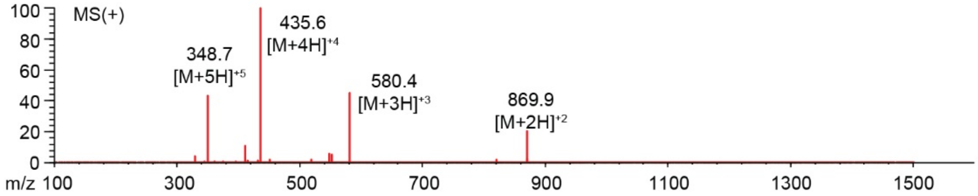

**Figure S27:** Synthesis summary of **2b**.

#### STCHYIGCGGKKK-PFS

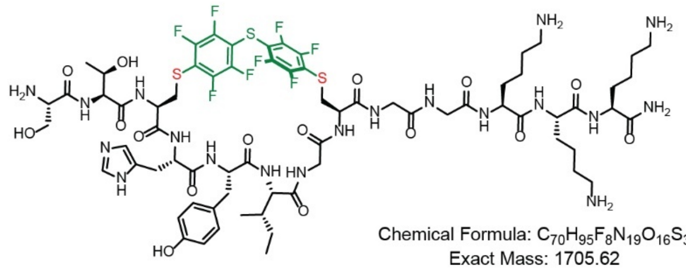

Chemical Formula: $C_{70}H_{95}F_{8}N_{19}O_{16}S_{3}$
Exact Mass: 1705.62
Molecular Weight: 1706.81

Starting material mass: 10 mg
Final product mass: 3.4 mg

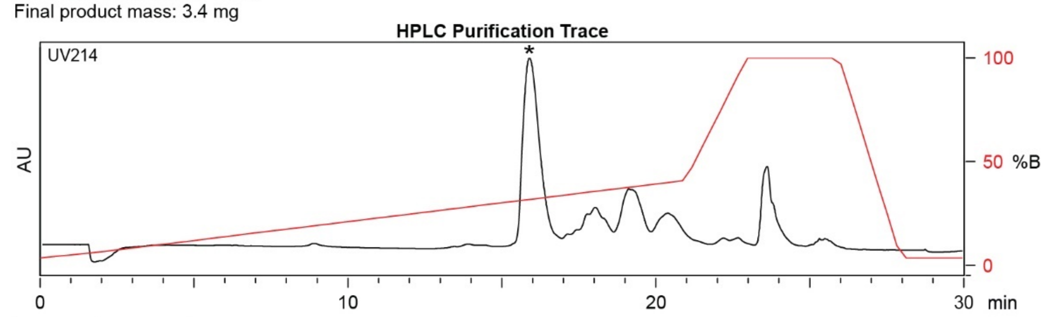

| Time (min) | Solvent B (%) |
|---|---|
| 0 | 2 |
| 2 | 5 |
| 21 | 50 |
| 23 | 100 |
| 26 | 100 |
| 29.5 | 2 |
| 30 | 2 |

Flow Rate: 13 mL / min
Phenomenex Kinetex EVO C18 Prep Column
(100 Å, 5 µm, 21.5 mm X 250 mm)

Solvent A: $H_2O$ + 0.1% (v/v) TFA
Solvent B: MeCN + 0.1% (v/v) TFA

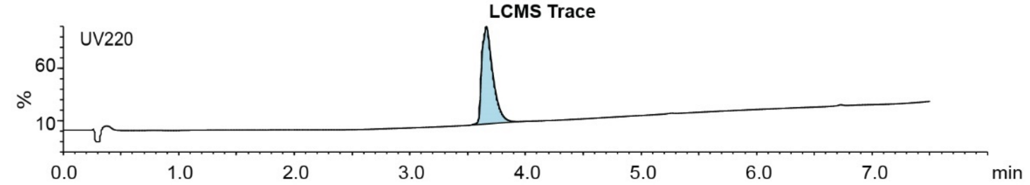

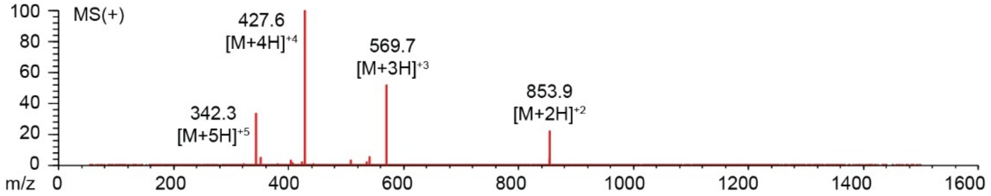

**Figure S28:** Synthesis summary of **2c**.

#### STCHANCGGKKK-DFS

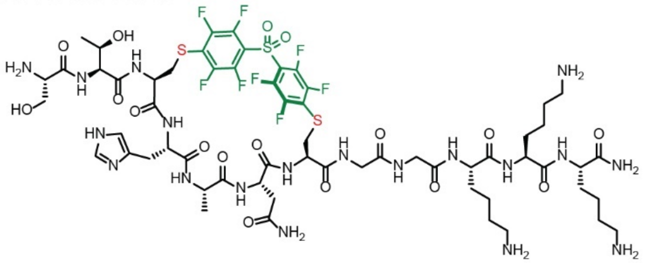

Chemical Formula: $C_{60}H_{83}F_8N_{19}O_{17}S_3$
Exact Mass: 1589.52
Molecular Weight: 1590.61

Starting material mass: 10.5 mg
Final product mass: 3.3 mg

**HPLC Purification Trace**

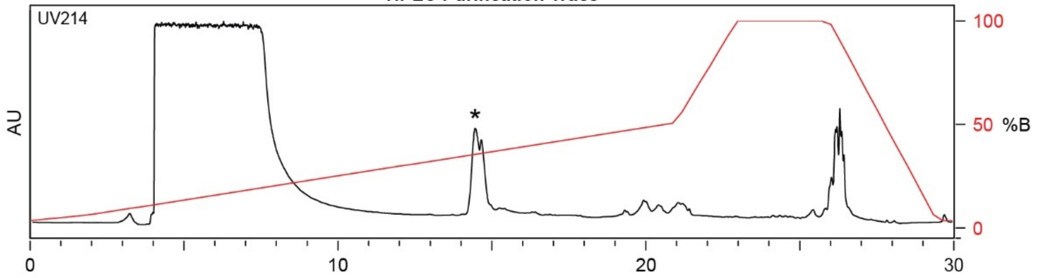

| Time (min) | Solvent B (%) |
|---|---|
| 0 | 2 |
| 2 | 5 |
| 21 | 50 |
| 23 | 100 |
| 26 | 100 |
| 29.5 | 2 |
| 30 | 2 |

Flow Rate: 13 mL / min
Phenomenex Kinetex EVO C18 Prep Column
(100 Å, 5 µm, 21.5 mm X 250 mm)

Slovent A: $H_2O$ + 0.1% (v/v) TFA
Slovent B: MeCN + 0.1% (v/v) TFA

**LCMS Trace**

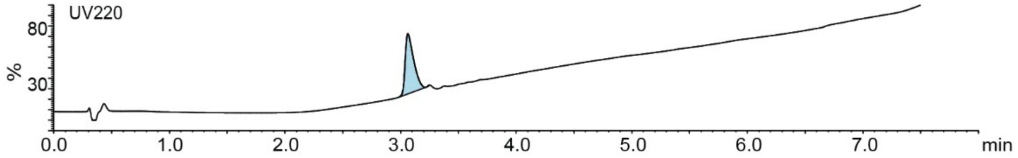

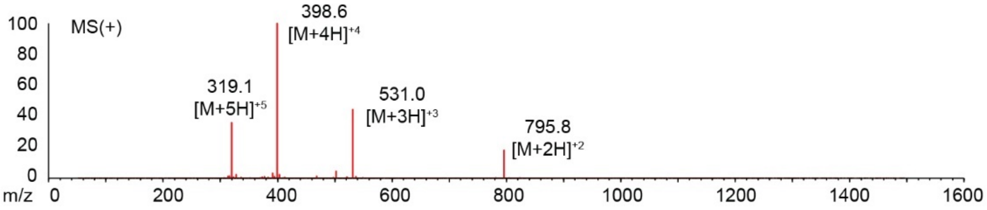

**Figure S29:** Synthesis summary of **$3b_1$**.

#### STCHANCGGKKK-PFS

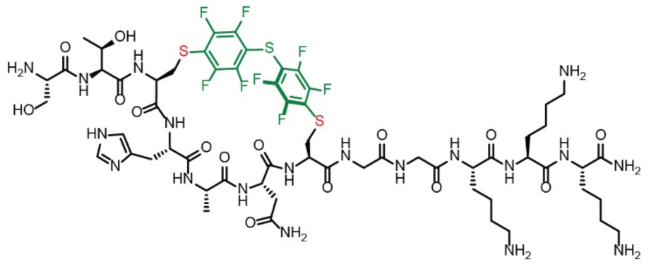

Chemical Formula: $C_{60}H_{83}F_8N_{19}O_{15}S_3$
Exact Mass: 1557.54
Molecular Weight: 1558.61

Starting material mass: 11.4 mg
Final product mass: 2.0 mg

**HPLC Purification Trace**

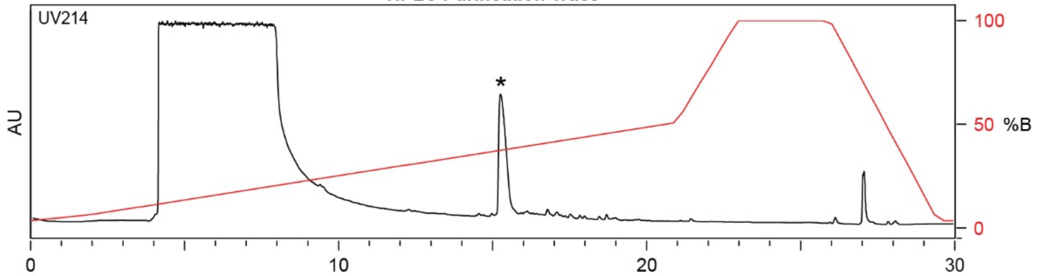

| Time (min) | Solvent B (%) |
|---|---|
| 0 | 2 |
| 2 | 5 |
| 21 | 50 |
| 23 | 100 |
| 26 | 100 |
| 29.5 | 2 |
| 30 | 2 |

Flow Rate: 13 mL / min
Phenomenex Kinetex EVO C18 Prep Column
(100 Å, 5 µm, 21.5 mm X 250 mm)

Solvent A: $H_2O$ + 0.1% (v/v) TFA
Solvent B: MeCN + 0.1% (v/v) TFA

**LCMS Trace**

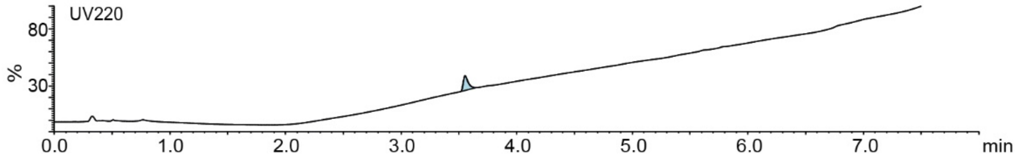

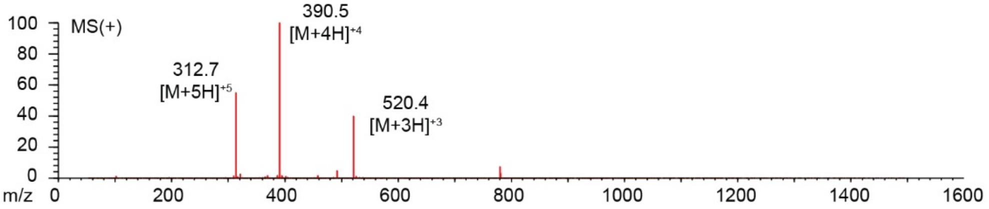

**Figure S30:** Synthesis summary of **3c**$_1$.

#### STCHANCGGG-PFS

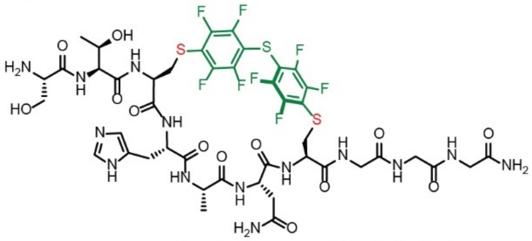

Chemical Formula: $C_{44}H_{50}F_8N_{14}O_{13}S_3$
Exact Mass: 1230.27
Molecular Weight: 1231.14

Starting material mass: 8.8mg
Final product mass: 3.3 mg

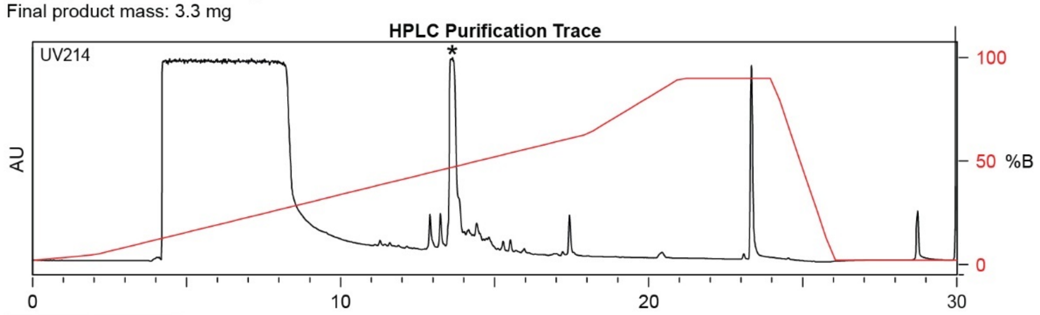

| Time (min) | Solvent B (%) |
|---|---|
| 0 | 2 |
| 2 | 5 |
| 18 | 70 |
| 21 | 100 |
| 24 | 100 |
| 26 | 2 |
| 30 | 2 |

Flow Rate: 13 mL / min
Phenomenex Kinetex EVO C18 Prep Column
(100 Å, 5 µm, 21.5 mm X 250 mm)

Solvent A: $H_2O$ + 0.1% (v/v) TFA
Solvent B: MeCN + 0.1% (v/v) TFA

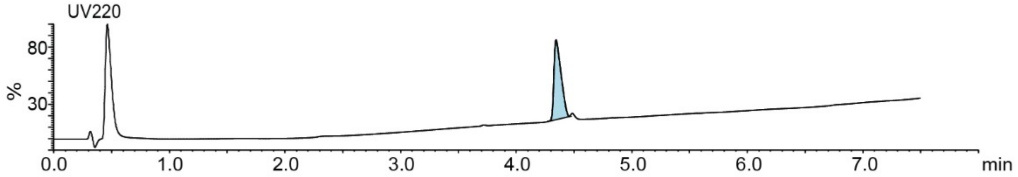

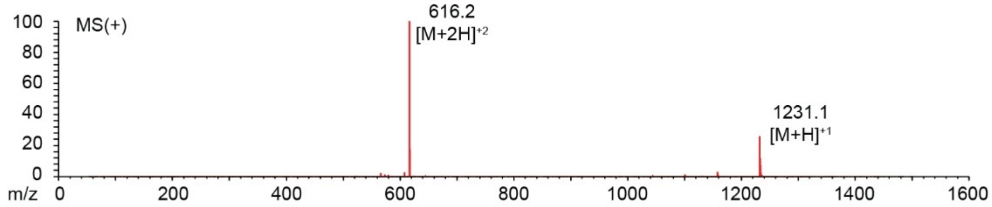

**Figure S31:** Synthesis summary of **3c₂**.

#### STCHTIYGGKKK-DFS

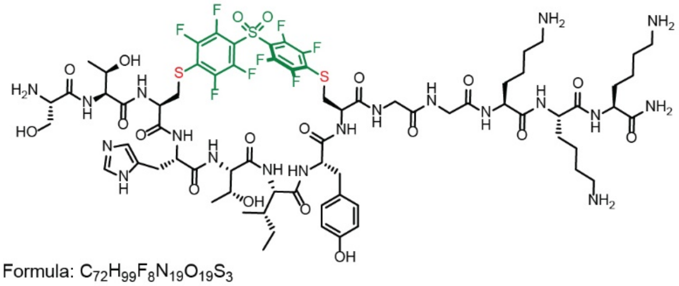

Chemical Formula: $C_{72}H_{99}F_8N_{19}O_{19}S_3$
Exact Mass: 1781.64
Molecular Weight: 1782.87

Starting material mass: 9.0 mg
Final product mass: 1.0 mg

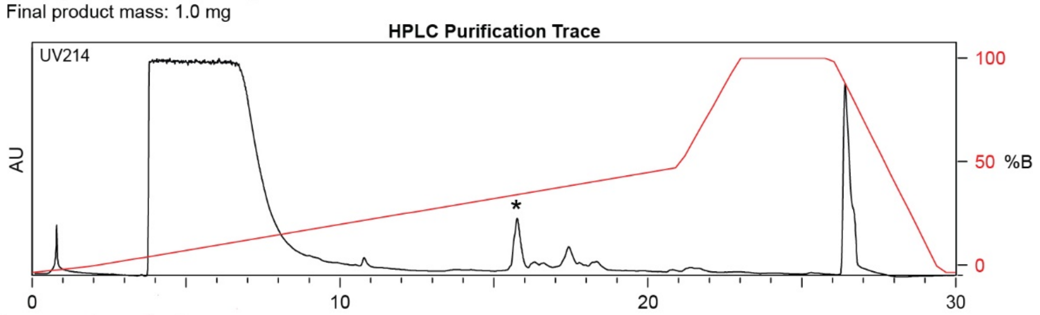

| Time (min) | Solvent B (%) |
|---|---|
| 0 | 2 |
| 2 | 5 |
| 21 | 50 |
| 25 | 100 |
| 27 | 100 |
| 28 | 2 |
| 30 | 2 |

Flow Rate: 13 mL / min
Phenomenex Kinetex EVO C18 Prep Column
(100 Å, 5 μm, 21.5 mm X 250 mm)

Slovent A: $H_2O$ + 0.1% (v/v) TFA
Slovent B: MeCN + 0.1% (v/v) TFA

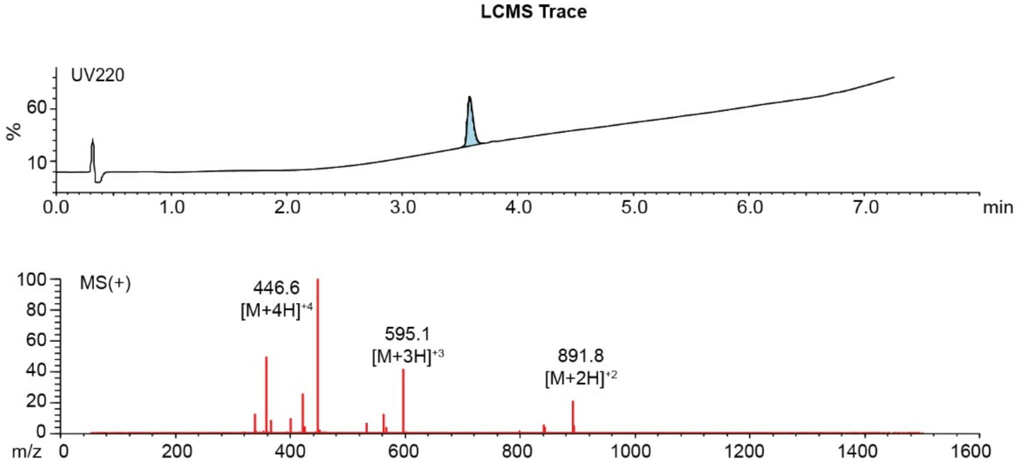

**Figure S32:** Synthesis summary of **4b**.

#### STCHTIYCGGG-PFS

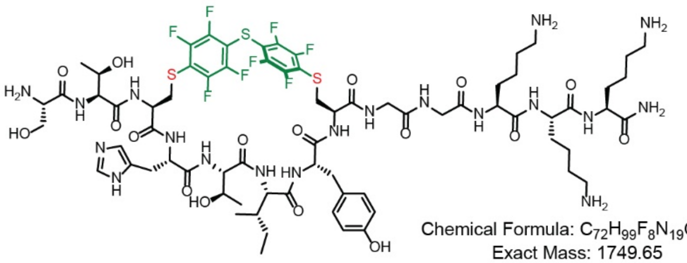

Chemical Formula: $C_{72}H_{99}F_8N_{19}O_{17}S_3$
Exact Mass: 1749.65
Molecular Weight: 1750.87

Starting material mass: 10 mg
Final product mass: 1.4 mg

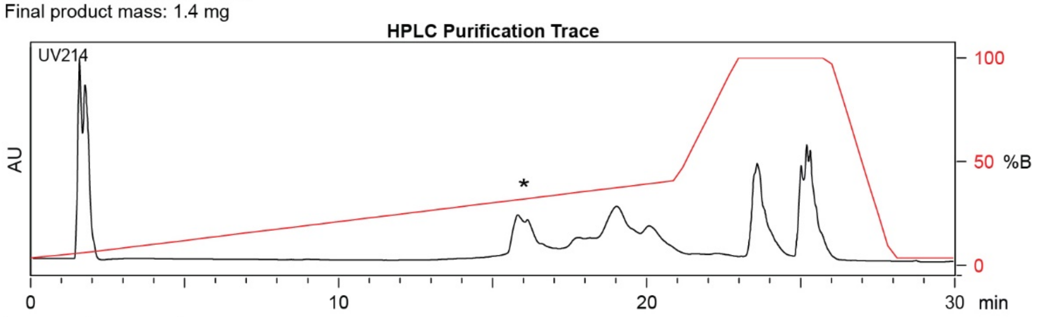

| Time (min) | Solvent B (%) |
|---|---|
| 0 | 2 |
| 2 | 5 |
| 21 | 70 |
| 23 | 100 |
| 26 | 100 |
| 29.5 | 2 |
| 30 | 2 |

Flow Rate: 13 mL / min
Phenomenex Kinetex EVO C18 Prep Column
(100 Å, 5 µm, 21.5 mm X 250 mm)

Solvent A: $H_2O$ + 0.1% (v/v) TFA
Solvent B: MeCN + 0.1% (v/v) TFA

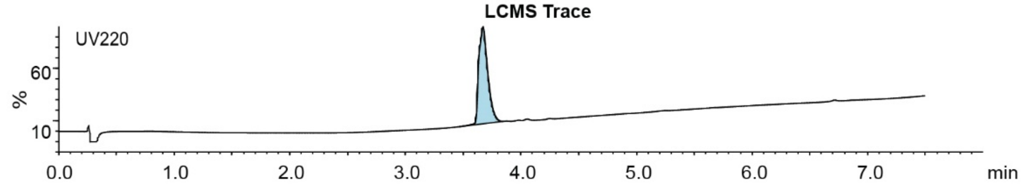

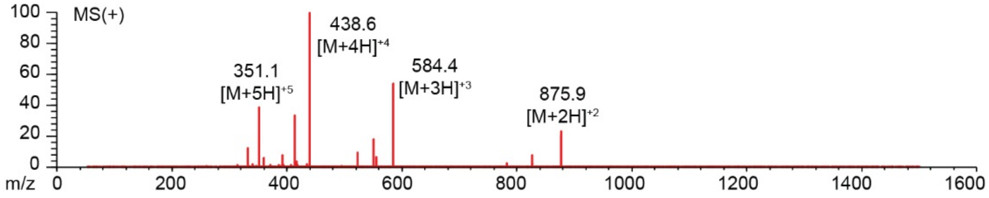

**Figure S33:** Synthesis summary of **4c**.

#### SICRFFCGGG-DFS

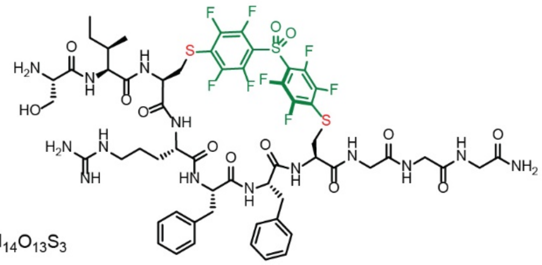

Chemical Formula: $C_{57}H_{66}F_8N_{14}O_{13}S_3$
Exact Mass: 1402.40
Molecular Weight: 1403.41

Starting material mass: 10 mg
Final product mass: 7.9 mg

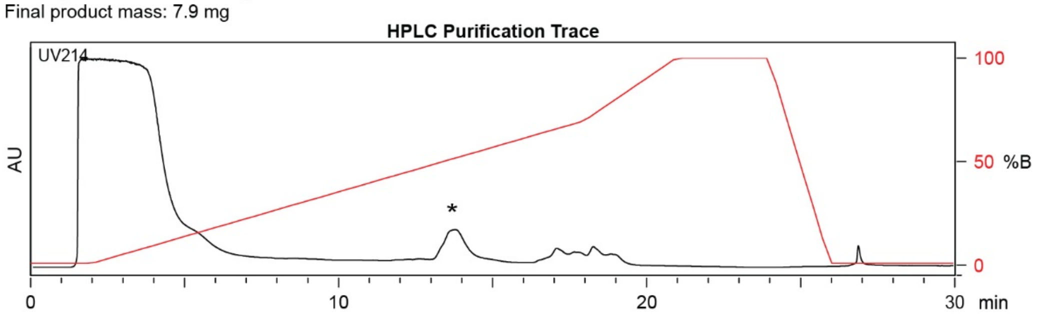

| Time (min) | Solvent B (%) |
|---|---|
| 0 | 2 |
| 2 | 5 |
| 21 | 50 |
| 23 | 100 |
| 26 | 100 |
| 29.5 | 2 |
| 30 | 2 |

Flow Rate: 13 mL / min
Phenomenex Kinetex EVO C18 Prep Column
(100 Å, 5 µm, 21.5 mm X 250 mm)

Solvent A: $H_2O$ + 0.1% (v/v) TFA
Solvent B: MeCN + 0.1% (v/v) TFA

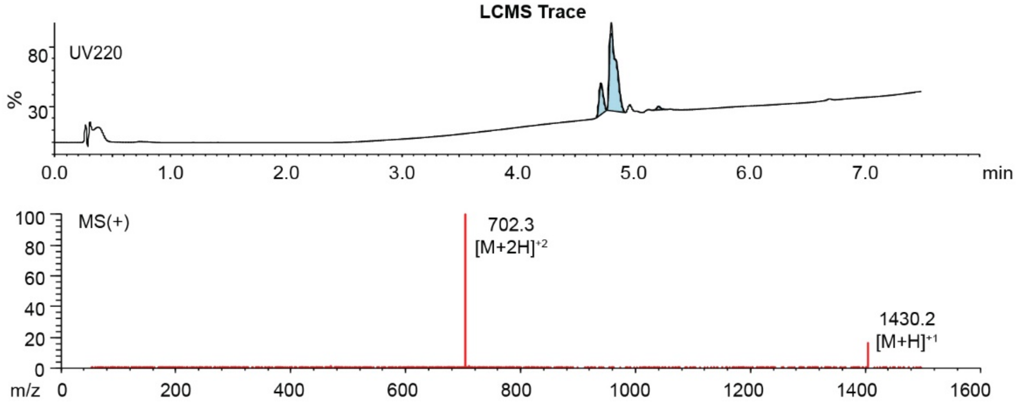

**Figure S34:** Synthesis summary of **5b**.

**SICRFFCGGG-PFS**

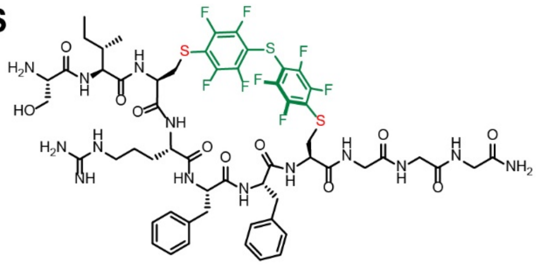

Chemical Formula: $C_{57}H_{66}F_8N_{14}O_{11}S_3$
Exact Mass: 1370.41
Molecular Weight: 1371.41

Starting material mass : 5.5 mg
Final product mass: 3.2 mg

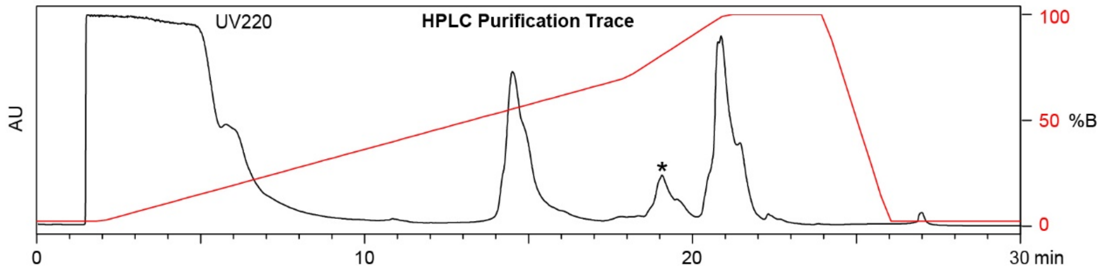

| Time (min) | Solvent B (%) |
|---|---|
| 0 | 2 |
| 2 | 2 |
| 18 | 70 |
| 21 | 100 |
| 24 | 100 |
| 26 | 0 |
| 30 | 0 |

Flow Rate: 8 mL/min
Symmetry C18 Prep Column
(100 Å, 5 µm, 10 mm X 50 mm)

Slovent A: $H_2O$ + 0.1% (v/v) TFA
Slovent B: MeCN + 0.1% (v/v) TFA

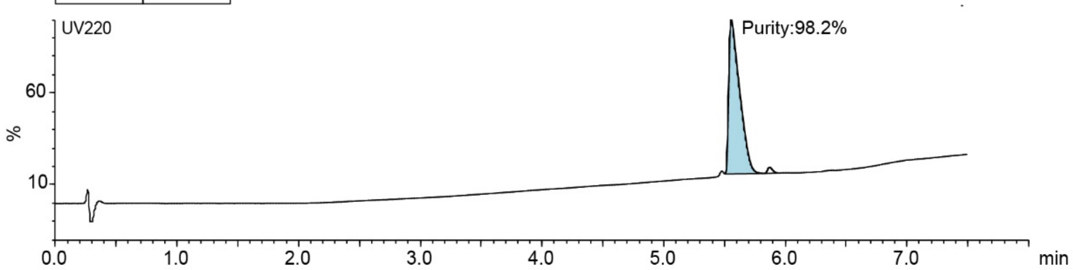

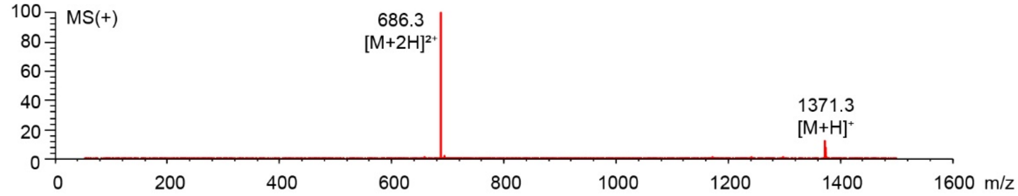

**Figure S35:** Synthesis summary of **5c**.

#### SFCPMFGGG-DFS

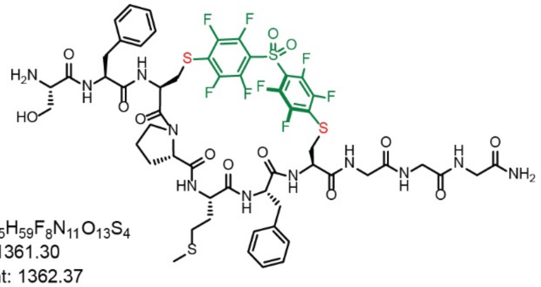

Chemical Formula: $C_{55}H_{59}F_8N_{11}O_{13}S_4$
Exact Mass: 1361.30
Molecular Weight: 1362.37

Starting material mass: 4.7 mg
Final product mass: 2.1 mg

**HPLC Purification Trace**

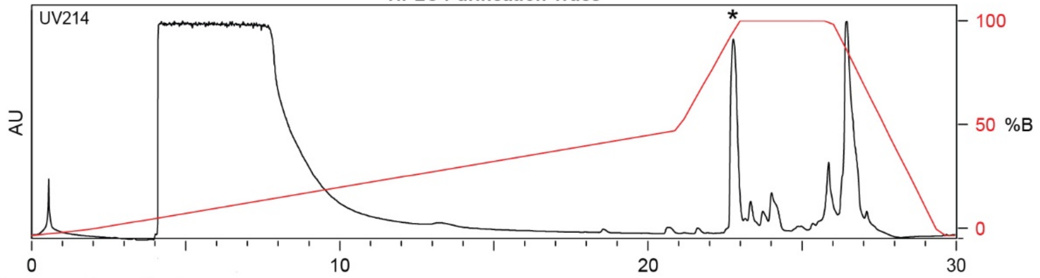

| Time (min) | Solvent B (%) |
|---|---|
| 0 | 2 |
| 2 | 5 |
| 21 | 50 |
| 25 | 100 |
| 27 | 100 |
| 28 | 2 |
| 30 | 2 |

Flow Rate: 13 mL / min
Phenomenex Kinetex EVO C18 Prep Column
(100 Å, 5 µm, 21.5 mm X 250 mm)

Slovent A: $H_2O$ + 0.1% (v/v) TFA
Slovent B: MeCN + 0.1% (v/v) TFA

**LCMS Trace**

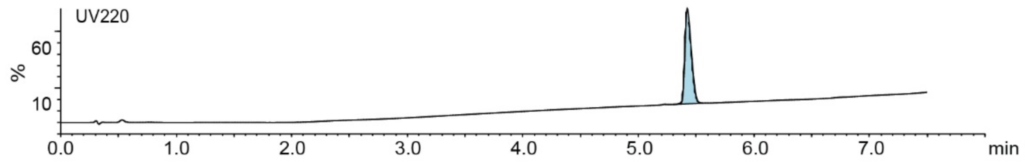

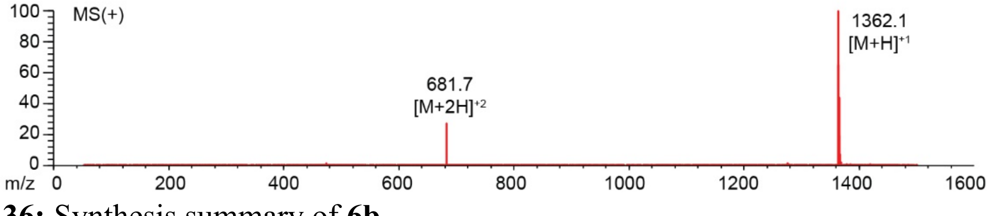

**Figure S36:** Synthesis summary of **6b**.

#### SFCPMFCGGG-PFS

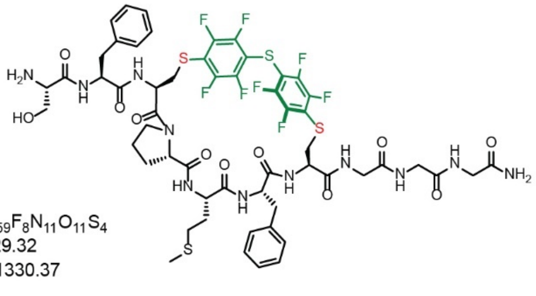

Chemical Formula: $C_{55}H_{59}F_8N_{11}O_{11}S_4$
Exact Mass: 1329.32
Molecular Weight: 1330.37

Starting material mass: 10 mg
Final product mass: 5.1 mg

**HPLC Purification Trace**

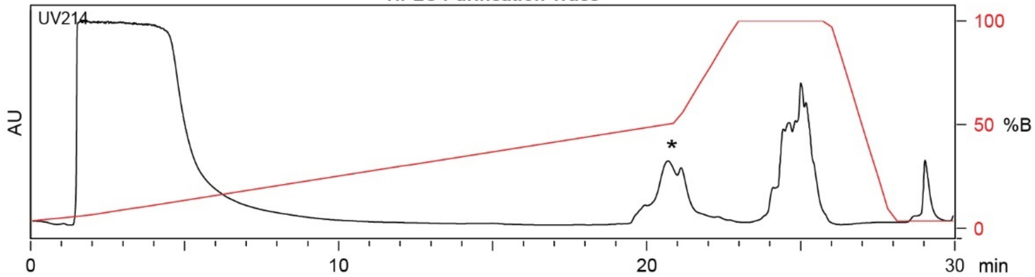

| Time (min) | Solvent B (%) |
|---|---|
| 0 | 2 |
| 2 | 5 |
| 21 | 50 |
| 23 | 100 |
| 26 | 100 |
| 29.5 | 2 |
| 30 | 2 |

Flow Rate: 13 mL / min
Phenomenex Kinetex EVO C18 Prep Column
(100 Å, 5 µm, 21.5 mm X 250 mm)

Solvent A: $H_2O$ + 0.1% (v/v) TFA
Solvent B: MeCN + 0.1% (v/v) TFA

**LCMS Trace**

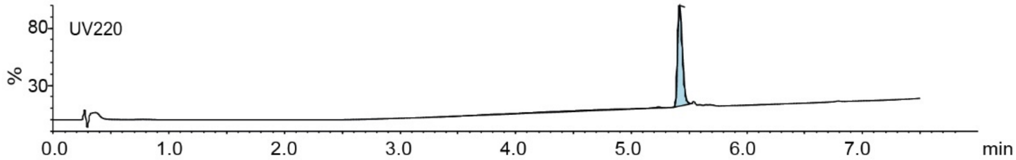

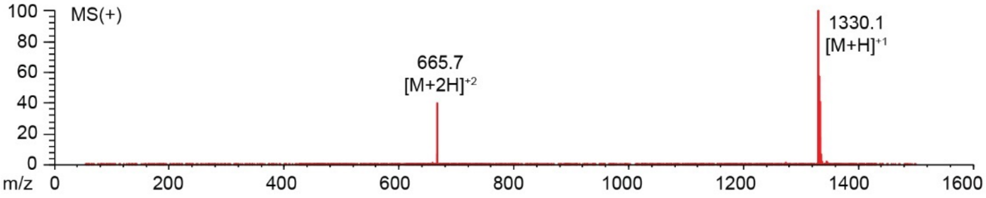

**Figure S37:** Synthesis summary of **6c**.

#### SLCKRECGGG-DFS

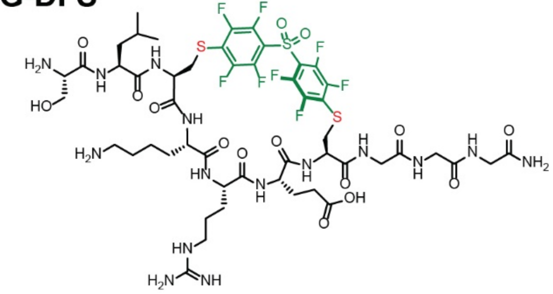

Chemical Formula: $C_{51}H_{68}F_8N_{14}O_{15}S_3$
Exact Mass: 1364.40
Molecular Weight: 1365.36

Starting material mass: 10.6 mg
Final product mass: 5.6 mg

**HPLC Purification Trace**

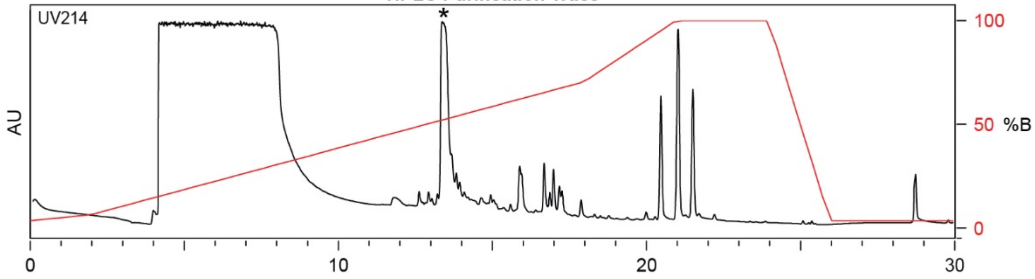

| Time (min) | Solvent B (%) |
|---|---|
| 0 | 2 |
| 2 | 5 |
| 18 | 70 |
| 21 | 100 |
| 24 | 100 |
| 26 | 2 |
| 30 | 2 |

Flow Rate: 13 mL / min
Phenomenex Kinetex EVO C18 Prep Column
(100 Å, 5 μm, 21.5 mm X 250 mm)

Solvent A: $H_2O$ + 0.1% (v/v) TFA
Solvent B: MeCN + 0.1% (v/v) TFA

**LCMS Trace**

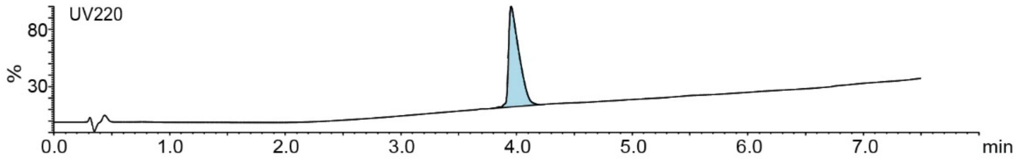

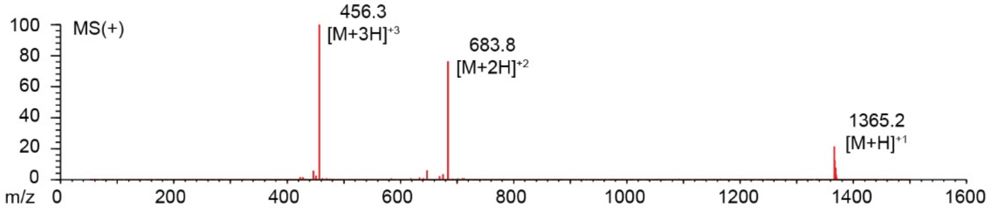

**Figure S38:** Synthesis summary of **7b**.

#### SLCKRECGGG-PFS

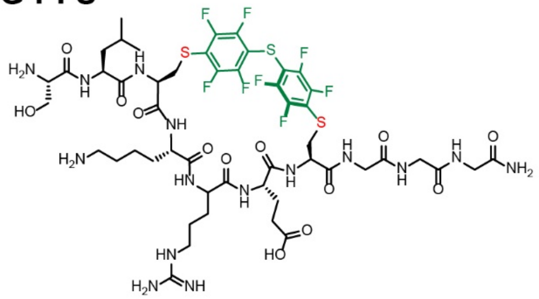

Chemical Formula: $C_{50}H_{67}F_{8}N_{15}O_{13}S_{3}$
Exact Mass: 1333.41
Molecular Weight: 1334.35

Starting material mass : 10 mg
Final product mass: 2.8 mg

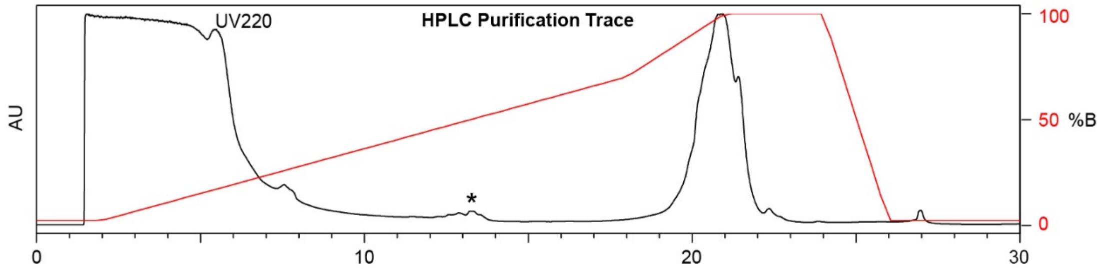

| Time (min) | Solvent B % |
|---|---|
| 0 | 2 |
| 2 | 2 |
| 18 | 70 |
| 21 | 100 |
| 24 | 100 |
| 26 | 0 |
| 30 | 0 |

Flow Rate: 8 mL/min
Symmetry C18 Prep Column
(100 Å, 5 µm, 10 mm X 50 mm)

Slovent A: $H_2O$ + 0.1% (v/v) TFA
Slovent B: MeCN + 0.1% (v/v) TFA

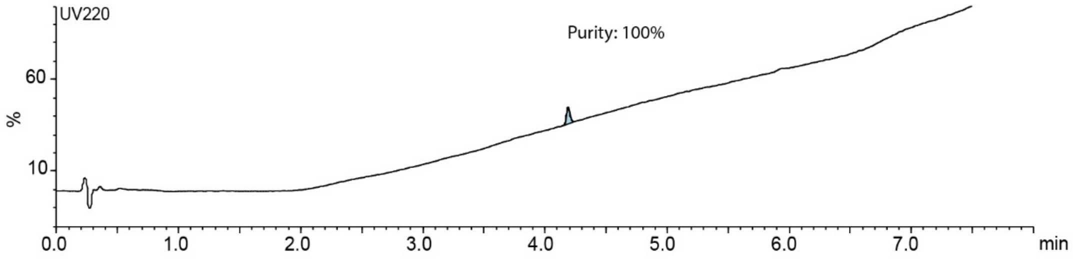

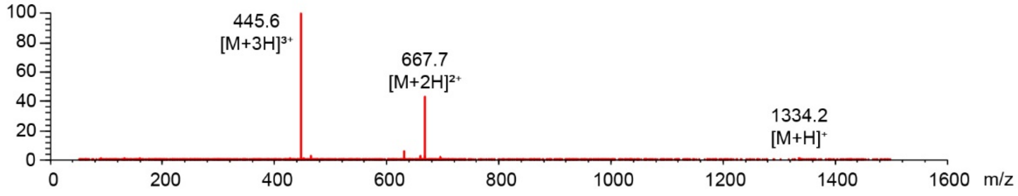

**Figure S39:** Synthesis summary of **7c**

#### STCQGECGGG-DFS

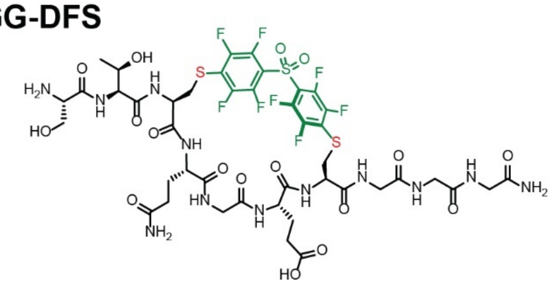

Chemical Formula: $C_{43}H_{50}F_8N_{12}O_{17}S_3$
Exact Mass: 1254.25
Molecular Weight: 1255.11

Starting material mass: 10 mg
Final product mass: 9.4 mg

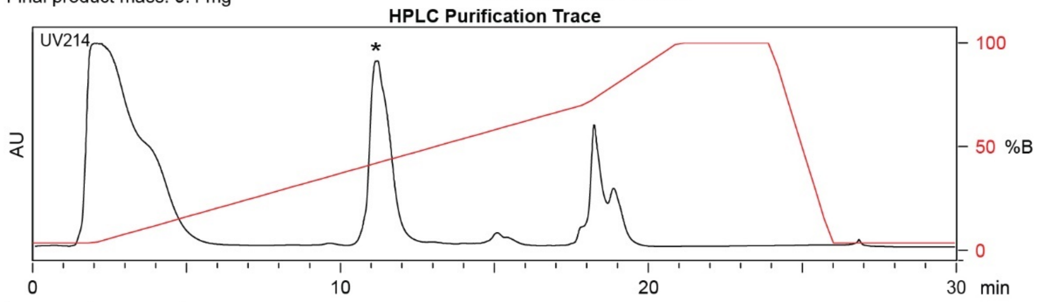

| Time (min) | Solvent B (%) |
|---|---|
| 0 | 2 |
| 2 | 5 |
| 21 | 70 |
| 23 | 100 |
| 26 | 100 |
| 29.5 | 2 |
| 30 | 2 |

Flow Rate: 13 mL / min
Phenomenex Kinetex EVO C18 Prep Column
(100 Å, 5 µm, 21.5 mm X 250 mm)

Solvent A: $H_2O$ + 0.1% (v/v) TFA
Solvent B: MeCN + 0.1% (v/v) TFA

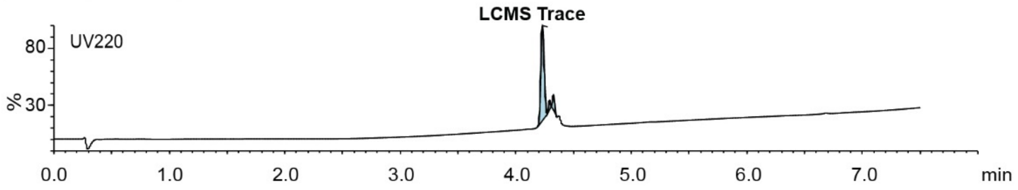

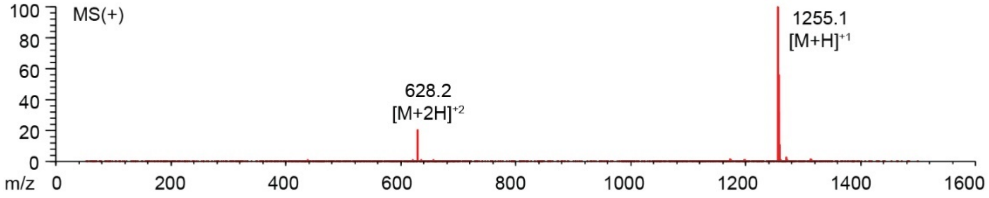

**Figure S40:** Synthesis summary of **8b**.

#### STCQGECGGG-PFS

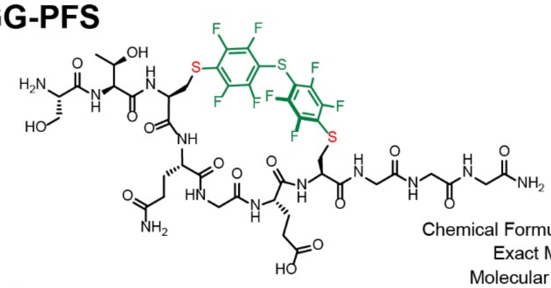

Chemical Formula: $C_{43}H_{50}F_8N_{12}O_{15}S_3$
Exact Mass: 1222.26
Molecular Weight: 1223.11

Starting material mass : 9.7 mg
Final product mass: 5.0 mg

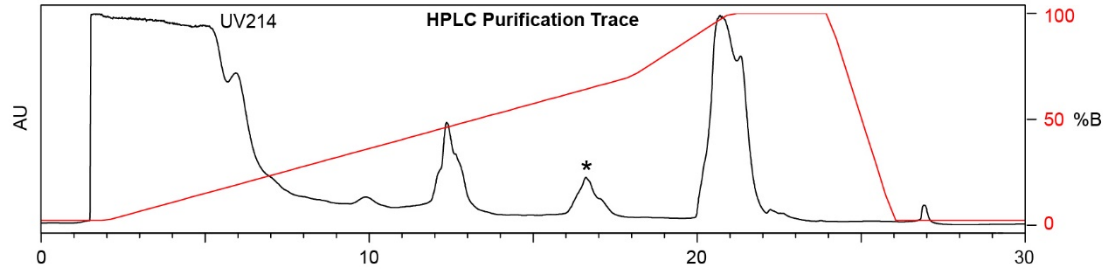

| Time (min) | Solvent B % |
|---|---|
| 0 | 2 |
| 2 | 2 |
| 18 | 70 |
| 21 | 100 |
| 24 | 100 |
| 26 | 0 |
| 30 | 0 |

Flow Rate: 8 mL/min
Symmetry C18 Prep Column
(100 Å, 5 μm, 10 mm X 50 mm)

Slovent A: $H_2O$ + 0.1% (v/v) TFA
Slovent B: MeCN + 0.1% (v/v) TFA

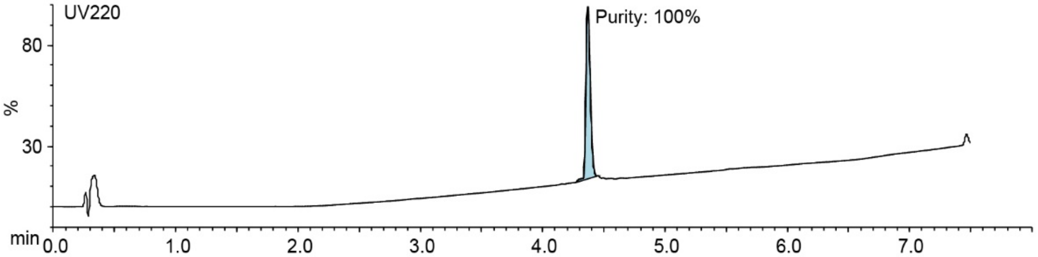

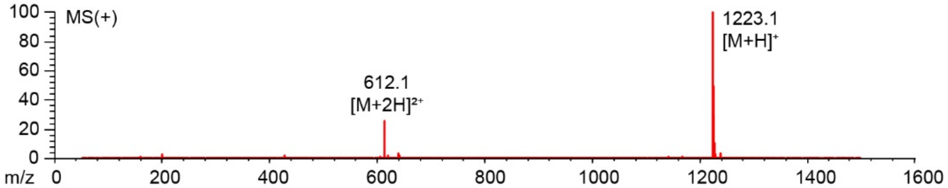

**Figure S41:** Synthesis summary of **8c**.

#### SICRFFCGGK-PFS-Bodipy

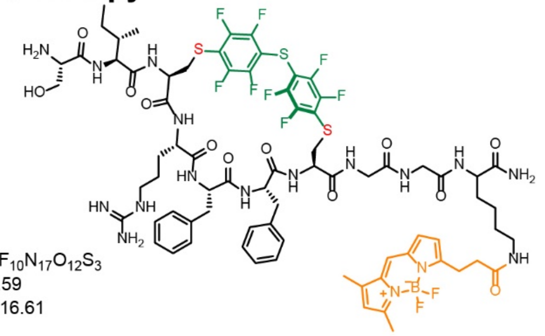

Chemical Formula: $C_{75}H_{88}BF_{10}N_{17}O_{12}S_3$
Exact Mass: 1715.59
Molecular Weight: 1716.61

Starting material mass : 1.2 mg
Final product mass: >0.1 mg

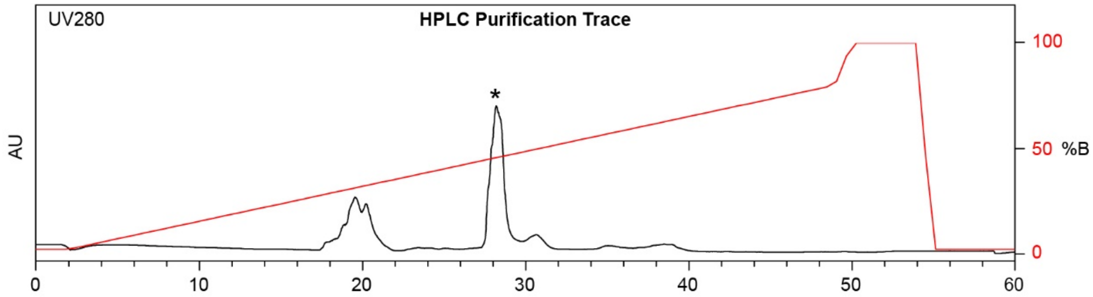

| Time (min) | Solvent B % |
|---|---|
| 0 | 2 |
| 2 | 2 |
| 49 | 80 |
| 50 | 100 |
| 54 | 100 |
| 55 | 2 |
| 60 | 2 |

Flow Rate: 8 mL/min
Symmetry C18 Prep Column
(100 Å, 5 µm, 10 mm X 50 mm)

Slovent A: $H_2O$ + 0.1% (v/v) TFA
Slovent B: MeCN + 0.1% (v/v) TFA

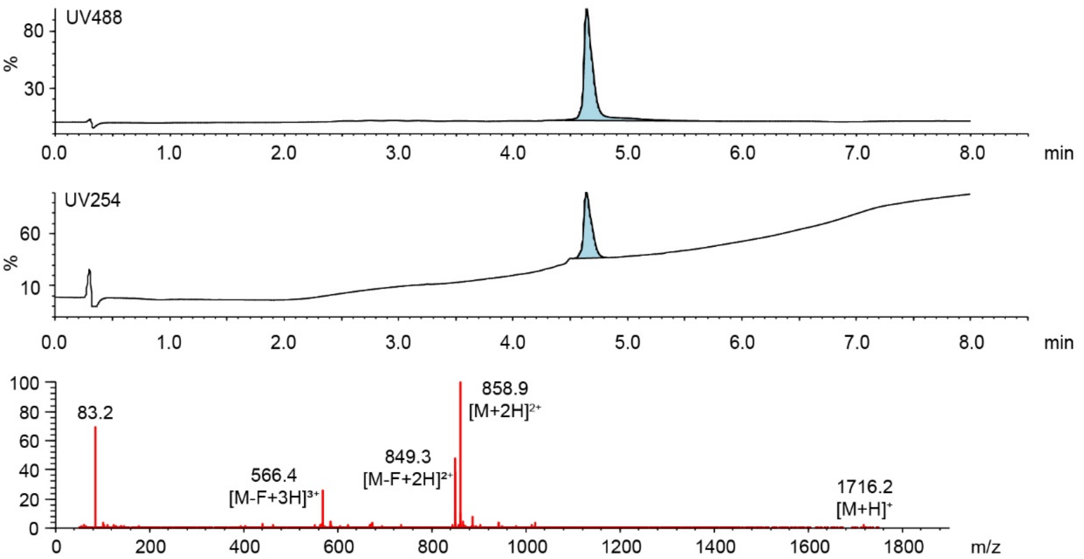

**Figure S42:** Synthesis summary of C-terminus labeled **5c**-BODIPY**.**

#### SICRFFCGGG-PFS-BODIPY

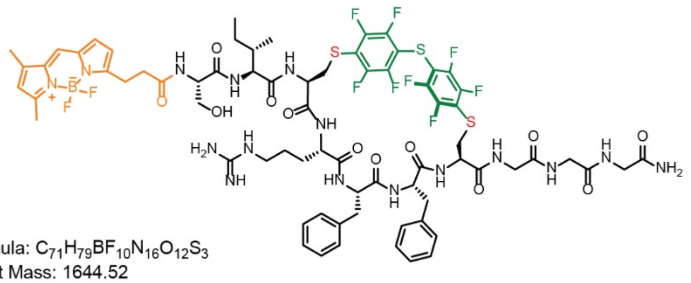

Chemical Formula: $C_{71}H_{79}BF_{10}N_{16}O_{12}S_3$
Exact Mass: 1644.52
Molecular Weight: 1645.49

Starting material mass: 1.6 mg
Final product mass: 1 mg

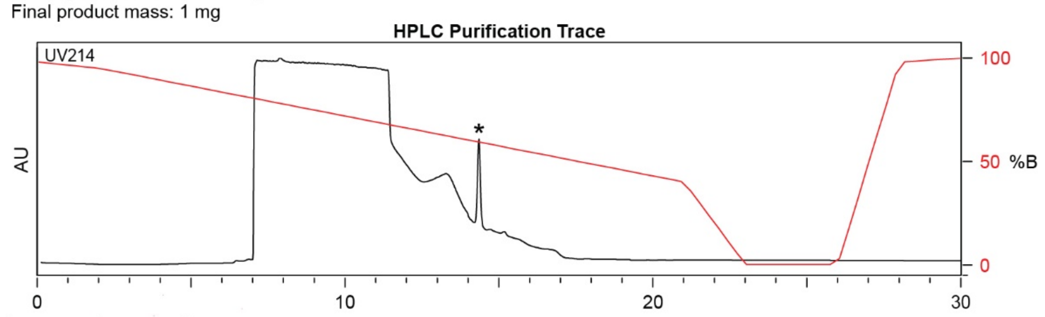

| Time (min) | Solvent B (%) |
|---|---|
| 0 | 98 |
| 2 | 95 |
| 21 | 40 |
| 23 | 0 |
| 26 | 0 |
| 28 | 98 |
| 30 | 98 |

Flow Rate: 8 mL / min
XBridge BEH Amide OBD Prep Column
(130 Å, 5 µm, 19 mm X 250 mm)

Slovent A: $H_2O$ + 0.1% (v/v) TFA
Slovent B: MeCN + 0.1% (v/v) TFA

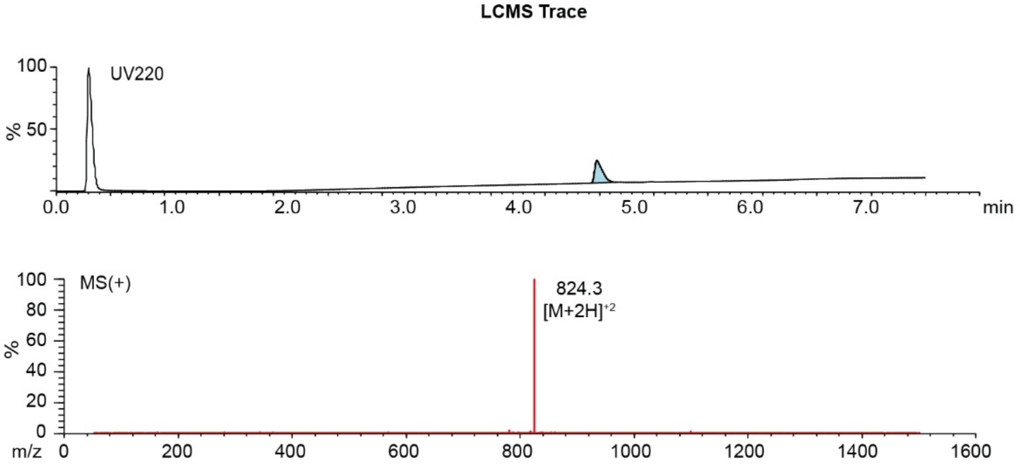

**Figure S43:** Synthesis summary of N-terminus labeled **5d-**BODIPY

### SICRFFCGGG-DFS-BODIPY

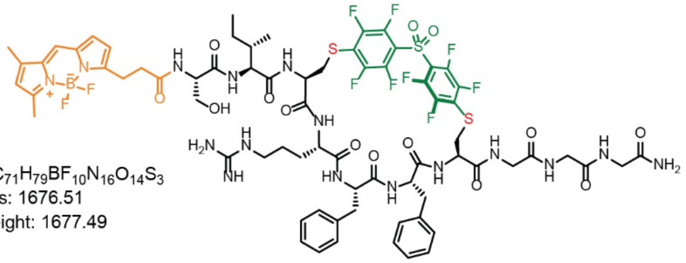

Chemical Formula: $C_{71}H_{79}BF_{10}N_{16}O_{14}S_3$
Exact Mass: 1676.51
Molecular Weight: 1677.49

Starting material mass: 2 mg
Final product mass: 1.2 mg

**HPLC Purification Trace**

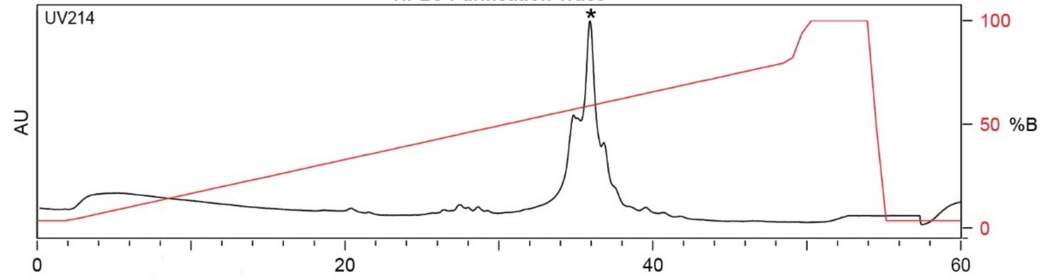

| Time (min) | Solvent B (%) |
|---|---|
| 0 | 2 |
| 2 | 2 |
| 49 | 80 |
| 50 | 100 |
| 54 | 100 |
| 55 | 2 |
| 55 | 2 |

Flow Rate: 8 mL / min
Symmetrey C18 Prep Column
(100 Å, 5 μm, 10 mm X 50 mm)

Slovent A: $H_2O$ + 0.1% (v/v) TFA
Slovent B: MeCN + 0.1% (v/v) TFA

**LCMS Trace**

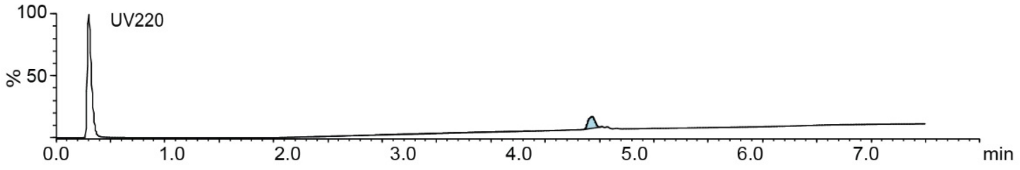

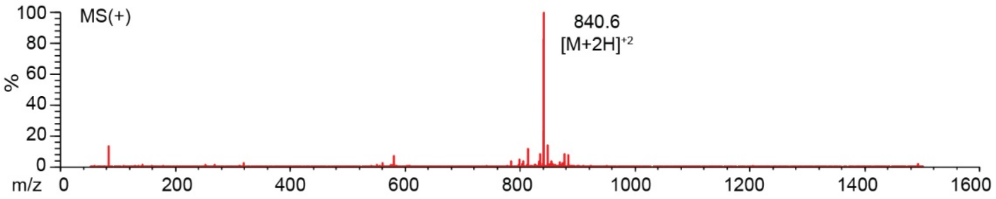

**Figure S44:** Synthesis summary of N-terminus labeled **5b-**BODIPY

#### STCQGECGGK-PFS-Bodipy

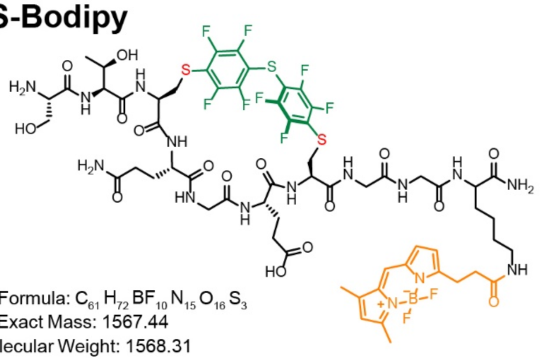

Chemical Formula: $C_{61}H_{72}BF_{10}N_{15}O_{16}S_3$
Exact Mass: 1567.44
Molecular Weight: 1568.31

Starting material mass : 1.2 mg
Final product mass: >0.1 mg

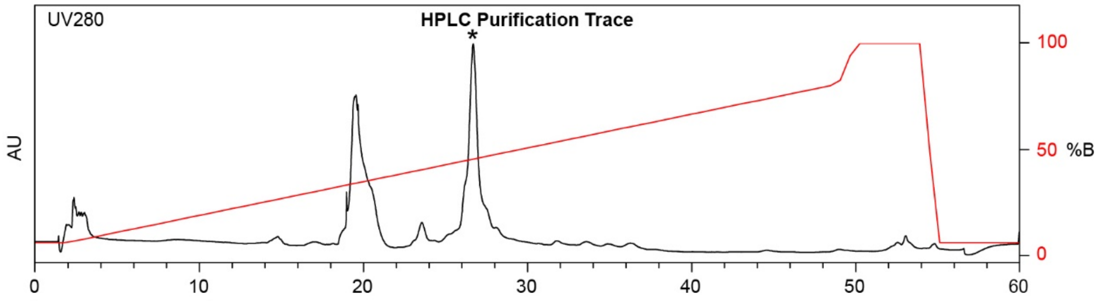

| Time (min) | Solvent B % |
|---|---|
| 0 | 2 |
| 2 | 2 |
| 49 | 80 |
| 50 | 100 |
| 54 | 100 |
| 55 | 2 |
| 60 | 2 |

Flow Rate: 8 mL/min
Symmetry C18 Prep Column
(100 Å, 5 µm, 10 mm X 50 mm)

Slovent A: $H_2O$ + 0.1% (v/v) TFA
Slovent B: MeCN + 0.1% (v/v) TFA

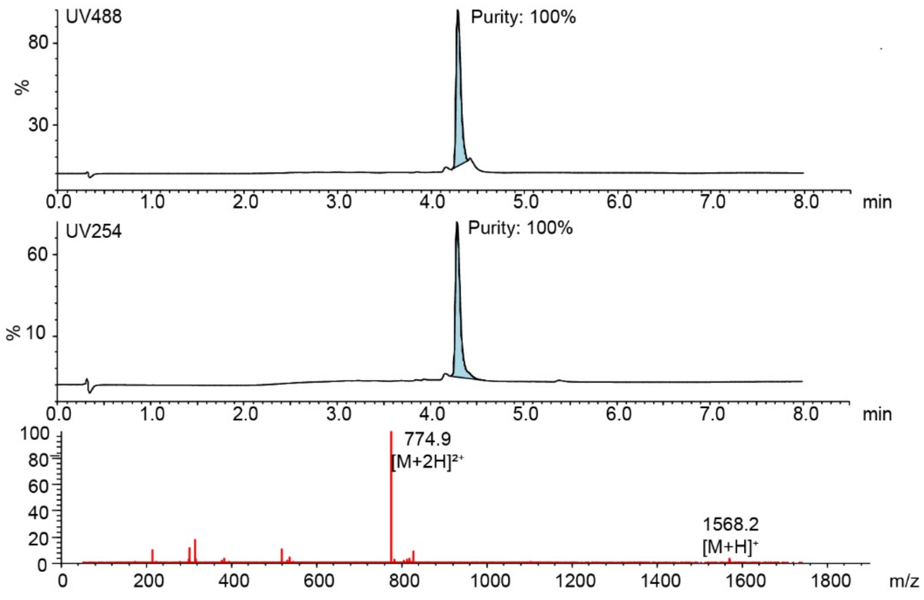

**Figure S45:** Synthesis summary of C-terminus labeled **8c-**BODIPY.

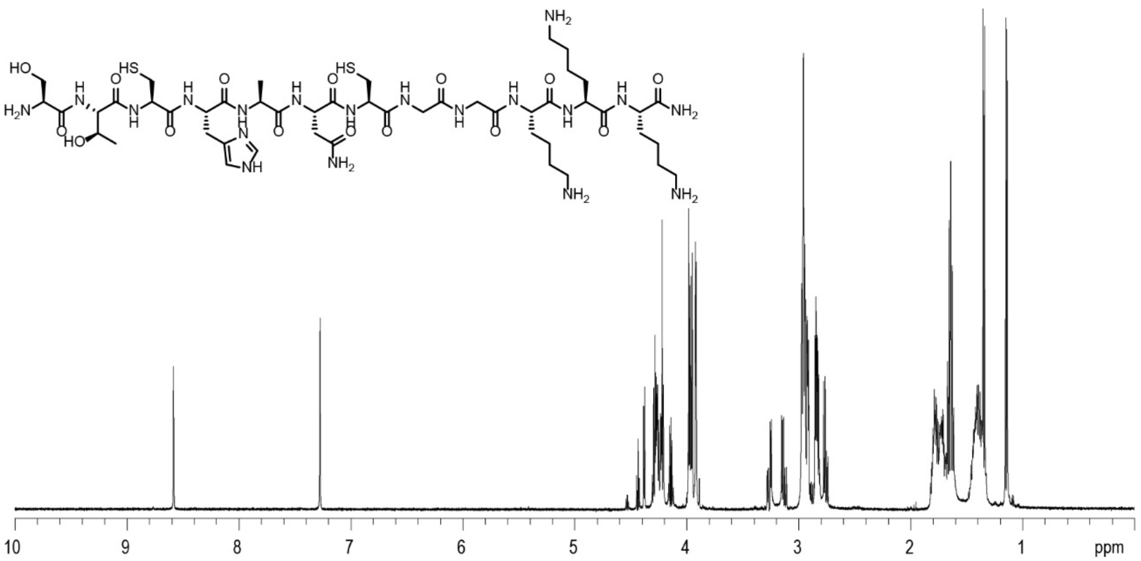

**Figure S46:** STCHANCGGKKK $^{1}$H NMR Spectra in $D_2O$, 600

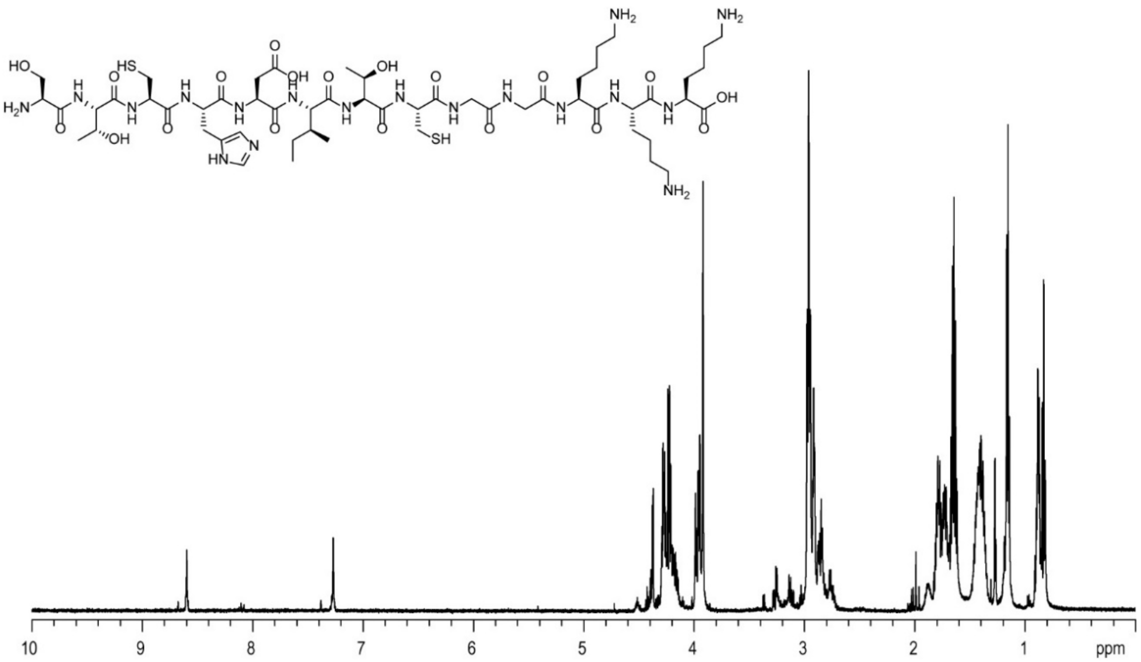

**Figure S47:** STCHDITCGGKKK $^{1}$H NMR Spectra in $D_2O$, 600

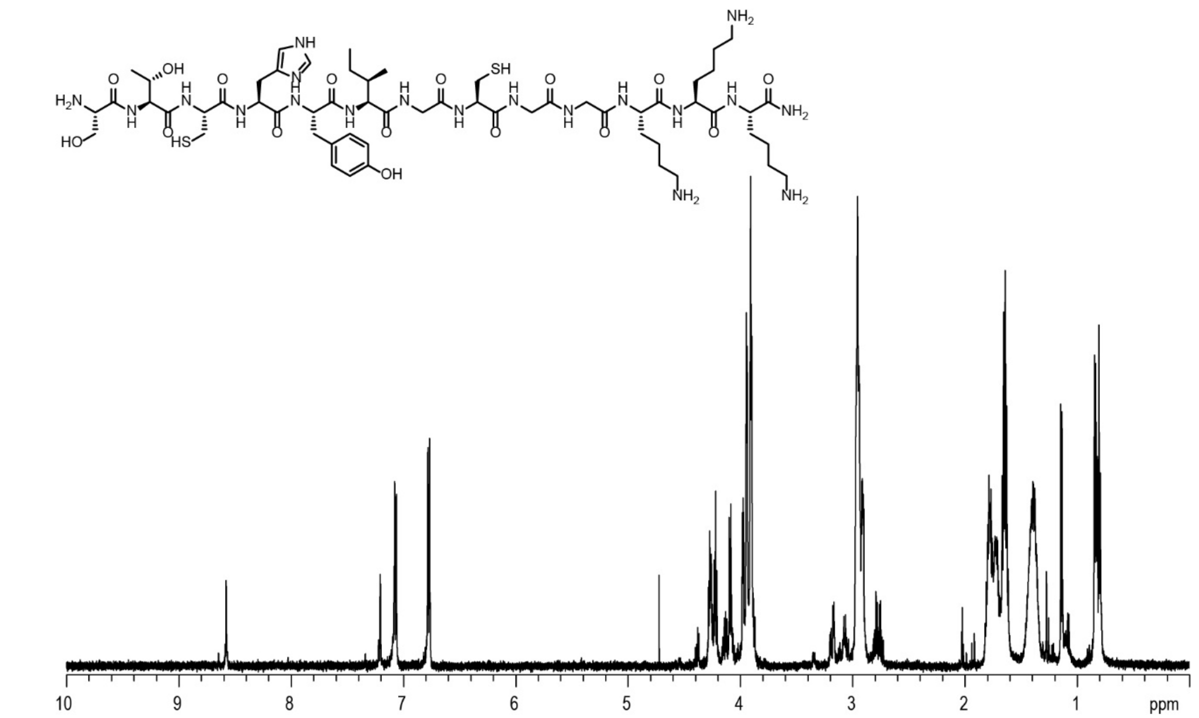

**Figure S48:** STCHYIGCGGKKK $^1$H NMR Spectra in $D_2O$, 600

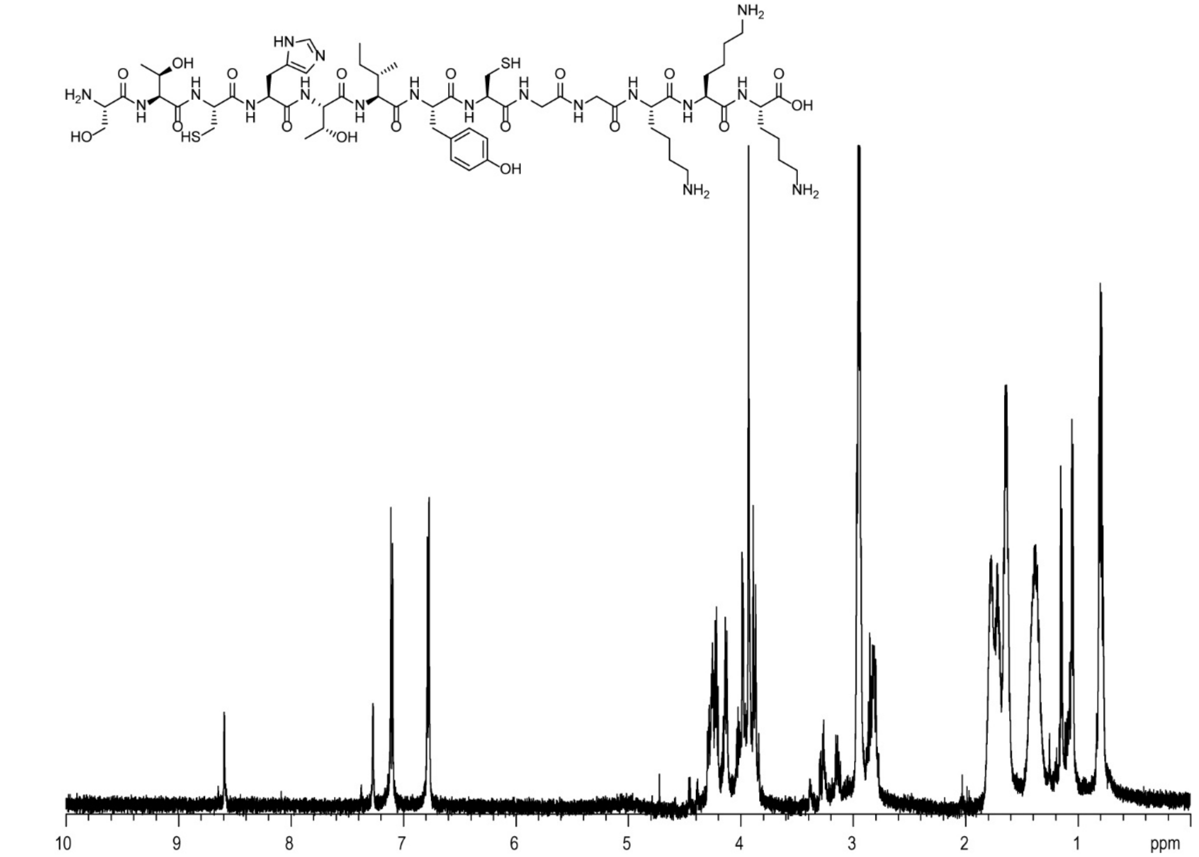

**Figure S49:** STCHTIYCGGKKK $^1$H NMR Spectra in $D_2O$, 600

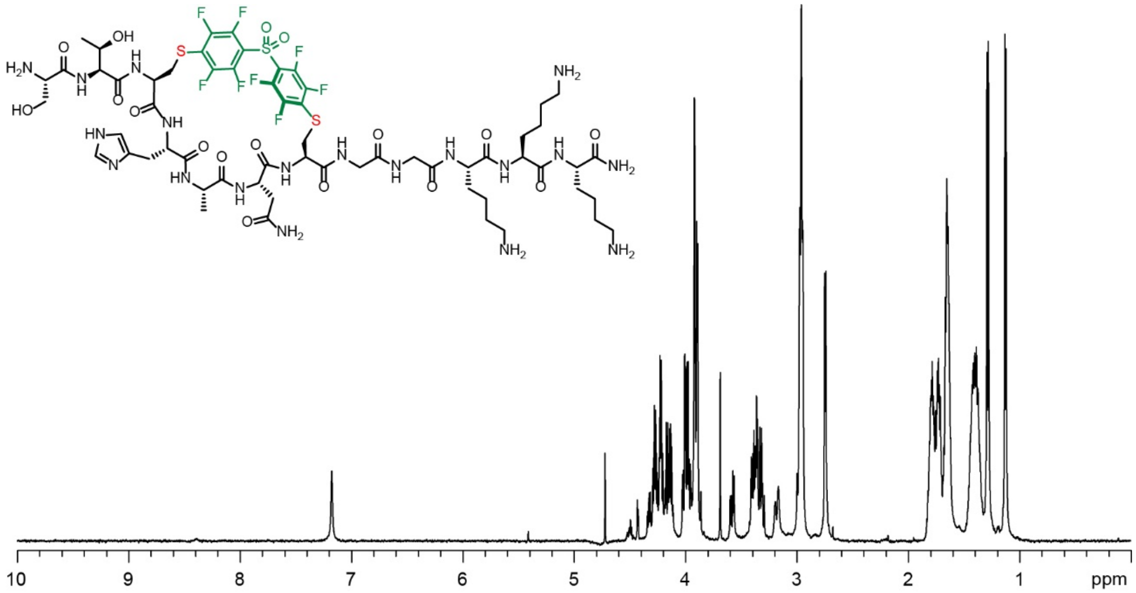

**Figure S50: DFS**-STCHANCGGKKK $^{1}$H NMR Spectra in $D_2O$, 600

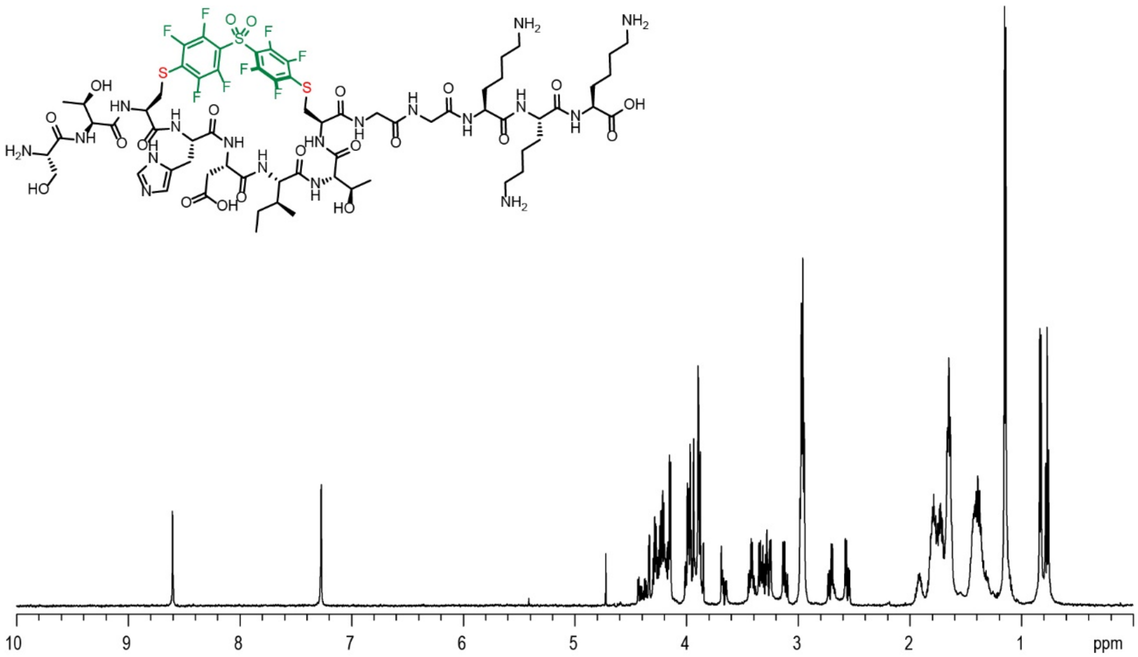

**Figure S51: DFS**-STCHDITCGGKKK $^{1}$H NMR Spectra in $D_2O$, 600

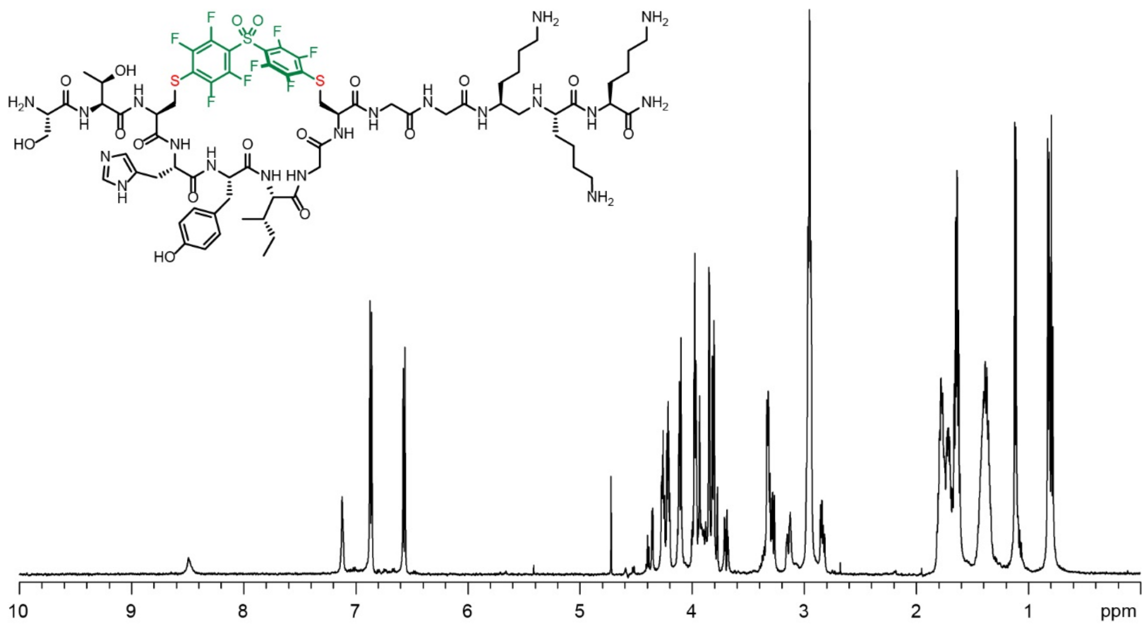

**Figure S52:** **DFS**-STCHYIGCGGKKK $^{1}$H NMR Spectra in $D_2O$, 600

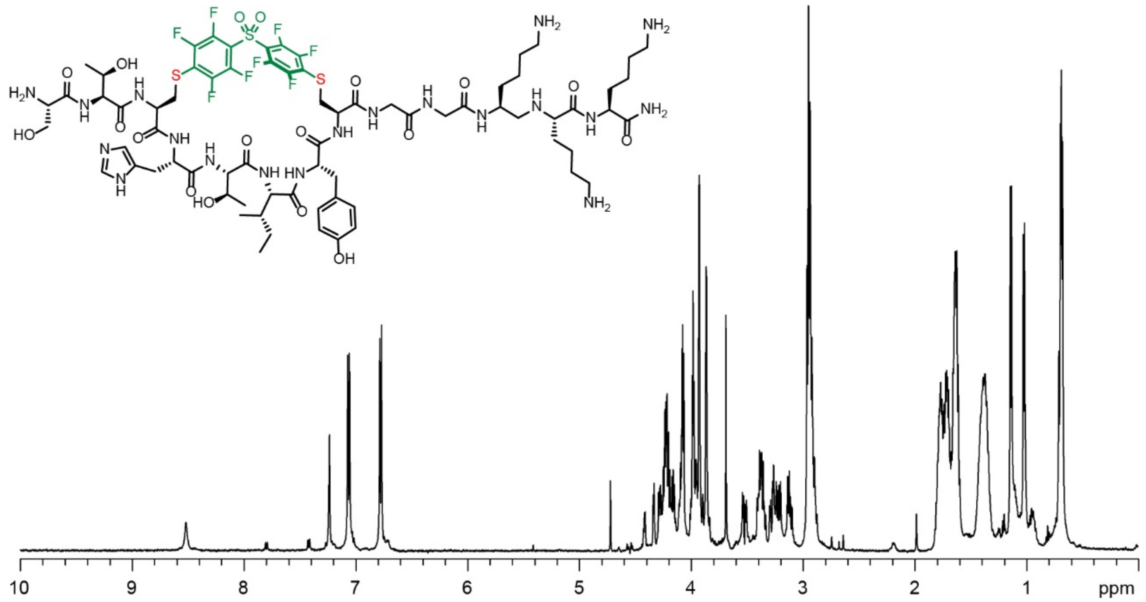

**Figure S53:** **DFS**-STCHTIYCGGKKK $^{1}$H NMR Spectra in $D_2O$, 600

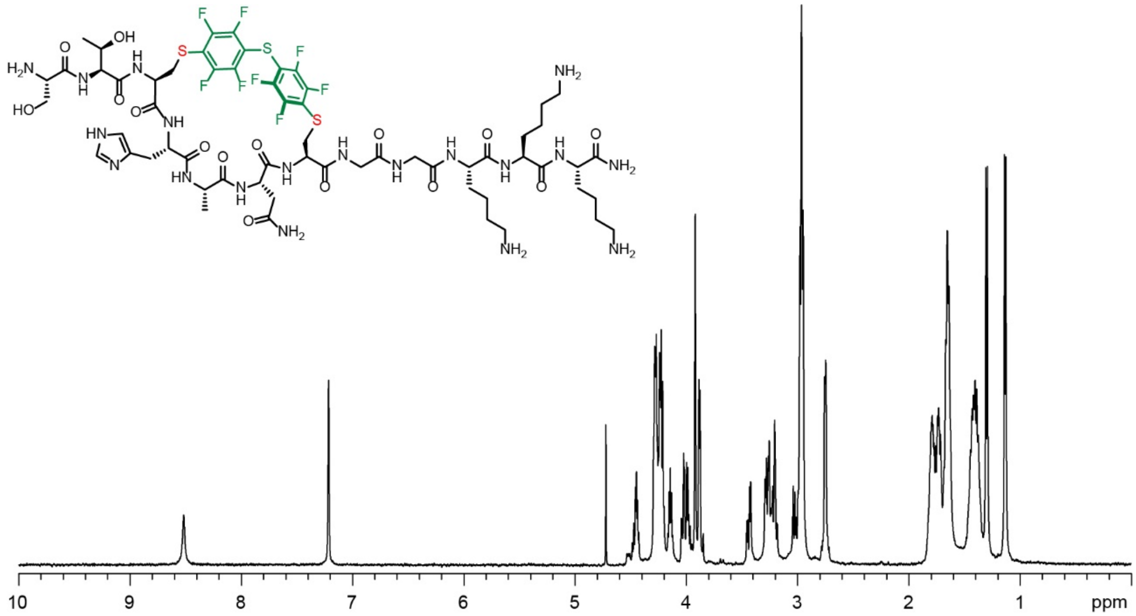

**Figure S54: PFS**-STCHANCGGKKK $^{1}H$ NMR Spectra in $D_2O$, 600

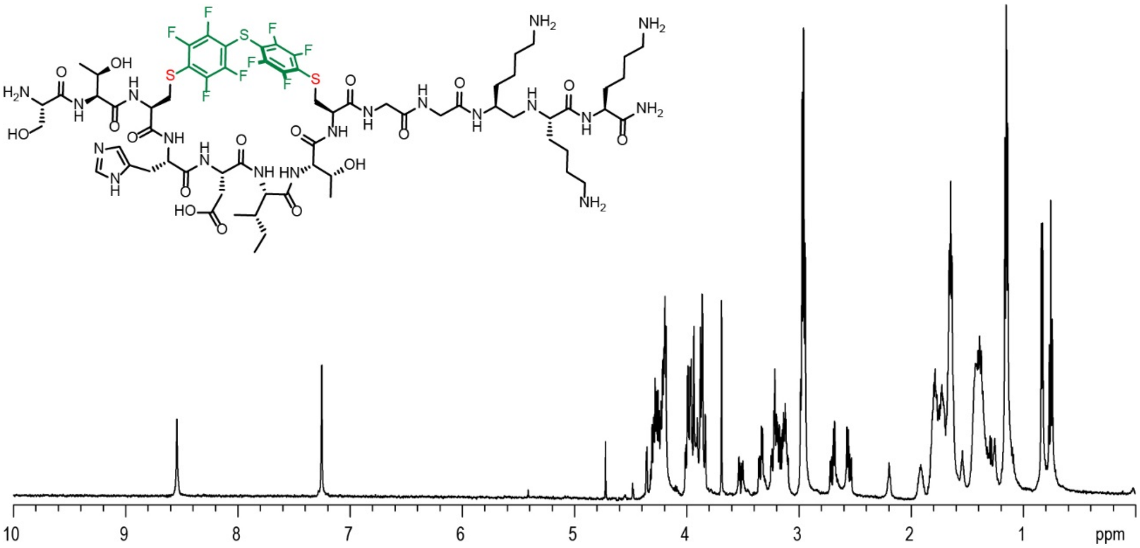

**Figure S55: PFS**-STCHDITCGGKKK $^{1}H$ NMR Spectra in $D_2O$, 600

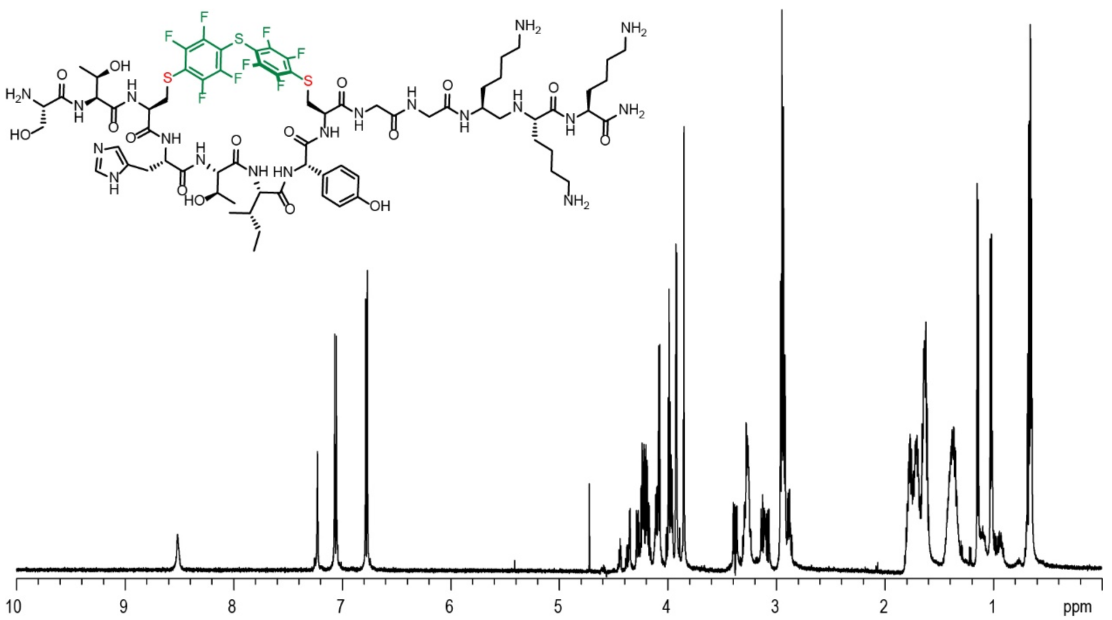

**Figure S60: PFS**-STCHTIYCGGKKK $^{1}H$ NMR Spectra in $D_2O$, 600

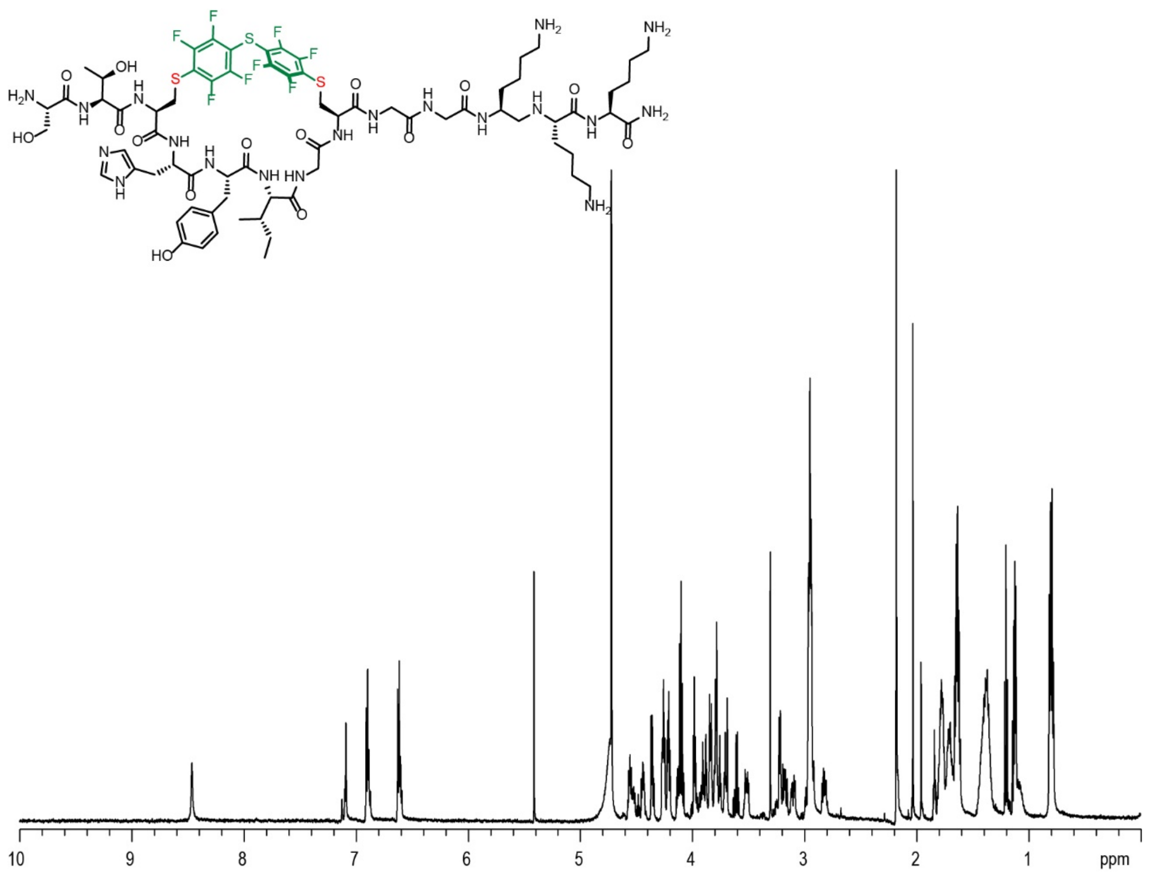

**Figure S57: PFS** stapled STCHYIGCGGKKK $^{1}H$ NMR Spectra in $D_2O$, 600

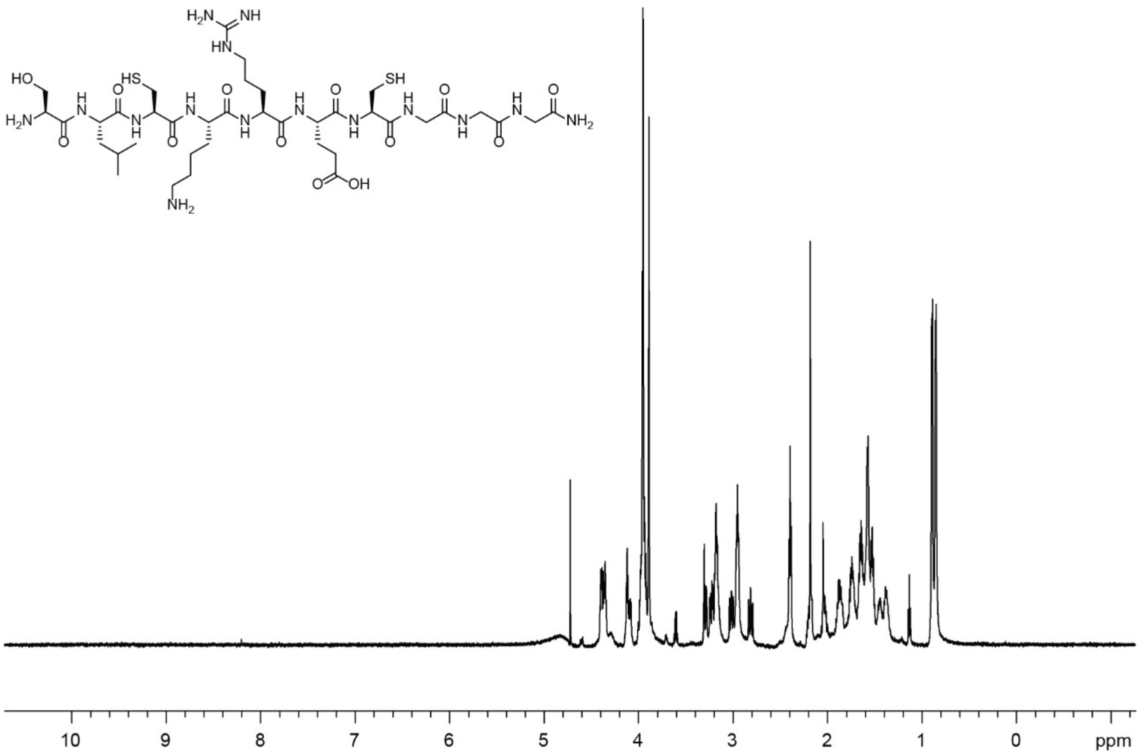

**Figure S58:** SLCKRECGGG $^1H$ NMR Spectra in $D_2O$, 600

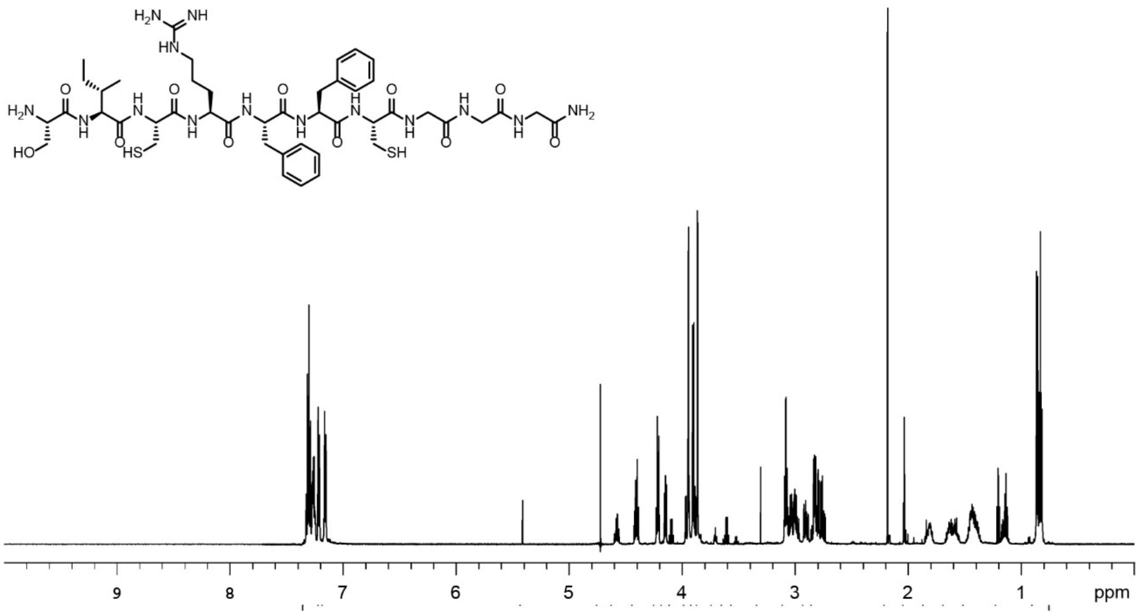

**Figure S59:** SICRFFCGGG $^1H$ NMR Spectra in $D_2O$, 600

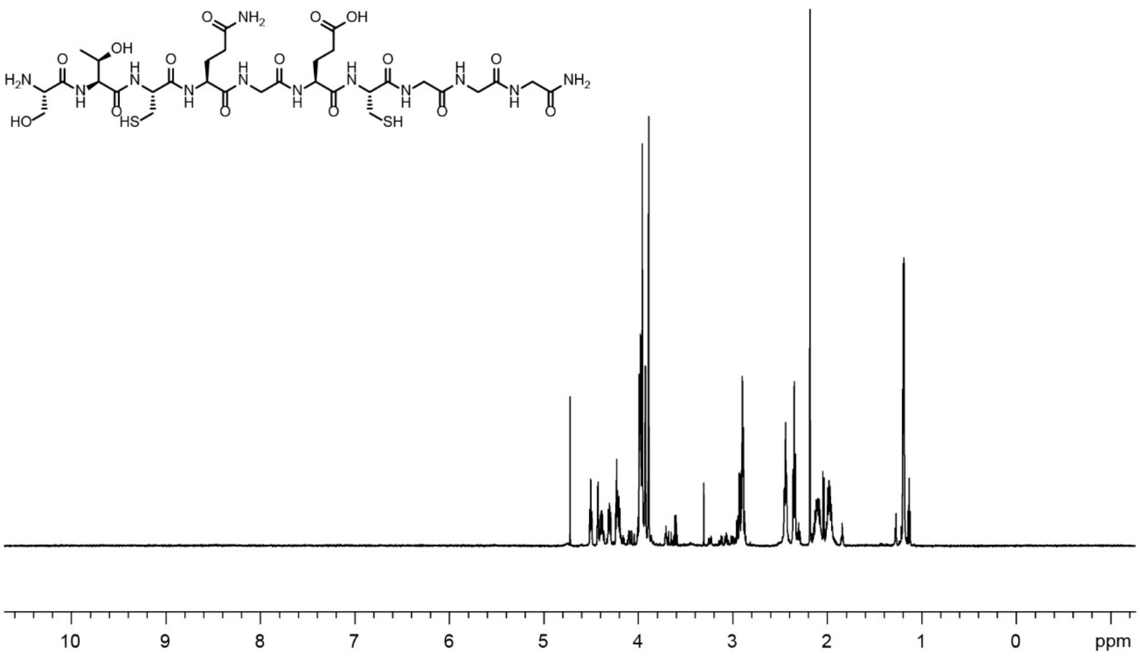

**Figure S60:** STCQGECGGG $^{1}H$ NMR Spectra in $D_2O$, 600

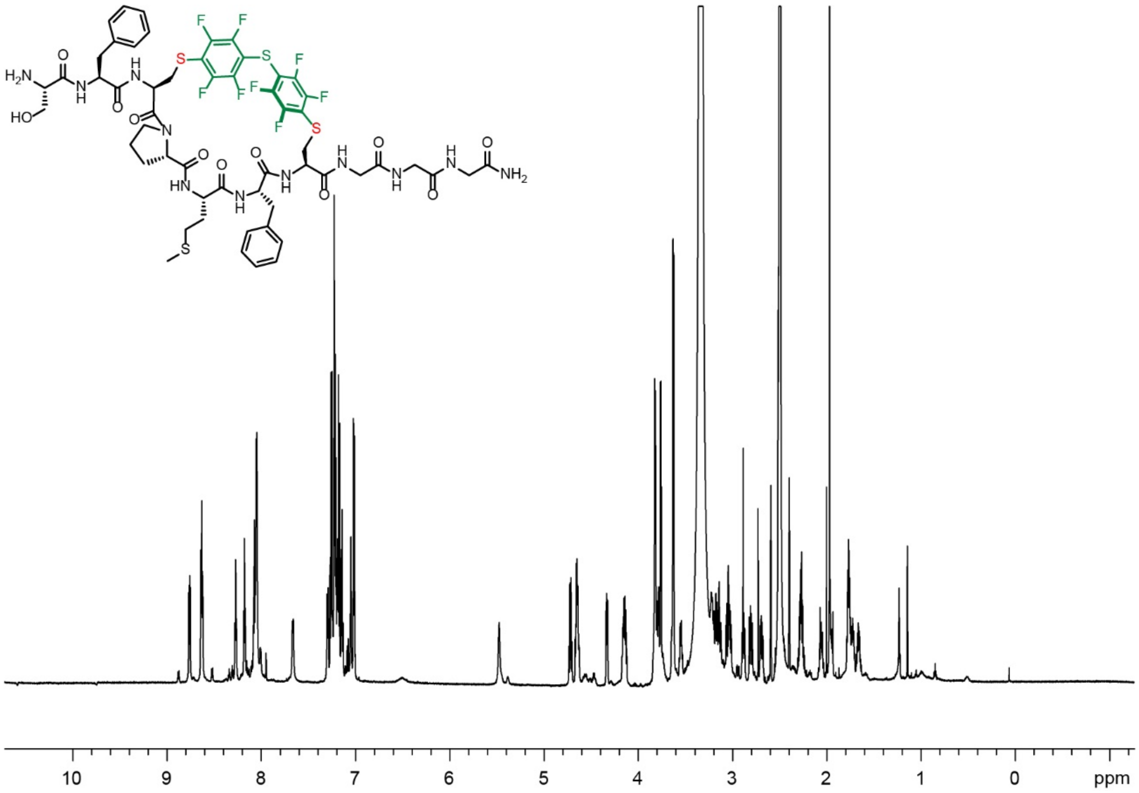

**Figure S61: PFS** stapled SFCPMFCGGG $^{1}H$ NMR Spectra in DMSO, 700

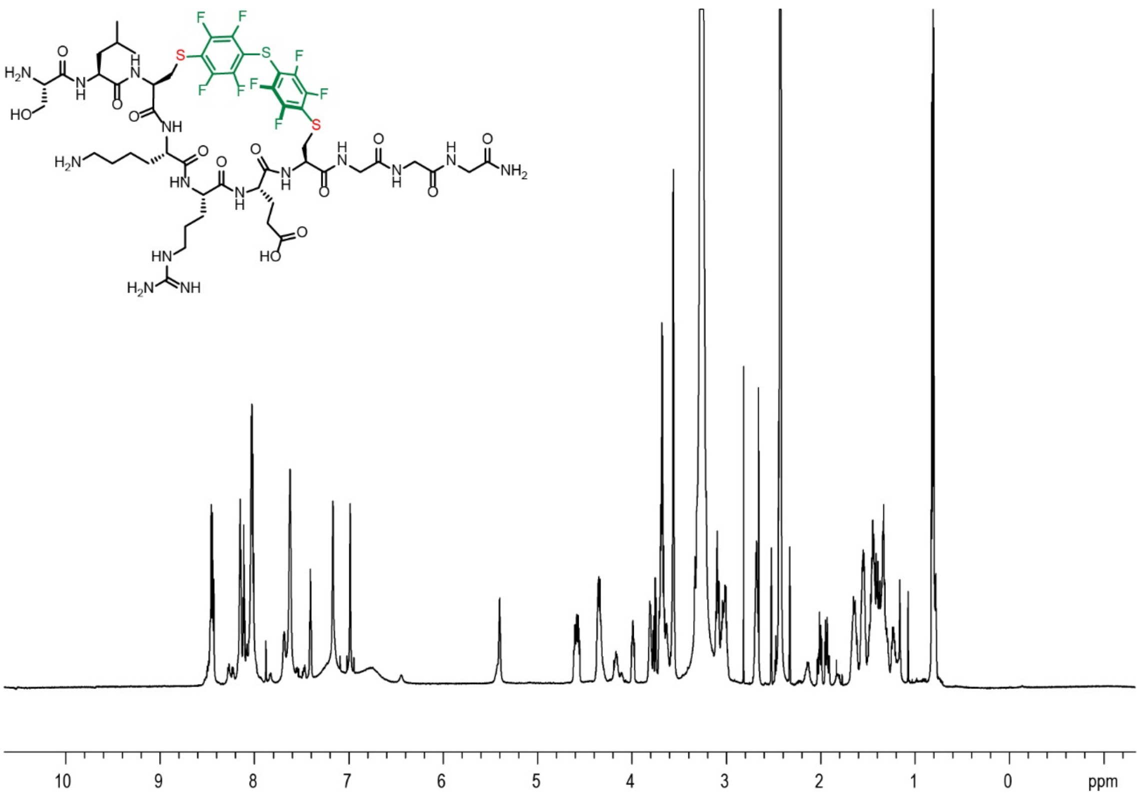

**Figure S62: PFS** stapled SLCKRECGGG $^{1}$H NMR Spectra in DMSO, 700

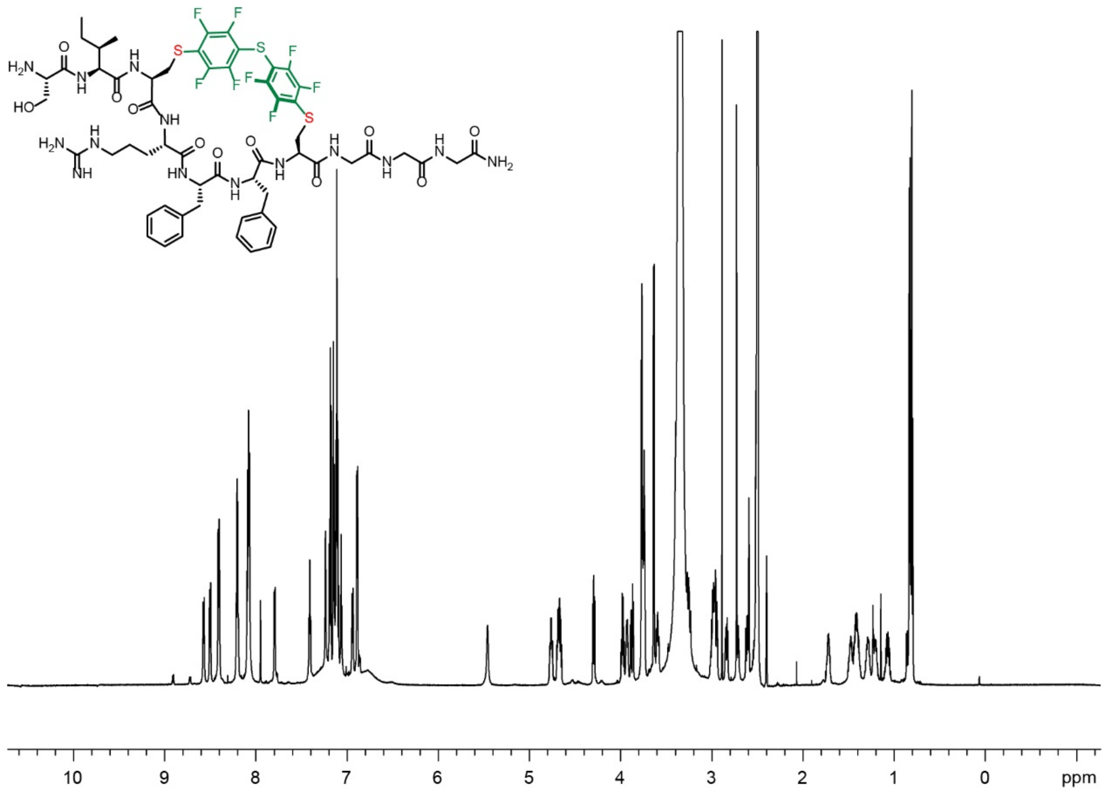

**Figure S63: PFS** stapled SICRFFCGGG $^{1}$H NMR Spectra in DMSO, 700

**21.** References

1. He, B.; Tjhung, K. F.; Bennett, N. J.; Chou, Y.; Rau, A.; Huang, J.; Derda, R., Compositional Bias in Naive and Chemically-modified Phage-Displayed Libraries uncovered by Paired-end Deep Sequencing. *Sci. Rep.* **2018,** *8* (1), 1214.

2. Sojitra, M.; Sarkar, S.; Maghera, J.; Rodrigues, E.; Carpenter, E. J.; Seth, S.; Ferrer Vinals, D.; Bennett, N. J.; Reddy, R.; Khalil, A.; Xue, X.; Bell, M. R.; Zheng, R. B.; Zhang, P.; Nycholat, C.; Bailey, J. J.; Ling, C.-C.; Lowary, T. L.; Paulson, J. C.; Macauley, M. S.; Derda, R., Genetically encoded multivalent liquid glycan array displayed on M13 bacteriophage. *Nat. Chem. Biol.* **2021,** *17* (7), 806-816.

3. Tjhung, K. F.; Kitov, P. I.; Ng, S.; Kitova, E. N.; Deng, L.; Klassen, J. S.; Derda, R., Silent Encoding of Chemical Post-Translational Modifications in Phage-Displayed Libraries. *J. Am. Chem. Soc.* **2016,** *138* (1), 32-35.

4. Bhattacharya, A. A.; Grune, T.; Curry, S., Crystallographic analysis reveals common modes of binding of medium and long-chain fatty acids to human serum albumin. *J. Mol. Biol.* **2000,** *303* (5), 721-32.

5. Trott, O.; Olson, A. J., AutoDock Vina: improving the speed and accuracy of docking with a new scoring function, efficient optimization, and multithreading. *J. Comput. Chem.* **2010,** *31* (2), 455-61.

6. Andersen, J. T.; Dalhus, B.; Cameron, J.; Daba, M. B.; Plumridge, A.; Evans, L.; Brennan, S. O.; Gunnarsen, K. S.; Bjørås, M.; Sleep, D.; Sandlie, I., Structure-based mutagenesis reveals the albumin-binding site of the neonatal Fc receptor. *Nat. Comm.* **2012,** *3* (1), 610.

7. Staquicini, F. I.; Ozawa, M. G.; Moya, C. A.; Driessen, W. H.; Barbu, E. M.; Nishimori, H.; Soghomonyan, S.; Flores, L. G., 2nd; Liang, X.; Paolillo, V.; Alauddin, M. M.; Basilion, J. P.; Furnari, F. B.; Bogler, O.; Lang, F. F.; Aldape, K. D.; Fuller, G. N.; Hook, M.; Gelovani, J. G.;

Sidman, R. L.; Cavenee, W. K.; Pasqualini, R.; Arap, W., Systemic combinatorial peptide selection yields a non-canonical iron-mimicry mechanism for targeting tumors in a mouse model of human glioblastoma. *J. Clin. Invest.* **2011,** *121* (1), 161-73.
